# Theory of interaction between untuned modulatory inputs and tuned sensory inputs

**DOI:** 10.1101/2025.04.28.651100

**Authors:** Tuan Nguyen, Kenneth D. Miller, Agostina Palmigiano

## Abstract

How does the brain integrate sensory inputs with non-feature-tuned signals, such as those arising from behavioral state changes or neuromodulation? Here, we show that the dynamics of disordered E/I networks with structured, feature-dependent connectivity can be well characterized by an effective model describing interactions between the responses of cells who prefer the current sensory stimulus (“matched” cells) and the responses of cells firing at the baseline. This effective network exhibits strong feedback from the baseline onto the matched responses but weak reverse projections. Thus, an untuned stimulus not only directly drives matched cells, but also indirectly drives them via modulation of the baseline. We demonstrate through a linear response analysis that the baseline effect on the matched response is suppressive if the network is strongly coupled and feedback-inhibition dominated. In particular, in this regime, feature-dependent networks produce “rate reshuffling”, wherein untuned optogenetic excitation yields large changes in the individual responses of matched cells without significantly changing their overall firing rate distribution, as the optogenetically-induced baseline response suppresses the matched response. Finally, if multiple sensory stimuli are presented, yielding sublinear response summation (“normalization”), the influence of the baseline on the matched responses is weakened. Thus, an untuned (*e*.*g*., optogenetic) stimulus is less suppressive to multiple stimuli than to a single stimulus, making normalization effectively weaker in the presence of an untuned stimulus. Our framework provides the first theory of the interaction of untuned modulatory and tuned sensory inputs, reconciles prior experiments, and provides testable predictions about tuned-untuned interactions in cortical processing.

**Significance Statement:** Sensory cortex receives both tuned sensory inputs, and untuned signals *e*.*g*. from global stimulus changes, behavioral state changes, or neuromodulation. We lack a theory of how these inputs are integrated. We demonstrate reduction of circuit models to a model of interactions between baseline and sensory-stimulus-matched responses, and develop an exact linear response analysis of activity perturbation by untuned inputs. We show baseline-to-matched coupling is strong, while matched-to-baseline is weak, and the former is suppressive given sufficiently strong and inhibition-dominated connectivity. We extend the theory to multiple tuned stimuli. The theory offers a mechanistic explanation of previous surprising observations (“rate reshuffling”), yields new predictions, and provides a general framework for understanding the impact of modulatory influences on sensory processing.

## Introduction

How does the cortex integrate tuned sensory inputs with untuned modulatory inputs? Sensory cortex receives sensory (*e*.*g*., visual) inputs that are tuned for features of the sensory stimulus, but also receives many untuned modulatory inputs, *i*.*e*. inputs that are received by cells without regard for their sensory tuning. These modulatory inputs arise from such factors as behavioral state changes, *e*.*g*. arousal or locomotion, which drive release of neuromodulators such as norepinephrine and acetylcholine (1–4); global stimulus changes, *e*.*g*. a visual luminance change (5); spatial attention (6); reinforcement signals (7); or artificial stimuli such as one-photon optogenetic stimulation. While many forms of neural response modulation show both additive (untuned) and multiplicative (tuned) components (8), multiplicative response modulation can result from additive modulatory input (9).

Here we develop a general theory of how tuned and untuned inputs are integrated by recurrent networks with feature-dependent connection probabilities composed of excitatory (E) and inhibitory (I) neurons. For simplicity, we neglect space, focusing only on variation in feature preferences of neurons at a single location. We show that, if the dependence of the firing rate moments on the feature can be approximated by a baseline-plus-Gaussian mixture, then the mean-field equations describing the feature-dependent mean and autocorrelation functions of the E and I populations reduce to those of an effective model of interactions between the population of neurons tuned to the current stimulus features (“sensorystimulus-matched” population or “matched” for short) and the population firing at the baseline level (“baseline” population).

Importantly, these effective networks exhibit strong connections from the baseline onto the matched population, but asymmetrically weak projections in the reverse direction. Through a linear response analysis we show that these connections are primarily inhibitory when the network is strongly coupled and feedback inhibition dominated. In conclusion, we show that an untuned modulatory stimulus can both directly drive the matched cells and indirectly suppress them, with the balance of these effects determining whether their mean response to the untuned stimulus is positive, negative, or zero.

We apply our theory to investigate the effect of artificial manipulations, such as non-specific optogenetic stimulation. In particular, recent work investigated how visually driven responses in macaque primary visual cortex (V1) are affected by optogenetically adding untuned excitatory input to excitatory cells. They found, in matched cells, that optogenetically induced response changes were large in magnitude, as large as the visual responses themselves, yet the distribution of firing rates across the matched population was unchanged, an effect called “rate reshuffing” (10). The authors developed a theory to show that reshuffing can arise in an unstructured, randomly connected network with su#ciently strong coupling, in which the mean response to the untuned stimulus is suppressed by a mechanism of precise cancellation or “tight balancing” of E/I input.

In the theory developed here, we show that structured networks, *e*.*g*. networks with neurons with preferred features, feature-tuned visual input, and feature-dependent connectivity (11–16), allow for activity reshuffing in a realistic “loosely balanced” regime (17). We show analytically that in structured networks that are su#ciently strongly coupled and dominated by feedback-inhibition, the mean matched response to the optogenetic stimulus is suppressed by the residual resulting from the cancellation between the direct optogenetic drive and the indirect baseline-mediated suppression of the matched response. Consistent with this mechanism, we show, in simulations of the full model fitted to experimentally recorded visual and optogenetic response statistics (10, 18), that weakening the network’s feature-dependent structure reduces rate reshuffing, while strengthening it increases the suppression of the mean matched response.

We next examine the model’s optogenetic responses when presented with two gratings offset by 45 ^°^ whose inputs to cortex add linearly, as when they are presented interocularly (19). We extend the linear response analysis to the case of two interocular stimuli and show analytically that additional stimuli weaken the baseline suppression of the matched response, compared to a single stimulus. This results in the mean responses of the two matched populations being slightly increased by the optogenetic stimulus whereas the one-grating mean response is unchanged by the laser. This leads to the prediction that an untuned optogenetic input would result in weaker response normalization than without an optogenetic perturbation (19–22). In summary, our theory provides a foundation for studying the effects of biologically-relevant untuned signals on sensory cortex and provides a general framework for understanding how tuned sensory and untuned modulatory inputs interact in feature-dependent cortical networks.

## Results

### Baseline-plus-Gaussian approximation characterizes ring model dynamics by the rate moments at three sites

We study a recurrent network model of the visual system, consisting of multiple excitatory (E) and inhibitory (I) rate-neurons located at each of multiple evenly spaced sites on a ring (23, 24) (Fig. 1a, see Supp. Eq. (S1) in the Extended Methods for the network equation). We parameterize these sites by a circular variable *θ* that ranges from 0 ^°^ to 180 ^°^, representing the orientation preference of cells located at a given site. For simplicity, we model the connection probability shape to depend solely on the presynaptic cell type, *E* or *I*. The connection probability *p*_*B*_ (*θ, θ*^′^) from a cell of type *B* at site *θ*^′^ to a cell of type *A* at site *θ* between two cells is proportional to a wrapped Gaussian function of the difference in preferred orientations. The synaptic efficacy of nonzero synapses is constant for a given pre/post cell type pair. The cells receive feedforward visual input peaked at the site corresponding to the stimulus orientation, and decreasing to a nonzero baseline as a wrapped Gaussian of the difference between stimulus orientation and preferred orientations of the cell. In V1 layers 2/3, the baseline represents input due to the spontaneous activity of layer 4 cells, which show little or no visual response to an orthogonal stimulus (25). The firing rate of each cell is then determined by a nonlinear function of the sum of the recurrent and feedforward inputs. The function is chosen to be a cell-type-specific Ricciardi nonlinearity *ϕ*_*A*_[*x*] (26–28) (Supp. Eq. (S2)).

**Fig. 1.**
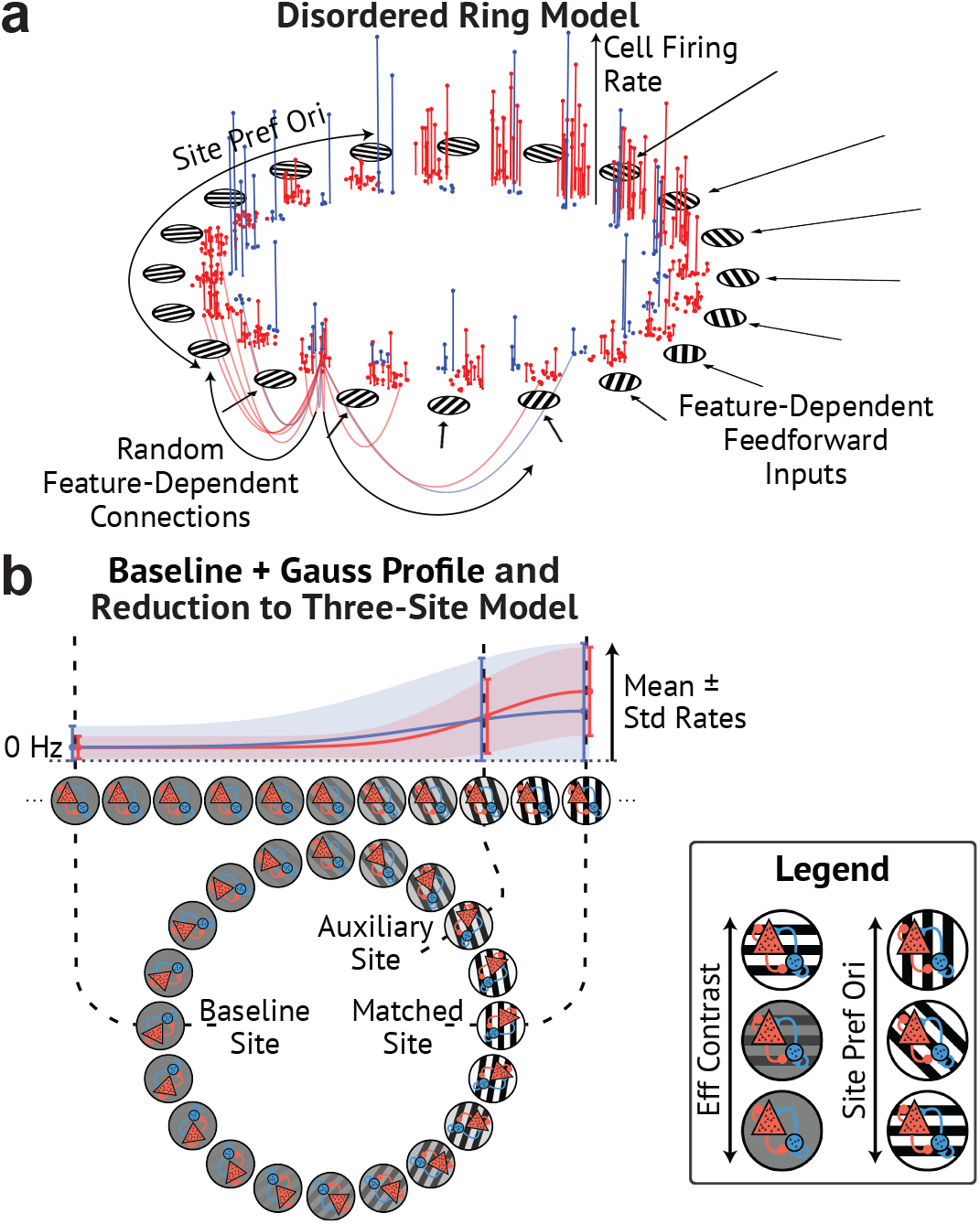
Disordered E/I ring model and mean-field analysis with baseline-plus-Gaussian profile approximation. **a** E (red) and I (blue) neurons are arranged at discrete sites on a ring model of orientation preference. The connection probability decays with the difference in feature (orientation) preference between neurons, while the afferent input to each cell depends on the difference between the cells’ preferred feature and the stimulus feature. Curved lines show connections from one E cell; concentric arrows indicate mean afferent input; vertical stems measure visually-evoked firing rates. We use the term “disordered” to mean connections are assigned stochastically, yielding heterogeneity in cell properties **b** Rate moments (mean and variance) follow an approximate baseline-plus-Gaussian profile on the ring and are fully characterized by the activity at three sites: the baseline, the visual stimulus site (the peak), and an auxiliary location used to infer the tuning widths. Circles represent E/I populations at each site; grating orientation and contrast indicate preferred orientation and relative feed-forward input (see legend). Above: feature-dependence of equilibrium mean ± std of E (red) and I (blue) rates under our feature-dependence approximation for the top half of the ring. Circles and caps highlight statistics at the three aforementioned sites.

We combine two techniques to statistically describe the neuronal activity on the ring (details in Methods). We first use standard mean-field theory for sparse diluted networks (10, 29) to self-consistently compute the steady-state moments of each population of cell type *A* located *θ* degrees relative to the visual stimulus. The rate moments – *i*.*e*. the mean 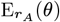 and autocorrelation function 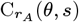 at time lag *s* – are averaged over the population at site *θ* in the limit of an infinite number of such cells. Second, inspired by previous studies (30–32), we approximate the feature-dependence of each cell type’s rate moments as a baseline-plus-wrapped-Gaussian (Fig. 1b). Using superscripts *b* and *m* to denote the baseline and matched values of each rate moment, and letting σ denote the Gaussian tuning widths, the rate moments can be parametrized as:

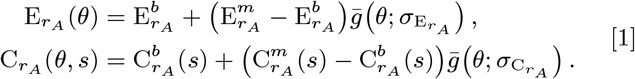

where 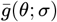 is a normalized baseline-subtracted wrapped Gaussian of width σ satisfying 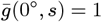 and 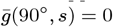 (see also Supp. Eq. (S8) in the Extended Methods). The autocorrelation tuning width 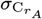 empirically has weak timelag-dependence (Supp. Fig. S1a), so for simplicity we take it to be time-lag-independent.

Given the rate moments, we can infer the distribution of the net recurrent E and I input received by cells in each population. The recurrent inputs of type B (E or I) received by a population of cell type *A* at site *θ* are approximately normally distributed and are characterized by the site-dependent mean 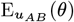 and autocovariance function 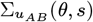 at time lag *s*. For rate moments parameterized by Eq. (1), the statistics of the recurrent inputs are given by

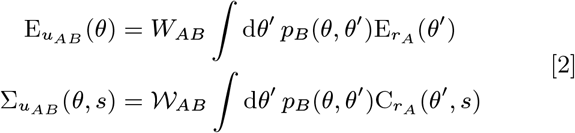

where *W*_*AB*_ is the mean total weight, summed over all presynaptic cells of type *B*, received by cells of type *A*, and 𝒲 _*AB*_ is the variance of the total weight. The convolutions in Eq. (2) are analytically tractable, resulting in closed form solutions (see Supp. Eq. (S11) in the Extended Methods for exact form of the recurrent input statistics).

The equilibrium rate moments satisfy the self-consistent equations

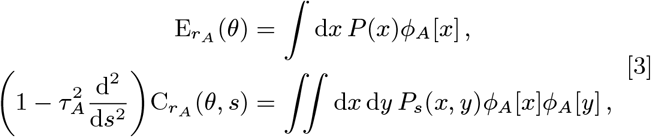

where *P*_*s*_(*x, y*) is the probability density function of the bivariate normal distribution

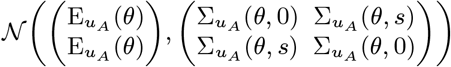

and *P*(*x*) = ∫ d*y P*_*s*_(*x, y*), which is independent of *s*. 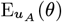 and 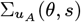 are the mean and autocovariance function of the normally distributed net inputs received by cells of type *A* at site *θ*. If the feedforward inputs are not normally distributed, then Eq. (3) must be separately integrated over the feedforward input distribution and the normally distributed recurrent inputs (see Supp. Eq. (S17) of the Extended Methods for the general self-consistent equations with non-normally distributed feedforward input). Since each rate moment per cell type in Eq. (1) has only three parameters (the matched rate, the baseline rate, and the Gaussian tuning width), we can specify these by evaluating the rates at three representative sites on the ring: the stimulus orientation, a baseline location far from the visual stimulus (*i*.*e*., orthogonal to the stimulus orientation), and an auxiliary site used to infer the Gaussian widths (Fig. 1b) (32). Evaluating Eqs. (1) to (3) at these locations yields a reduced set of self-consistent equations that characterizes the dynamics on the ring by the activity at the three sites.

### To characterize stimulus responses with and without an additional perturbation, 5 quantities must be calculated

This three-site framework can be extended to describe the activity in response to stimuli, and in response to an external perturbation (as an optogenetic perturbation). The response moments to stimuli alone are described by 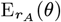 and 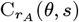. We can call the response moments to the stimulus plus the perturbation 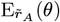 and 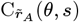. Both sets of moments result from solving the three-site reduction of Eqs. (1) to (3) with the corresponding net input distributions. In many cases, we are not only interested in the activity distribution with and without the perturbation, but also in the distribution of perturbation-induced changes in individual cell rates 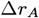, which are not fully determined by the rate moments with and without perturbation. Although the mean perturbation-induced response is given by 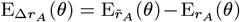, the autocorrelation function of the perturbation-induced responses 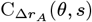 must be separately computed (see Extended Methods for self-consistent equation of 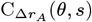). Empirically, we find that a perturbation that is statistically homogeneous across the ring results in a response autocorrelation function whose feature-dependence is also well-approximated by a baseline-plus-Gaussian. Thus, these 5 quantities can be solved by a three-site mean-field analysis.

### Fit ring model generates matched rate reshuffling matching experimental data

We now apply our theoretical framework to investigate rate reshuffing, by which non-specific optogenetic perturbations of excitatory cells strongly affect individual responses of matched cells, but leave the distribution of their firing rates unchanged (10). We use data from (18), consisting of the trial-averaged mean rate ⟨ *r* ⟩ per stimulus for each cell of neurons in macaque V1. The recorded cells are presented with their preferred visual stimuli at varying stimulus contrasts and either in the presence or absence of the optogenetic stimulus. We compute time-averaged model responses either from numerically simulated activity or from solutions of the three-site mean-field equations (Fig. 2a) (see Extended Methods for procedure to compute time-averaged statistics from rate moments). We fit model parameters by constraining numerical simulations to reproduce i) the mean and standard deviation of the matched cells in the absence (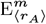 and 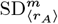) and presence (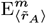 and 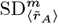) of optogenetic input, and ii) the standard deviation of the optogenetically-induced responses 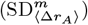, at all visual contrasts presented to the animal (Fig. 2a, see Extended Methods for fitting procedure).

**Fig. 2.**
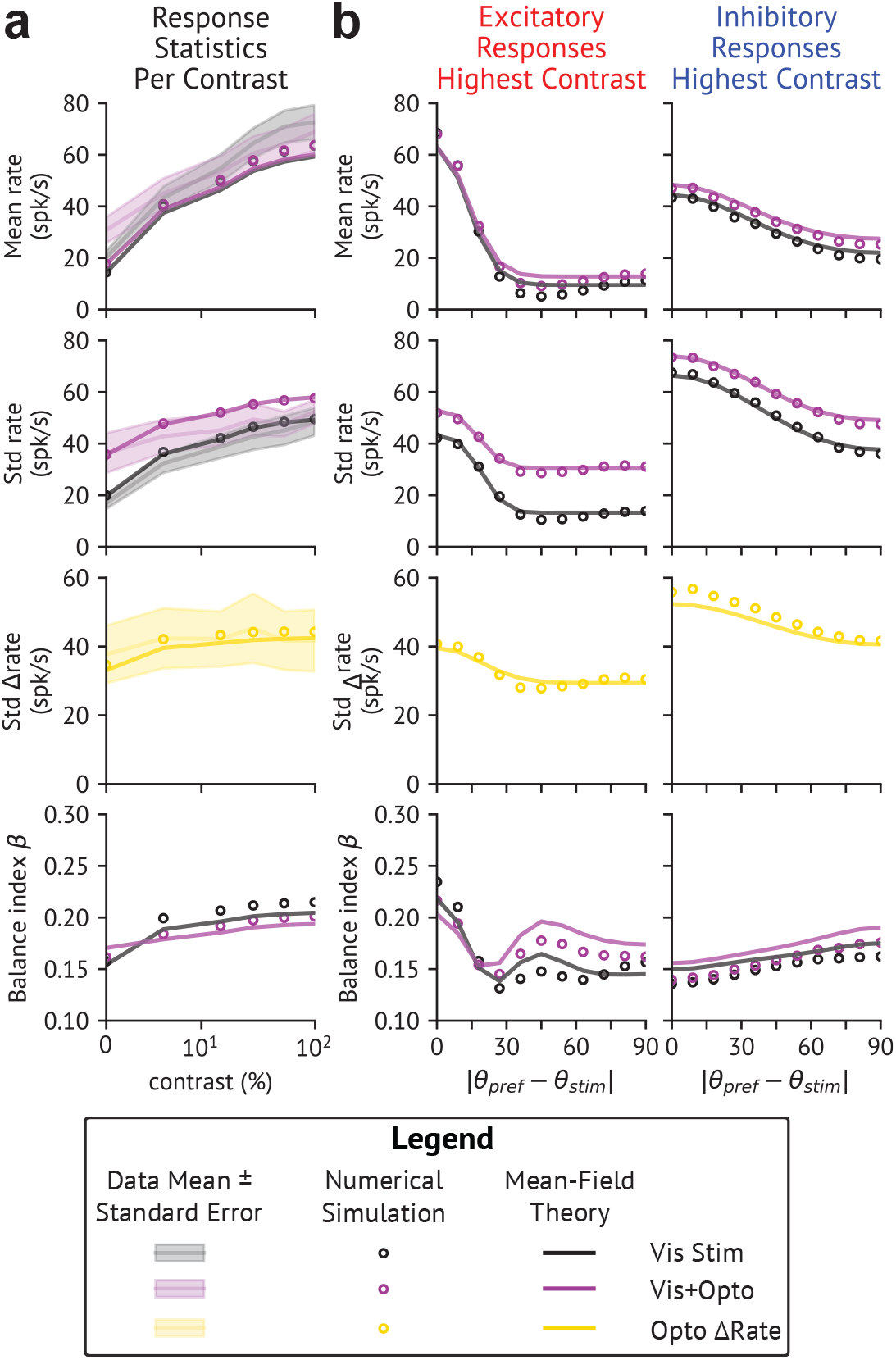
Feature-dependent response statistics show rate reshuffling is strongest in matched populations. **a** Best-fit simulations and theory match experimental data (18) across visual-stimulus contrasts, and do so in a loosely balanced regime (*i.e*. with balance index *β >* 0.1, see Methods). Data shown as bands, simulations as open circles, and theory as solid lines. **b** Simulations and theory demonstrate strongest rate reshuffling (least optogenetically-induced changes) in response statistics in matched populations. Response statistics at highest contrast for E (left) and I (right) populations plotted against absolute distance from preferred orientation to stimulus orientation.

The fitted model successfully captures the dependence of the response statistics of the matched cells on stimulus contrast and optogenetic stimulation. The mean-field values are within 10% of numerical simulations, in which discrepancies are primarily due to deviations from the assumed feature-dependence and finite size effects (Supp. Fig. S2a,b). Comparing theory and simulations across orientation preferences (Fig. 2b) shows qualitatively good agreement. In addition, our model makes a directly testable prediction: visual-stimulus-matched cells are more strongly reshu”ed than mismatched cells.

The model exhibits emergent behaviors beyond our fitting criteria. It produces rate reshuffing in a loosely balanced regime (balance index *β* > 0.1, where *β* is the average over the ratio of cells’ rectified net inputs after E/I cancellation to their E input alone) (17, 33), contrasting with previous unstructured models which required tight balance (10) (Fig. 2a,b, final row). In addition, not only are the rate moment tuning widths nearly invariant across visual stimulus contrasts and optogenetic conditions (Fig. 3a), but individual cells’ tuning curves also exhibit approximate contrast-independence (Supp. Fig. S3e), consistent with V1 observations (25, 34, 35). Intriguingly, the model also predicts that the optogenetic stimulus does not strongly affect individual cells’ orientation selectivities and tuning widths (Supp. Fig. S3a-d).

**Fig. 3.**
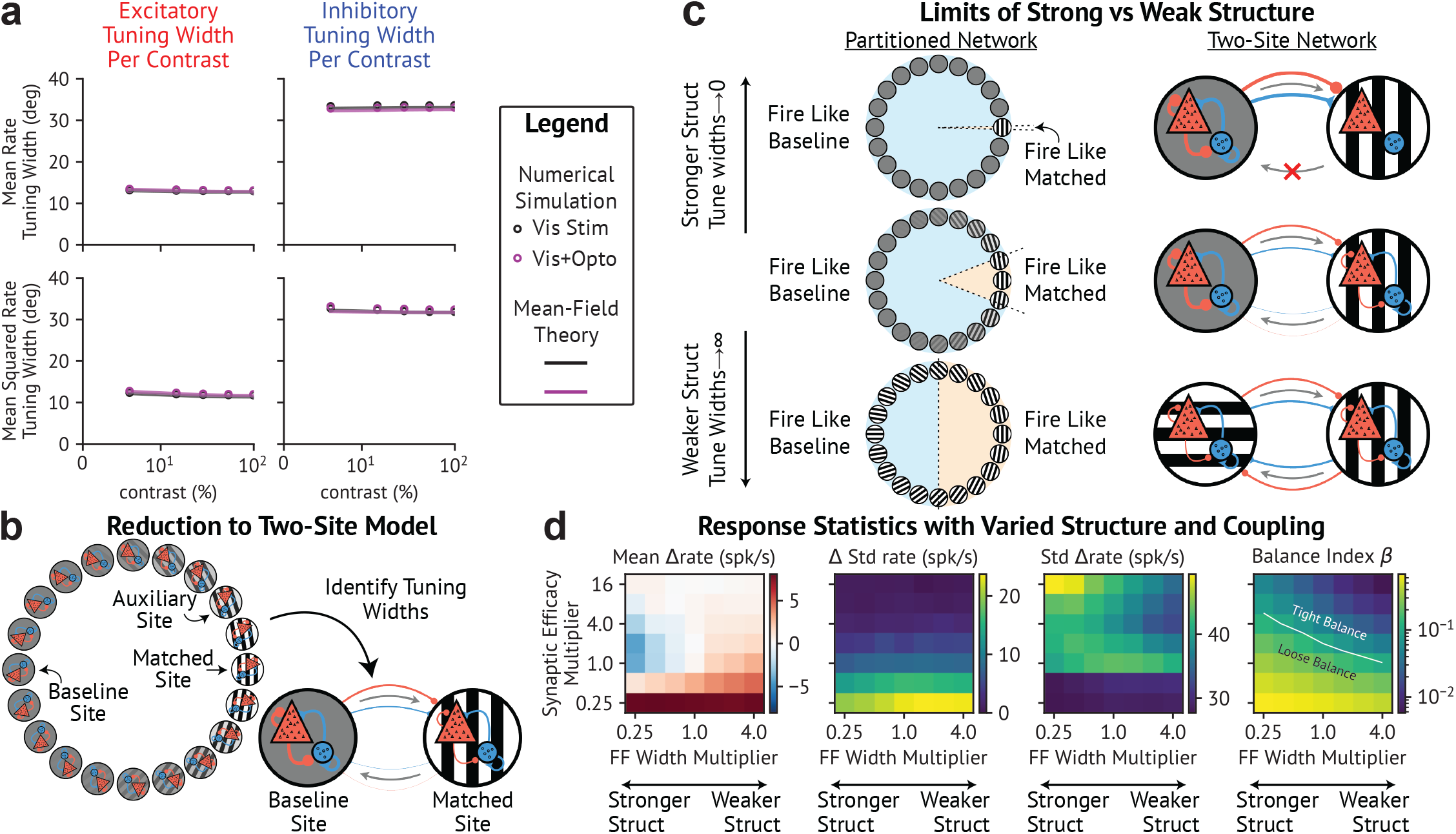
Effective two-site model is possible due to stimulus-invariant width of the rate moments. **a** Mean-rate and mean-squared-rate tuning widths from simulations and theory remain nearly invariant across stimulus conditions. **b** After identifying tuning widths, the network is fully characterized by the baseline and matched activity, allowing reduction to an effective two-site model. **c** Partitioning cells based on preferred orientation approximately determines mean projection strengths in effective two-site model. Cells are categorized as “firing like the matched activity” if their mean rates are at least 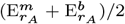 while all others are categorized as “firing like the baseline”. Asymmetric effective baseline-to-matched feedback strengthens in the strongly structured, narrow tuning width limit. In the weakly structured, wide tuning width limit, with uniform visual input the network becomes equivalent to an unstructured model. **d** Reshuffling weakens as network structure is reduced by increasing all widths. Strengthening structure increases mean-response suppression but shows no clear pattern for other statistics. Plots show simulated matched-response statistics varying feedforward input widths (x-axis) and synaptic efficacy (y-axis), where in both cases 1.0 represents the value used in our fits (Fig. 2).

### Networks can be reduced to an effective model between matched and baseline populations

Given that the rate moment tuning widths are approximately independent of the stimulus contrast and optogenetic conditions (Fig. 3a), these widths can be determined once and fixed. Then, the auxiliary rate moments used to infer the Gaussian widths can be dropped, allowing us to derive a reduced effective model. In this model, E and I activities are each described by the rate moments of two sites (the matched and the baseline sites), capturing the essential network interactions (Fig. 3b; see Extended Methods). The effective connectivity between sites *μ, ν* = {*b, m*} in this two-site model is given by

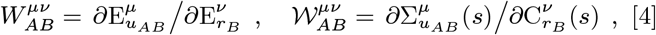

where 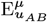 and 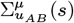 are the statistics of the recurrent inputs of cells at the corresponding site on the ring (see Supp. Eq. (S25) & (S26) in the Extended Methods for their exact forms). Note that 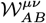 in Eq. (4) is time-lag-independent since 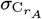 is assumed to be time-lag-indepenent.

### Feature-dependent structure creates strong, asymmetric effective baseline-to-matched feedback in the reduced model

To gain intuition for the effective connectivity in the reduced model, we can imagine partitioning the network into two groups. For each cell type, we categorize cells in the interval within which the mean rates are at least 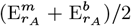 as “firing like the matched activity”. The remaining cells are categorized as “firing like the baseline”. The mean weights within or between the two groups, which approximately scale with the proportion of cells within each partition, would roughly match the mean weights of the effective two-site model (Eq. (4); Supp. Fig. S4b).

For moderate tuning widths, on the order of experimentally reported values (Gaussian tuning widths ∼ 15 ^°^ or full-width at half-height (FWHH) ∼ 40 ^°^) (36–38), most cells fall outside the interval satisfying 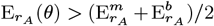, so the average neuron on the ring will receive more connections from cells firing like the baseline than from cells firing like the matched activity. Therefore, in this picture, the baseline-to-matched projections are much stronger than matched-to-baseline, an intuition that is di#cult to appreciate from Eq. (4) alone (Fig. 3c, middle). As a result, untuned stimuli targeting both matched and baseline populations not only drive the matched response directly but also indirectly through the baseline-to-matched projections.

In the limit of small tuning widths, which yield strongly structured responses, nearly all cells fire like the baseline, so the effective model only contains projections from the baseline to the matched site, and not vice-versa (Fig. 3c, top). Conversely, in the weakly-structured limit of large tuning width, half the cells fire like the matched activity and half like the baseline (because the FWHH of a wrapped Gaussian covers half the ring in this limit), leading to symmetric connectivity between baseline and matched sites. In this case, if the baseline and matched sites receive equal feedforward input, the two sites become interchangeable, rendering the network equivalent to an unstructured network with a single site (Fig. 3c, bottom). This partitioning heuristic gives an intuition for why, in the mean-field analysis, the role of structure is to cause the asymmetry by which the connection from baseline to matched sites is much stronger than that from matched to baseline sites.

### Structure robustly strengthens rate reshuffling effect

We now assay how structure and coupling strength each contribute to rate reshuffing. To vary the degree of structure, we varied the width of the afferent inputs while keeping the matched input constant, thus changing the rate moment tuning widths. To vary the coupling strength, we scaled the synaptic strengths.

In agreement with previous results (10), we find that increasing the coupling strength, which causes tighter balance (lower balance index *β*), yields stronger reshuffing as measured by smaller absolute changes in the mean and standard deviation of the activity (Fig. 3d). Increasing network structure (narrowing tuning widths) also strengthens reshuffing and its effects are comparable in magnitude to those of increasing the synaptic efficacy: for all but the weakest coupling, increasing structure decreases the mean optogenetic response and increases heterogeneity of the optogenetic response as measured by the standard deviation of the optogenetically-induced change in rate. Thus, our model provides another testable prediction: narrowing neurons’ tuning curves, either by increasing spatial frequency of the presented grating (39, 40) or by controlled PV or SST activation (41), will yield a more strongly suppressed matched optogenetic response.

In large unstructured networks, reshuffing can be achieved by tight balance induced by strong coupling (10), but the mean response always remains positive, whereas in structured networks, the mean response can be zero or negative. However, we find that the change in the overall standard deviation of the population’s rates is only weakly modulated by structure, whereas it is more sensitive to changes in coupling.

### Linear response analysis reveals inhibitory cross-site feedback

To develop an intuition of the contributions of network structure to the responses to perturbations, we compute the linear response of the reduced two-site model. We label sites as *μ* = *b* for the baseline and *μ* = *m* for the matched site. To highlight the main features of the linear response, we will only discuss a “mean-interaction approximation” in which we focus solely on the linear *mean* response 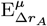 while assuming the perturbation does not change the rate autocorrelation function. Empirically, this approximation yields results comparable to the linear response including rate autocorrelation changes (Fig. 4, see also Eq. (13) of the Methods for the linear response including rate autocorrelation changes).

**Fig. 4.**
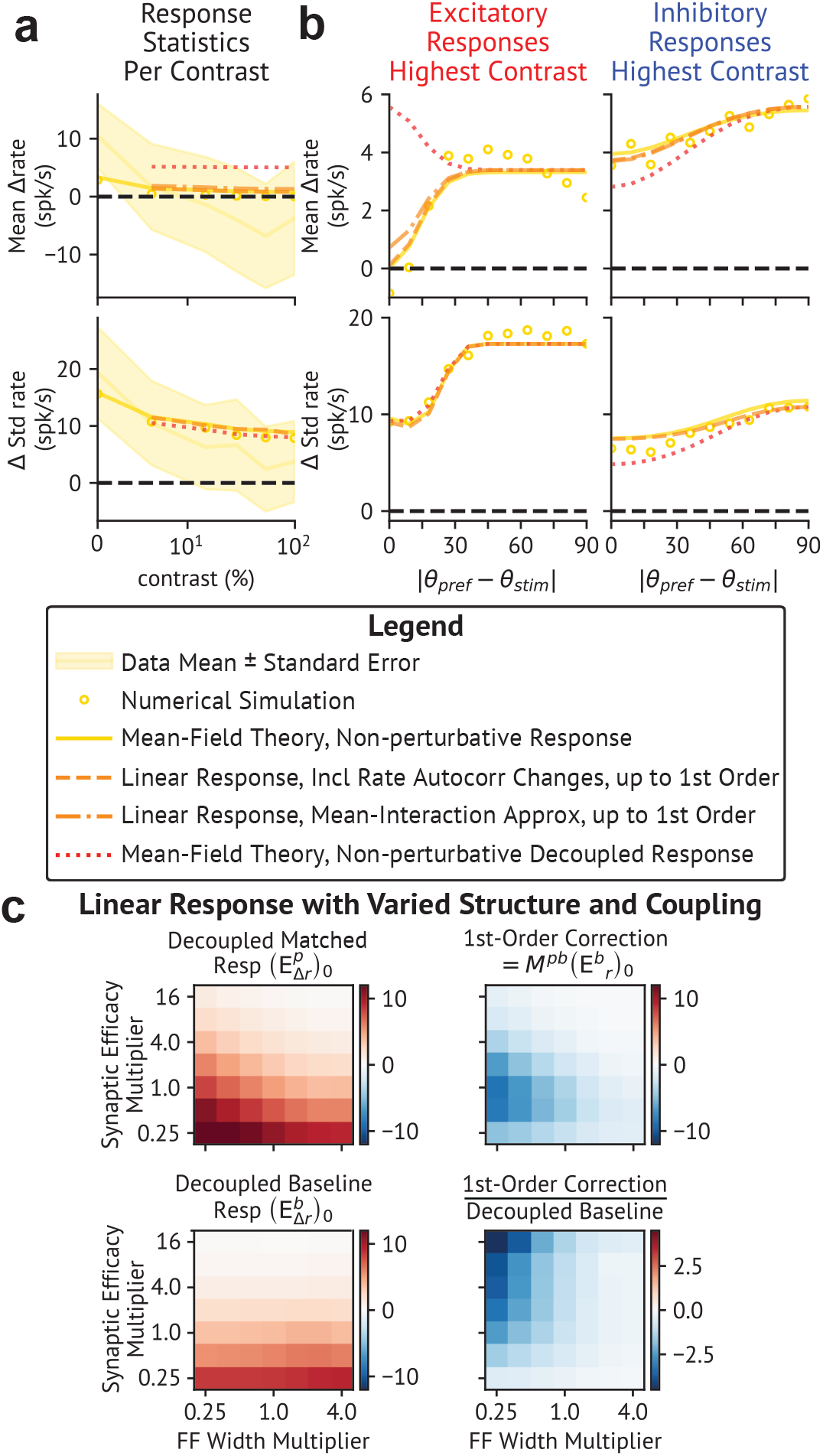
Linear response analysis reveals baseline-induced suppression of the matched response. **a** Effective cross-site feedback is crucial for strong suppression of matched-mean response, but not for rate-standarddeviation response suppression. Yellow bands, circles, and lines show values from the data, simulations, and three-site mean-field theory, respectively. Orange lines show truncated linear responses (dashed: linear response including rate autocorrelation response; dash-dotted: mean-interaction approximation) whereas red dotted lines show decoupled responses. **b** First-order linear response with effective baseline-to-matched feedback accurately predicts the feature-dependence of nonlinear optogenetic stimulus responses. Highest contrast response statistics shown for E (left) and I (right) populations versus absolute distance from preferred to stimulus orientation. **c** Increasing structure causes stronger baseline-to-matched effective interactions, driving increased matched response suppression. Plots show linear response analysis results computed in the mean-interaction approximation while varying feedforward input widths (x-axis) and synaptic efficacy (y-axis), where in both cases 1.0 represents the value used in our fits. The top row shows the decoupled matched response and first-order correction to the matched response. The bottom row shows the decoupled baseline response and the ratio of the first-order matched response correction to the decoupled baseline response.

Given the limited spatial range of lateral connections, the majority of outward connections from matched or baseline sites are predominantly received by the same site, as can be seen from the analogy of partitioning the ring into baseline and matched segments (Supp. Fig. S4c). Thus cross-site projections are generally weak compared to within-site projections, allowing us to express the linear response as a series expansion in powers of the cross-site interactions. For each cell type *A, B* and each site *μ*, ν = {*b, m*}, this series expansion is given by:

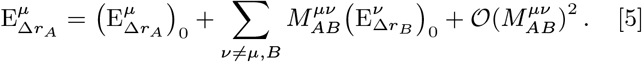

Here, the effective cross-site interaction matrices 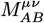 encode how the mean response of cell type *B* of site ν affects the mean response of cell type *A* of site μ ≠ ν Note that, because the expansion is in the cross-site projections, which are assumed small and treated as a perturbation, there are no couplings *M*^*μ μ*^ a site to itself (see Methods and SXX of Extended Methods). The zeroth-order responses 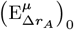 cannot actu -ally be computed in linear response, because the optogenetic perturbation is not normally distributed (see Methods). Therefore, in our calculations, we compute this as the full mean field result when the baseline and matched sites are decoupled, and refer to it as the “decoupled response” since it includes no dynamic cross-site feedback.

If we neglect matched-to-baseline feedback due to its relative weakness, *i.e*. set 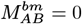, then we can exactly truncate our expansion in Eq. (5) to express the matched and baseline linear response as

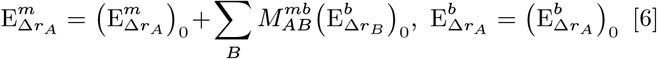

The two terms in the expression for 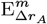 formally define the direct and indirect components, respectively, of the matched response to an untuned perturbation.

The matrix 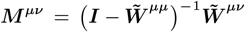 for *μ* ≠ ν with elements 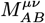, is given by

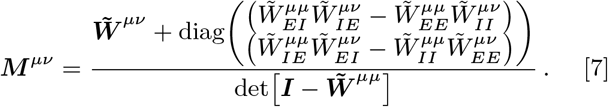

Here 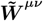 is the matrix of the effective *μ* →ν couplings whose elements are 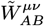. Without disorder, these couplings are the product of the mean input-output gains of the pre-synaptic cell type and the mean recurrent weight strengths (Eq. (4)). Disorder and the activation function’s nonlinearity augments this relationship (Supp. Eq. (S85)). Our perturbation theory depends on the equilibrium rate moments (Eq. (3)) being stable to perturbations of the input in the decoupled two-site network, which implies the determinant in the denominator of Eq. (7) is positive (29) (see Appendix I in the Supplement for clarification). Eq. (7) comprises a monosynaptic term and a diagonal disynaptic term, with the latter dominating in strongly coupled networks. If feedback inhibition is strong, this disynaptic term becomes purely inhibitory. Empirically, the determinant in Eq. (7) is positive (Supp. Fig. S1b) and the disynaptic term is generally purely inhibitory (Supp. Fig. S1c) for our best-fit model even as we vary coupling and structure. Therefore, in strongly coupled and feedback-inhibition-dominated networks, the indirect effect of an untuned stimulus on the matched response is suppressive.

### Structure induces strong baseline-to-matched inhibition that drives rate reshuffling

We now apply our analysis to elucidate the feature-dependent mechanisms underlying rate reshuffing. We compare numerical simulations, which give us the ground truth of model behavior, to four analyses that incorporate different aspects of structure (Fig. 4d): 1) the response 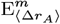 and 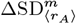 computed using the non-perturbative three-site mean-field framework, which tells us how well our overall mean-field framework works; 2) the linear response including optogenetically-induced changes in the rate autocorrelation truncated to first order in the matched site. 3) the linear response, but now computed in the mean-interaction approximation, given by Eq. (6). Note that, since the mean-interaction approximation neglects changes in the rate autocorrelation function, that method cannot accurately compute standard deviation changes and thus we omit those plots from Fig. 4a,b. Finally, 4) the decoupled response 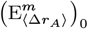 and 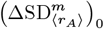, computed non-perturbatively in mean-field theory, which tells us how the network responds to the perturbation without interactions between the effective sites (*i.e*. between the baseline and the matched site).

Each of the four methods computes neural response statistics with different sets of approximations, and comparing them with simulations and between each other informs us of how good these approximations are. The three-site mean-field framework assumes the rate moments are well-described by a baseline-plus-Gaussian profile. In agreement with our previous comparison between simulations and mean-field values (Fig. 2a,b), we find strong alignment with simulation results both as a function of visual contrasts and across orientation preferences (Fig. 4a,b respectively). The remaining methods assume that the Gaussian part of the feature-dependence describing a given rate moment has a stimulus-invariant tuning width, allowing construction of an effective two-site model. The truncated linear response including rate autocorrelation changes further assumes that the effective matched-to-baseline projections are negligible. We find that this approximation closely matches the three-site mean-field result, which validates both the weak matched-to-baseline assumption and the stimulus-invariant tuning width assumption. The truncated linear response computed in the mean-interaction approximation additionally neglects changes in the rate autocorrelation when computing mean responses, and again exhibits good agreement with the mean-field calculation. These results suggest the mean-interaction approximation, which was used to derive the form of the effective interaction matrices (Eq. (7)), can accurately compute optogenetically-induced changes in mean activity.

Finally, the decoupled response represents changes in activity in the absence of cross-site interactions. In the partitioned ring analogy (see Fig 3c), decoupling amounts to severing connections between cells firing like the matched activity and cells firing like the baseline so that each group is a separate unstructured network. The role of structure is to create strong asymmetric projections between them, primarily from baseline to the matched site, which are lost in the decoupled network. Since the matched site network in the decoupled case can be thought as an unstructured network, we expect it to show larger mean responses than in the coupled case, and indeed we find that the decoupled response produces a larger change in the mean matched response (Fig. 4a) and the excitatory matched response (Fig. 4b) than the simulations and the other three methods. Assuming matched-to-baseline projections are negligible, decoupling should not significantly change the baseline response, which is indeed what is found (Fig. 4b).

From our analysis of the effective cross-site interactions in the mean-interaction approximation (see Eq. (7)), we predicted that in strongly-coupled, feedback-inhibition-dominated networks, the baseline-to-matched interaction is net inhibitory. In agreement with this prediction, while the mean excitatory decoupled response is positive (dotted line in Fig. 4b), the addition of the first order correction to the linear response, either with or without rate autocorrelation changes (dashed and dot-dashed lines in Fig. 4b), largely suppresses the excitatory response. This means that the first-order correction must be comparable in magnitude to the decoupled response, but inhibitory. The truncated linear response of the inhibitory population, on the other hand, is more excitatory than the decoupled response, which shows that the effective interaction matrix need not be purely inhibitory across all cell types. Additionally, the decoupled and linear response calculations of standard deviation changes yield similar results, suggesting that the network structure is not essential for suppressing changes in higher-order rate moments. This structure-independence of the standard deviation changes corroborates our previous finding that this quantity is more strongly modulated by changing the network’s coupling than by changing its structure (Fig. 3d).

### The baseline’s suppression of the matched response almost entirely structure dependent given sufficient coupling

We previously showed that strengthening the network structure (narrowing the tuning width) yields more inhibited matched optogenetic responses, given su#ciently strong coupling (Fig. 3d). We hypothesize that this effect arises because increased structure results in stronger effective baseline-to-matched inhibition (as opposed to increased structure making the decoupled matched response more inhibitory). To test this hypothesis, we recompute the zeroth order decoupled matched response and the first-order correction in the mean-interaction approximation while independently adjusting the afferent input width (decreasing input width corresponds to stronger structure) and the synaptic coupling strength (Fig. 4c). The first order correction computed with and without rate autocorrelation changes largely agree (Supp. Fig. S5b), but we focus on the mean-interaction approximation since it permits an interpretable analytic solution. Consistent with our hypothesis, the decoupled matched response is always positive and increases with increasing structure at all coupling strengths. In fact, the strength of the decoupled response is strongly correlated with the balance index: decoupled matched response strength increases nearly monotonically with increasing balance index (looser balance, which results both from stronger structure and from weaker coupling) (Supp. Fig. S5a). Thus, stronger structure yields a more excitatory decoupled matched response. The first-order correction to the linear response, on the other hand, is always negative, and grows stronger with stronger structure at fixed coupling, in agreement with our

The first-order correction to the linear response, on the other hand, is always negative, and grows stronger with stronger structure at fixed coupling, in agreement with our hypothesis (Fig. 4c, top right panel). This first-order cor-rection, 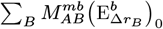 is equal to the product of the effective baseline-to-matched interaction matrix and the decoupled baseline response. To further break down why this term grows more strongly inhibitory with structure, we separately study the decoupled baseline response and the ratio of this first-order correction term to the decoupled baseline response (Fig. 4c, bottom panels). The ratio, and thus the effective interaction matrix, is always negative and grows stronger with increasing structure for all but the weakest coupling (Fig. 4c, bottom right panel). The decoupled baseline response is always positive, and is largely independent of structure. Thus, the increase in suppression of matched optogenetic responses with increasing structure is due to the strengthening of the effective baseline-to-matched projections, which is induced by structure provided coupling is not very weak.

### Reduced mean-field equations and linear response analysis generalize to multi-matcheded rate profiles

Our approach can be extended beyond single gratings to multiple visual stimuli. While such patterns invalidate the baseline-plus-Gaussian assumption, our analyses can be generalized if the rate moments are well approximated by a baseline-plus-Gaussian-mixture (*i.e*. a sum of Gaussians). In this extended framework, Eqs. (1) and (2) involve sums of *N* Gaussians with 2*N*+1 degrees of freedom per rate moment per cell-type: the theoretical baseline response (which may not correspond to activity at any physical site on the ring if the rate moments never fully decay to baseline), the *N* matched responses, and *N* auxiliary sites used to determine each Gaussian’s tuning width. Once the *N* tuning widths are determined, and if they remain approximately stimulus invariant, they can be fixed to derive an *N*+1-site effective model.

We often study configurations of visual inputs that give rise to identical firing rate distributions and tuning widths for populations matched to any of the visual stimuli. This symmetry permits defining a further reduced three-site mean-field framework and an effective two-site model akin to our earlier single-grating results. The linear response of the generalized effective models and the effective interactions (Eqs. (5) and (7)) are unchanged.

### Optogenetic input decreases response normalization

Presentation of multiple visual stimuli in a neuron’s receptive field can yield sublinear response summation or “normalization”. This effect can be due to sublinear summation (or saturation) of the inputs induced by each stimulus to cortex (42, 43), or emerge from recurrent interactions driven by linearly-summing inputs induced by the stimuli. The latter case describes the scenario where different stimuli are presented to each eye (19), in which case normalization is also observed. We will refer to normalization observed when inputs add linearly as “interocular normalization”. We investigate whether interocular normalization could impact rate reshuffing in response to optogenetic stimulation, by adding an equal-contrast interocular visual stimulus oriented 45^°^ away from the first grating. We then model the feedforward input as the linear sum of the untuned baseline input driving spontaneous activity and two offset Gaussians.

As expected, the two-grating-evoked matched mean rates are lower than the single-grating-evoked activity, indicating sublinear response summation (Fig. 5a,b). As in the onegrating-driven network, an untuned optogenetic perturbation, added to the interocular visual stimulus, more strongly suppresses changes in the activity statistics in populations matched to either of the two visual stimuli than in non-matched populations. However, both stimulus-matched sites show a small but noticeable increase in their optogenetically-induced mean response compared to the single-grating network. As a result, the strength of response normalization is reduced by the optogenetic stimulus. This can be quantified by the normalization weight (ratio of two-grating to one-grating matched mean rates). Since the mean two-grating, but not the mean one-grating, response is increased by the optogenetic input, the addition of the optogenetic input yields weaker normalization (normalization weight closer to unity, Fig. 5c).

**Fig. 5.**
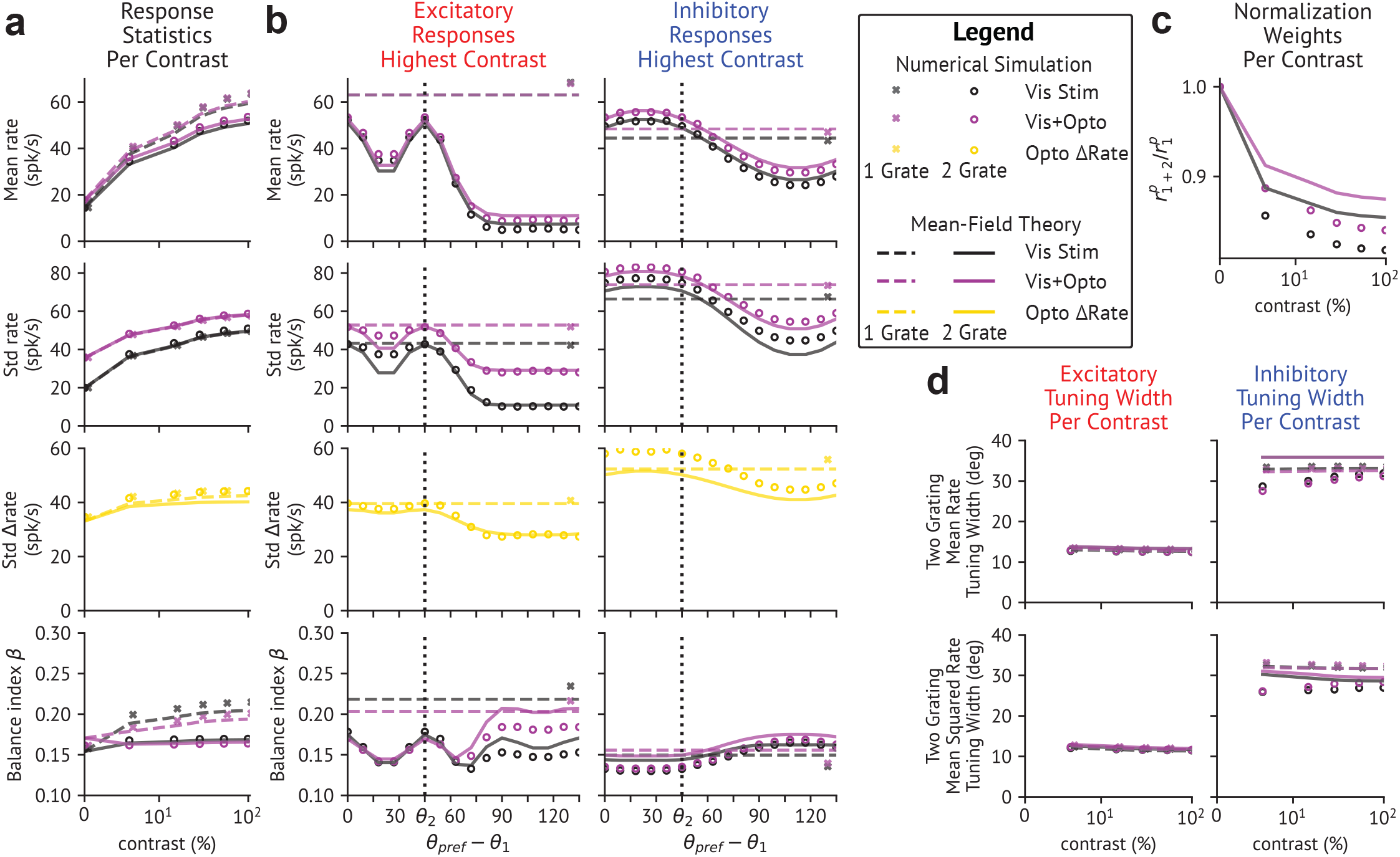
Optogenetic stimulation increases mean matched responses with two gratings present. **a**,**b** Rate reshuffling is slightly weakened in the two-grating driven network. Response statistics for matched cells are shown in roughly the same layout as Fig. 2a,b. Matched response statistics of the one-grating driven network are plotted as ‘x’ markers for simulation results and dashed lines for three-site mean-field computed values. **a** One and two-grating response statistics as a function of contrast. **b** E (left) and I (right) population response statistics at the highest contrast relative to the left-most grating orientation. Vertical dashed line indicates the location of the second grating orientation. Horizontal dashed line indicates one-grating matched response statistic from mean-field theory whereas the ‘x’ marker indicates matched response statistics from simulations. **c** Optogenetic stimulation weakens normalization across all contrasts. Normalization weight (ratio of two-grating to one-grating matched mean rates) vs contrast. **d** Tuning widths remain nearly stimulus-invariant, similar to the single-grating condition.

### Additional visual stimuli weaken baseline-to-matched inhibition, in turn weakening mean response suppression

In the previous section we showed that two gratings increase the optogenetically-induced mean response compared to a single grating, leading to weaker normalization. We now investigate why interocular visual stimulation results in weaker reshuffling than single grating stimuli. Analogously to single-grating driven networks, the two-grating driven tuning widths are nearly stimulus invariant (Fig. 5d), which when combined with the identical matched response statistics and tuning widths of each Gaussian component allows us to construct an effective two-site model of the interactions between the identical matched responses and the baseline (Fig. 6a). Neglecting matched-to-baseline projections results in the truncated linear response expansion series of Eq. (6).

**Fig. 6.**
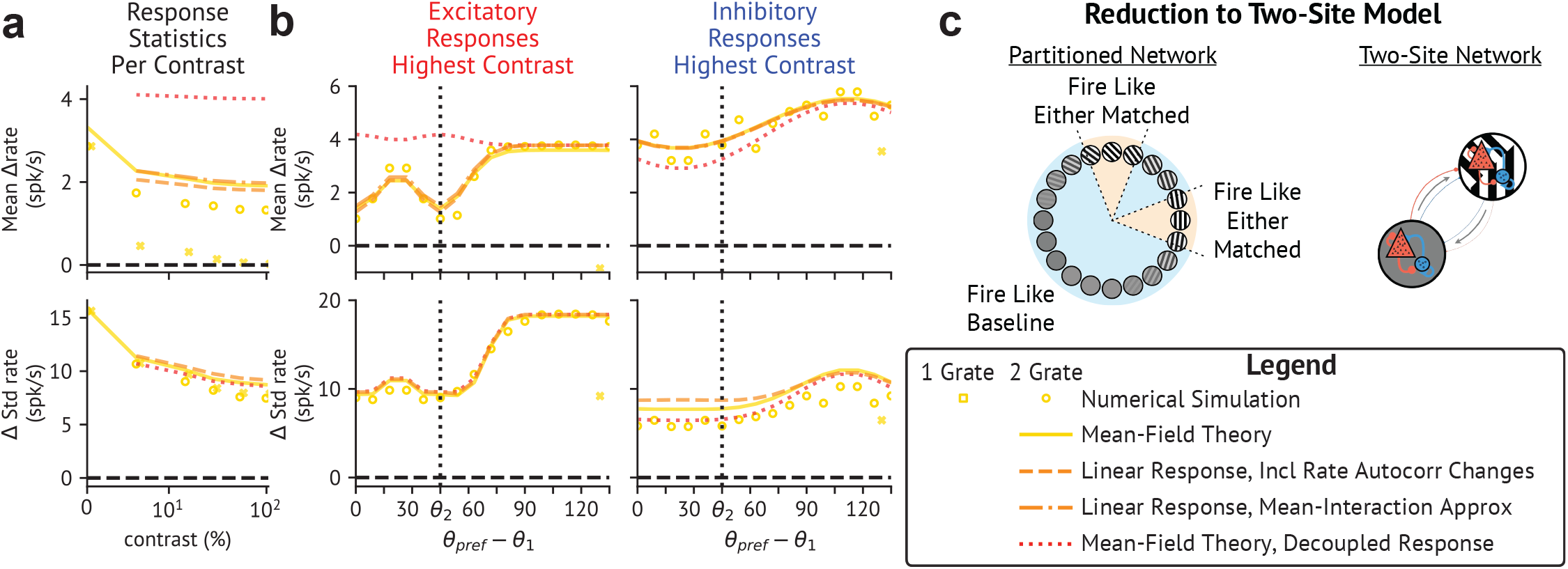
Strong effective matched-to-matched coupling and weaker baseline-to-matched feedback with multiple visual stimuli increase susceptibility of mean-matched responses to untuned modulation. **a**,**b** In the presence of two interocular stimuli, weaker baseline-to-matched projections yield weaker suppression of the mean response. Response statistics are shown in roughly the same layout as Fig. 4a,b. **a** Response statistics as a function of contrast. **b** E (left) and I (right) population response statistics at the highest contrast relative to the vertical grating orientation. **c** If both gratings have equal contrast, then cells firing like either matched population can be combined into a single partition to derive an effective two-site model. This results in more a strongly coupled matched population and weaker baseline-to-matched projections compared to the one-grating network.

As in the one-grating network, we repeat the four analyses computing the optogenetically-induced changes in response statistics for the two-grating driven network. We find qualitatively similar results as before when comparing results between each method with simulations and between each other (Fig. 6a,b). However, we find that the two-grating driven network produces a less suppressed matched response than the one-grating ring even though the two-grating decoupled response is more suppressed than the one-grating response.

To understand the origin of these differences, we extend the heuristic analysis based on the idea of partitioning the ring presented for the one-grating configuration in Fig. 3c, to the two-stimuli case studied here. This analysis provides an intuitive understanding, and gives similar effective two-site connectivity compared to the full analysis (Supp. Fig. S6a,b). As before, for each cell type, we categorize all cells satisfying 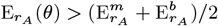 as “firing like the matched activity”. The presence of two response Gaussian components results in fewer cells being categorized as firing like the baseline (compare Fig. 6c with Fig. 3c, middle panel). This causes some baseline-to-matched projections to be diverted into projections across the two matched sites, resulting in stronger matched-to-matched coupling and weaker baseline-to-matched coupling.

The stronger, suppressive matched-to-matched coupling leads to more tightly balanced matched activity (Fig. 5a,b). This explains why the two-grating decoupled matched response is more suppressed than the one-grating response, yielding response normalization (as in (33)). While the additional visual stimulus strengthens the matched-to-baseline coupling, this coupling remains weak enough to neglect, as shown by the strong agreement between the truncated linear response including rate autocorrelation changes and the three-site meanfield framework (Fig. 6c,d). The weaker baseline-to-matched coupling explains the weakened suppression of the mean response to the optogenetic stimulus, compared to the one-grating case. To rule out the alternative hypothesis that this weaker suppression is due to the increased matched-to-matched coupling, we repeat the linear response analysis using the one-grating baseline-to-matched coupling strength and find that this results in fully suppressed matched excitatory responses (Supp. Fig. S7). In summary, we have shown that the weakened optogenetically-induced reshuffing in the presence of multiple visual stimuli and the weakened normalization in the presence of the optogenetic stimulus are both due to the decrease in baseline-to-matched suppression with an increasing number of visual stimuli.

## Discussion

In this study, we investigate how untuned stimuli interact with tuned sensory stimuli in an E/I ring model with feature-dependent connection probabilities. We demonstrate that models with stimulus-invariant tuning widths can be reduced to an effective model describing interactions between the sensory-stimulus-matched and baseline populations. Through this reduction, we show that feature-dependent structure creates strong baseline-to-matched connections, but only weak connections in the reverse direction. Through a linear response analysis, we reveal that the baseline-to-matched connection is suppressive given su#ciently strong coupling and feedback inhibition. Thus, in this regime, untuned stimuli directly drive the matched response, and indirectly suppress it through the strong baseline-to-matched projections.

Previous work (10) showed that untuned optogenetic activation of excitatory cells in macaque V1 caused rate reshuffing, resulting in no significant mean response of matched cells to the optogenetic activation. In models without structure, the data can only be explained in a regime of unrealistically strong coupling, in which strong recurrently-evoked inhibition very largely (but not entirely) cancels the mean external input. Our work gives analytical insight into how structured networks allow reshuffing to occur with more plausible levels of coupling: The indirect suppression of the matched response by the baseline-to-peak connection adds to other recurrent inhibition to allow full suppression of the mean matched response.

Given sufficiently strong coupling, increasing (decreasing) the degree of structure, by input manipulations that narrow (widen) tuning, produces strengthened (weakened) optogenetically-induced mean response suppression. We show that this effect, predicted by our theory, holds in a ring model fit to experimental data (18). By repeating our linear response analysis as we vary the degree of network structure, we verify that, given sufficiently strong coupling, the baseline-to-matched interaction is purely suppressive, with a strength that depends almost entirely on the strength of network structure. Our findings demonstrate how and why a structured network’s response to untuned stimuli depends strongly on the degree of selectivity in the network, *i.e*. its tuning widths.

Finally, we extend our investigation to the case of two equal-contrast visual stimuli, presented interocularly (*i.e*., two stimuli whose inputs to cortex add linearly). The second stimulus weakens the matched responses compared to a single stimulus, *i.e*. there is sublinear response summation or “normalization”. We find that this effect can be explained by a stronger effective matched-to-matched connection (and weaker baseline-to-matched connection) in the presence of the second stimulus, which can be understood heuristically from the increased number of cells matched to the visual stimulus (*i.e*. to either of the two stimuli) and correspondingly fewer cells firing at the baseline. Given su#ciently strong coupling, the matched-to-matched connection is suppressive, so the strengthening of this connection yields normalized responses (33).

Adding an optogenetic stimulus to the two visual stimuli induces a slightly positive mean response of the matched activity, instead of a zero mean response for a single stimulus, due to the weakened baseline-to-matched connection given two visual stimuli. The increased mean response results in weakened rate reshuffing and weaker response normalization under optogenetic stimulation.

In summary, the second grating induces suppression of matched responses, via the strengthened suppressive matched-to-matched interaction, even as it weakens the suppression of the mean response to optogenetic stimulation, due to weakening of the suppressive baseline-to-matched interaction. Because the impact of an untuned stimulus on matched responses depends on the number of visual stimuli present, untuned perturbations can modulate the degree of sublinear response summation, which is a prominent aspect of the canonical computation of response normalization (20, 22).

### Comparison with previous reduction techniques for ring networks

Our work builds upon and extends previously developed analytical methods for analyzing E/I ring networks without disorder. These works assume either an approximately Gaussian response profile (30, 32) or that the net input shape approximates the feedforward input shape, which need not have a Gaussian profile (33). Given the fixed response shape, Ahmadian et al. (33) then reduced the model to a single-site model of the excitatory and inhibitory responses at the matched site, similarly to our reduced models.

Our reduction procedure generalizes these approaches in three ways. First, our mean-field methods enable the analysis of models with disordered inputs and connectivity. Second, our method self-consistently computes the shape of the rate moment profiles, rather than assuming that feedforward inputs drive the response shape. This allows us to capture differences in features selectivity between cell types – *i.e*., broader orientation tuning of inhibitory responses vs. excitatory responses (14, 44–47). Lastly, our procedure explicitly accounts for baseline responses, often overlooked in previous feature-dependent models (30, 33, 48).

Although our analysis considers only a single circular feature, such as preferred orientation, it can be generalized to an arbitrary number of features, provided responses can be approximated by a baseline plus one or more Gaussian response profiles (See Appendix J). It thus potentially offers a robust framework for more generally analyzing and predicting the behavior of feature-dependent neural networks.

### Implications for rate reshuffling

Tight E/I balance is required to produce rate reshuffing in unstructured models (10). Our study establishes the necessary conditions on the structured connectivity – sufficiently strong coupling and feedback inhibition that the disynaptic term in Eq. (7) is suppressive and outweighs the monosynaptic term – for rate reshuffing to occur with loose balance.

Our results may help explain the differences in rate reshuffling between experiments in mice, where neurons were recorded irrespective of visual-stimulus-matching (49), and those in macaques, where only matched cells were recorded (18). The mouse dataset exhibited only weak reshuffing at the highest visual stimulus contrasts, which was previously attributed to weaker coupling and corresponding looser levels of E-I balance compared to macaque V1 (10). This explanation is likely to be at least partly true, as evidenced by stronger reshuffing in macaque recordings even without a visual stimulus than in mice under any stimulus conditions. However, our ring model suggests an additional factor, namely the featurespecificity of rate reshuffing : rate reshuffing is strongest in matched cells. Averaging across all cells would result in a positive mean change, as observed with optogenetic excitation of E cells in both mice (49) and primates (50). Furthermore, we predict that manipulations that narrow tuning widths will yield increasingly suppressed matched responses compared to the response averaged across all cells.

Lastly, our findings suggest a possible relationship between rate reshuffing and visual feature encoding. A previous study found that, for higher mean luminance levels, increasing grating luminance induced a form of rate reshuffing in matched cells: these cells exhibited heterogeneous individual responses but little or no mean response to increased luminance (5). If luminance changes were transmitted to V1 as an untuned input (an additive change in responses), then our results would suggest that a mean response to luminance changes would be carried by the stimulus-unmatched cells, while the matched cells have a mean response that depends on contrast but is invariant to luminance changes.

### Comparison with results from targeted optogenetic studies

Reducing the dynamics to an effective model of the matched and baseline response populations is straightforward for nonspecific, untuned optogenetic perturbations. However, translating specific, tuned perturbations to the effective model, such as those induced by perturbation of co-tuned cells in recent holographic experiments (51), is challenging.

Chettih and Harvey (52) found, considering cells outside of the stimulation site (further than 25 microns away), that single-cell optogenetic stimulation in mouse V1 during presentation of a visual stimulus most strongly suppresses cells with feature tuning matching the stimulated neuron. In apparent contradiction, Oldenburg et al. (51) found that stimulating iso-tuned ensembles of cells in the absence of a visual stimulus produces the opposite effect. A recent theoretical study (53) resolved this apparent inconsistency by showing that, outside of the stimulation site, networks with feature-selective, like-to-like *I* → *E* and *E* → *I* connectivity preferentially suppress oppositely tuned cells for weak enough neuronal gains, but preferentially suppress similarly tuned cells for stronger neuronal gains. The visual stimulus shown in (52), but not in (51), would increase neuronal firing rates and thus increase neuronal gains, given a supralinear activation function.

We hypothesize that these findings would also naturally result from our linear response analysis applied to a twodimensional feature-dependent model representing one spatial dimension and one orientation dimension. In that case, one could create an effective network by grouping cells into coarsegrained bins of equal width across the two dimensions, rather than grouping by cells that fire like the matched or baseline activity. If these bins are su#ciently wide, then applying our linear response analysis to the course-grained effective network results in a convergent series expansion in powers of the cross-partition projections, as in Eq. (5). Bins su#ciently far from the stimulation site receive no direct optogenetic drive and thus are driven by cross-site interactions. Bins with orientation preferences similar to the optogenetically targeted cells receive stronger projections from the targeted population than orthogonally tuned cells. For stronger or weaker effective coupling (resulting from the presence or absence of a visual stimulus), the cross-site interactions will be suppressive or facilitatory respectively, and so yield stronger suppression of similarly tuned cells or oppositely tuned cells, respectively. This change in regime depends on the network being in the weaker-coupling regime absent a visual stimulus, so a visual stimulus (which, given a supralinear input/output function, increases neuronal gain and thus effective coupling strength, *e.g*. (17)) can then drive the network into the stronger-coupling regime.However, the specific model studied here, fit to data from macaque V1, is always in the stronger-coupling regime, even absent a visual stimulus.

### Connection to Behavioral Modulations

The role of behavioral modulatory signals in sensory cortices, *e.g*. in response to locomotion, is poorly understood. These signals are often unrelated to sensory signals or expectations and thus should be untuned. For example, although the effects of arousal and locomotion are distinct, both are evoked by running (54) and both drive increases in release of both acetylcholine and noradrenaline (2–4, 55), which should evoke inputs that are untuned, though cell-type-specific (3, 56). Thus, our analyses of the effects of untuned inputs on sensory responses may give insights into how neuromodulation affects sensory encoding.

Behavioral modulations show signs of reshuffing. Though the mean firing rate modulation induced by locomotory signals in mouse V1 is not small, individual putative excitatory cell modulations are often larger than the mean response and display mixed activation and suppression (57). Strikingly, the mean locomotor response is suppressed – slightly negative – in marmoset foveal (58), though not peripheral (59), V1, despite large individual cell responses, reminiscent of the optogenetically-induced reshuffing found in macaque foveal V1 (10). The difference of marmoset foveal V1 from mouse or marmoset peripheral V1 could be due to differences in the strength of coupling, as reshuffing requires su#ciently strong coupling (10), or to other differences in neural modulation or connectivity. Arousal has also been shown to elicit effects similar to reshuffing (60).

Our model suggests that untuned optogenetic stimulation does not strongly affect individual cells’ orientation selectivity and tuning widths. Similarly, although running increases the gain of visual responses in mouse V1, it does not affect tuning widths or other measures of tuning (2, 57, 61).

Thus, the mechanisms uncovered here may help explain the mix of response increases and decreases evoked by behavioral perturbations, which in some cases involves little change in mean response as in reshuffing. More insight into these behavioral modulations might be gained by extending our model to include further cell types, such as VIP-, parvalbumin-, and somatostatin-expressing interneurons (62).

## Materials and Methods

### Model details

The model is an *L* = 180 ^°^ ring with *N*_*E*_ excitatory (E) and *N*_*I*_ inhibitory (I) neurons at each of *N* _*θ*_ evenly-spaced sites. Site locations are 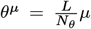 for *µ* = 1,…, *N* _*θ*_ (Fig. 1A). The connection probability from a cell of type *B ∈ {E, I}* at *θ*^*ν*^to any other cell at *θ*^*µ*^ is a Gaussian function of their difference in location:

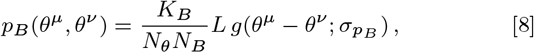

where 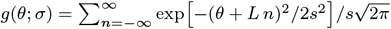 is the probability density function of a wrapped normal distribution. In Eq. (8), 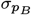 is the connection probability width and *K*_*B*_ the mean in-degree of connections from cell type *B*. The synaptic efficacies per connection type *J*_*AB*_ are homogeneous and are parameterized as *J*_*EE*_ = *J, J*_*EI*_ = −*Jg*_*E*_, *J*_*IE*_ = *J/*β*, J*_*I*_ *I* = −*Jg*_*I*_ */β*.

For a single visual stimulus of orientation *θ*^*m*^ and contrast *c*, the mean afferent input to cells of type *A* at location *θ*^*µ*^ is:

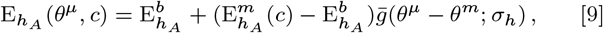

where 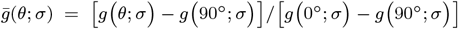 is the baseline-subtracted and normalized wrapped-Gaussian function, 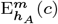 and 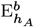 are the mean matched and baseline values, respectively, of the inputs, and *σ*_*h*_ is their width. For a two-grating stimulus, we add another equal-amplitude Gaussian, offset by Δ*θ*^*m*^ = 45^°^. The input to the *i*-th cell of type *A* with preferred orientation *θ*^*µ*^ is 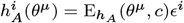, where 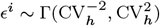 and CV_*h*_ is the coefficient of variation for the afferent inputs.

The baseline and matched mean visual afferent, or external (*X*), inputs are parameterized as:

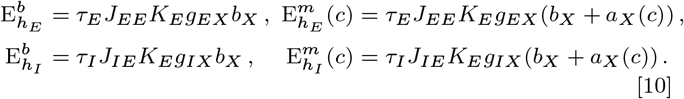

Here *τ*_*A*_ is the membrane time-constant of cell-type *A*, and *b*_*X*_ is the baseline and *a*_*X*_ (*c*) the matched afferent firing rate at contrast *c*. Optogenetic currents to E cells, 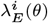, are sampled from a lognormal distribution parameterized by the mean optogenetic input *L* and the coefficient of variation of the optogenetic input CV_*L*_.

The model follows 1^*st*^-order rate dynamics as in (10). Rate model dynamics, numerical simulation details, and network fitting procedure are presented in full in App. A. Briefly, the synaptic weights are *J*_*AB*_ × Bernoulli distributed random variables, 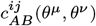, equal to 1 with probability 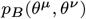 and 0 otherwise. The input-output function is the Ricciardi activation function with parameters of (10). We numerically integrate the rate dynamics using a Dormand-Prince method of order 5(4) and average the activity over the final second of simulation time to compute the response statistics, which we compare to the data of (18). From our data analysis, we compute the value and standard errors for five response statistics at all visual contrasts: The mean and standard deviation of matched activity with and without optogenetic stimulation and the standard deviation of the optogenetically-induced responses. Given a set of model parameters Θ other than the visual input parameters, the associated loss value is the mean square error between the model output and the observed values of the response statistics weighted by the standard errors, which is minimized with respect to the visual input parameters. To fit the model, we first sample values for Θ uniformly within the intervals listed in Table S2. We then sample new values for Θ by perturbing model parameters from the initial sample set that produce the lowest loss. We then chose the best-fit model from the combined set of initial and perturbed parameter samples as the model that minimizes the loss function.

### Dynamical mean-field theory analysis

For cell population *A* at location *θ*^*µ*^ the rate mean at time *t* is 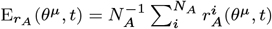 and the rate autocorrelation function between times *t* and *t*^′^ is 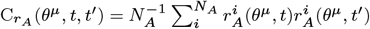. The autocovariance function is 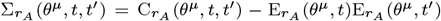. The moments of the activity after optogenetic stimulation 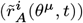 and of the optogenetically-induced response 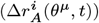 are defined analogously. We study 5 rate moments: the m^*A*^eans and autocorrelations of the rate with or without optogenetic input, and the autocorrelation of the optogenetic response. For each moment at each time or time pair, we approximate its feature-dependence as a baseline-plus-Gaussian. We denote with superscript *µ* = *b, m, a* the values of the rate moments at the baseline (orthogonal to *θ*^*m*^), the stimulus orientation (at *θ*^*m*^), and the auxiliary site (*θ*^*m*^ + Δ*θ*^*a*^), respectively. The baseline-plus-Gaussian feature-dependence approximation allows us to derive the self-consistent equations governing the dynamics of the rate moments at these three sites. Asymptotically, the rate mean becomes stationary and the autocorrelation approaches a function of the absolute time lag.

The self-consistent equations, the numerical solving procedure, and the methods for computing the time-averaged statistics are presented in full in App. B. In summary, we replace the time derivatives in the rate moment dynamics with forward finite differences to derive a naive iterative numerical scheme for evolving the rate moments forward in time from which we can extract the equilibrium rate moments after a su”ciently long time evolution. Alg. S1 details pseudocode for the naive iterative scheme that solves for the rate means and autocorrelations, whereas Alg. S2 shows the pseudocode that solves for the autocorrelation of the optogenetically-induced rate changes. We adopt changes to the naive iterative scheme introduced in (10) to solve the rate moments more e”ciently. Finally, the time-averaged response means equal the rate means their variances equals an appropriate weighted average over the time-lags of the corresponding autocovariance function. We compute the mean balance indices through Monte-Carlo simulation of the net inputs using the mean-field theory inferred distributions.

### Effective Two-Site Model and Linear Response Analysis

If the rate moment tuning widths are stimulus-invariant, fixing them reduces the number of self-consistent equations to just those for the matched and the baseline sites per rate moment per cell type. Suppose we introduce small changes in the statistics of the afferent inputs. To derive the linear response induced by these changes, we differentiate the self-consistent equations for the equilibrium rate moments (see Eq. (3)). Denoting the vector of rate moment linear responses across cell types per site *µ* as 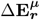 and 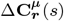, and introducing a quantity we call the rate moment Jacobian 𝒥 ^*µθ*^ (*s, s*^′^), the linear response can be written as

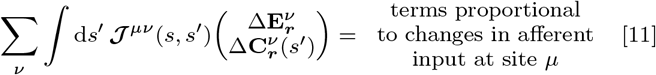

Inverting the Jacobian while treating cross-site projections as perturbative allows us to relate the linear response and the decoupled response as a series expansion in powers of effective cross-site interaction matrices 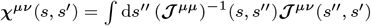, whose sub-blocks we donte as

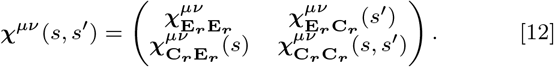

Denoting the decoupled response vectors across cell types as 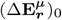 and 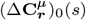, the series expansion for the linear response is then given by

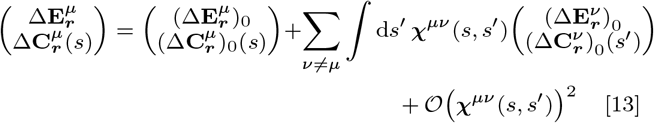

In the mean-interaction approximation, we define 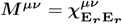. for notational brevity and neglect rate autocorrelation changes.

The expressions for the connectivity of the effective two-site network, the elements of the Jacobian matrix, and the methods for numerically computing the linear response are presented in full in App. C. In short, we decouple the effective two-site model by freezing the visually-evoked cross-site recurrent input to their pre-perturbation levels and calculate the optogenetically-induced decoupled response by solving the decoupled self-consistent meanfield equation equations in the presence of the optogenetic stimulus. Next, we discretize the autocorrelation time lags and treat the linear and decoupled responses as vectors of the concatenation of the mean responses and the discretized autocorrelations at each site. Using the boundary conditions 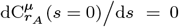 and 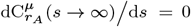, we cast ***𝒥*** ^*µθ*^ (*s, s*^′^) as a matrix that we invert numerically to compute ***χ*** ^*µθ*^ (*s, s*^′^). Finally, we use Eq. (5) or Eq. (13) to compute the linear response.

## Data, Materials, and Software Availability

Code and data for reproducing figures is available at https://github.com/thn2112columbia/Interaction_Untuned_Tuned.

## ACKNOWLEDGMENTS

This work is supported by (support is for work at Columbia except work at UCL indicated by “to A.P.”) the Gatsby Charitable Foundation (GAT3708; and GAT3850 to A.P.), the Kavli Foundation, the NSF (DBI-1707398), the NIH (U01 NS108683, R01 EY029999, U19 NS107613, and T32NS064929), and the Simons Foundation (1156607, to A.P.).

## Supporting Information Text

**Table S1.**
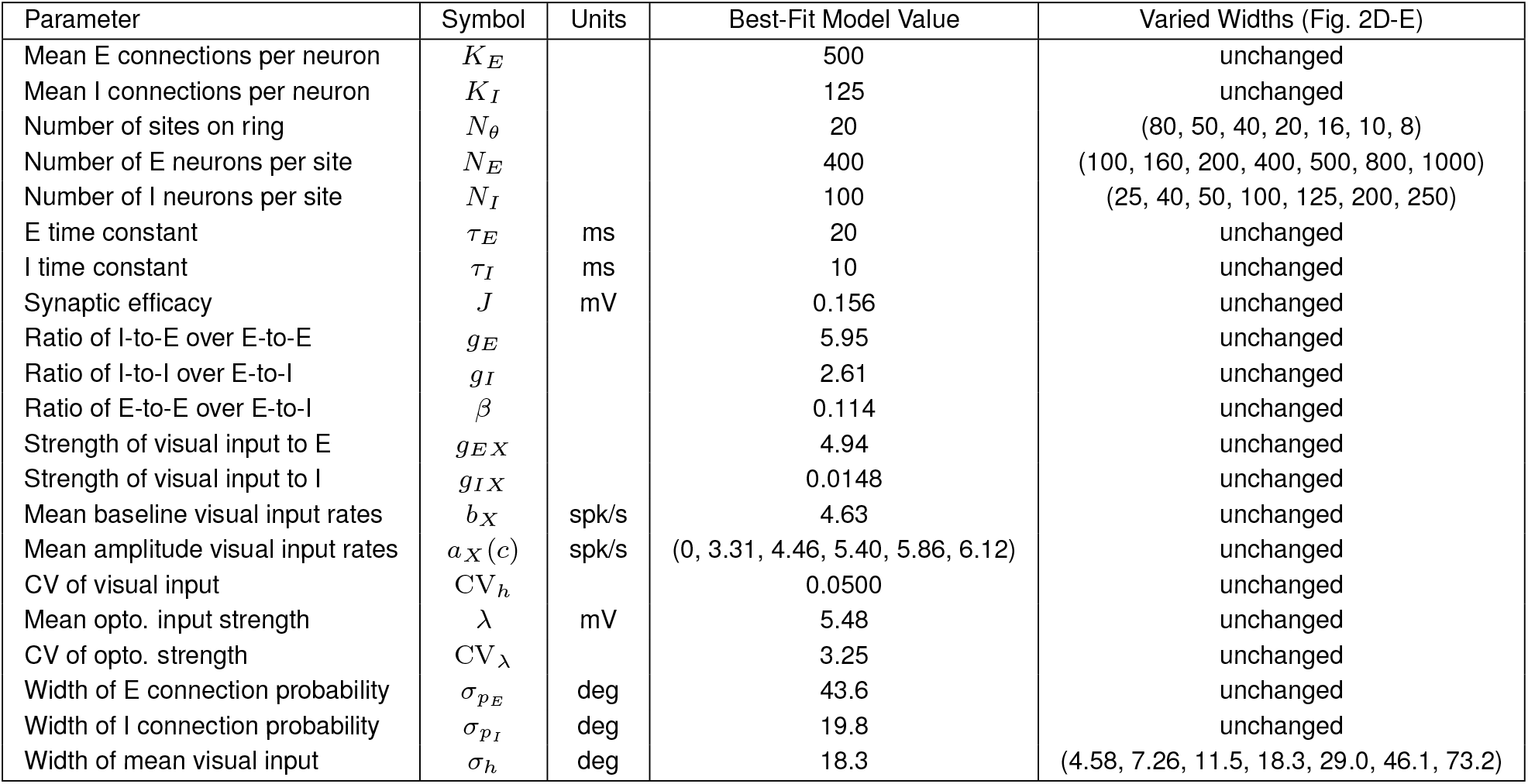
List of network parameters for best fit random ring model for Figures 3-6.

| Parameter | Symbol | Units | Best-Fit Model Value | Varied Widths (Fig. 2D-E) |
| --- | --- | --- | --- | --- |
| Mean E connections per neuron | $K_E$ | | 500 | unchanged |
| Mean I connections per neuron | $K_I$ | | 125 | unchanged |
| Number of sites on ring | $N_\theta$ | | 20 | (80, 50, 40, 20, 16, 10, 8) |
| Number of E neurons per site | $N_E$ | | 400 | (100, 160, 200, 400, 500, 800, 1000) |
| Number of I neurons per site | $N_I$ | | 100 | (25, 40, 50, 100, 125, 200, 250) |
| E time constant | $\tau_E$ | ms | 20 | unchanged |
| I time constant | $\tau_I$ | ms | 10 | unchanged |
| Synaptic efficacy | $J$ | mV | 0.156 | unchanged |
| Ratio of I-to-E over E-to-E | $g_E$ | | 5.95 | unchanged |
| Ratio of I-to-I over E-to-I | $g_I$ | | 2.61 | unchanged |
| Ratio of E-to-E over E-to-I | $\beta$ | | 0.114 | unchanged |
| Strength of visual input to E | $g_{EX}$ | | 4.94 | unchanged |
| Strength of visual input to I | $g_{IX}$ | | 0.0148 | unchanged |
| Mean baseline visual input rates | $b_X$ | spk/s | 4.63 | unchanged |
| Mean amplitude visual input rates | $a_X(c)$ | spk/s | (0, 3.31, 4.46, 5.40, 5.86, 6.12) | unchanged |
| CV of visual input | $CV_h$ | | 0.0500 | unchanged |
| Mean opto. input strength | $\lambda$ | mV | 5.48 | unchanged |
| CV of opto. strength | $CV_\lambda$ | | 3.25 | unchanged |
| Width of E connection probability | $\sigma_{p_E}$ | deg | 43.6 | unchanged |
| Width of I connection probability | $\sigma_{p_I}$ | deg | 19.8 | unchanged |
| Width of mean visual input | $\sigma_h$ | deg | 18.3 | (4.58, 7.26, 11.5, 18.3, 29.0, 46.1, 73.2) |

**Table S2.** List of parameter intervals considered for model fitting.

| Parameter | Sample Interval |
| --- | --- |
| $g_E$ | [3,7] |
| $g_I$ | [2, $g_E - 0.5$] |
| $g_{EX}$ | [0.2, 3.5] |
| $g_{IX}$ | [0.01, 1.5] |
| $\log_{10}(\beta)$ | [-1, 0.7] |
| $\log_{10}(J)$ | [-4, -3] |
| $L$ | [0.2, 7.8] |
| $\log_{10}(CV_L)$ | [-4, -3] |
| $\sigma_{p_E}$ | [15, 45] |
| $\sigma_{p_I}$ | [15, 45] |
| $\sigma_h$ | [15, 45] |
| $b_X$ | [1, 19] |
| $a_X$ | [0, 30] |
| $CV_h$ | [0, 0.6] |

**Table S3.** List of parameters and initial conditions used for the dynamical mean-field procedure.

| Parameter | Symbol | Value |
| --- | --- | --- |
| Integration time step | $\Delta t$ | 0.002 s |
| Warm up time | $T_{warm}$ | 1.2 s |
| Saving time | $s_{sav}$ | 0.4 s |
| Distance from auxiliary to visual-stimulus-matched site | $\Delta\theta^a$ | 15° |
| Initial mean baseline rate | $E_{r_A}^b(t_0)$ | 1 |
| Initial mean auxiliary rate | $E_{r_A}^a(t_0)$ | 2 |
| Initial mean visual-stimulus-matched rate | $E_{r_A}^m(t_0)$ | 4 |
| Initial baseline rate autocorrelation function | $C_{r_A}^b(t_0, t_i)$ | 100 |
| Initial auxiliary rate autocorrelation function | $C_{r_A}^a(t_0, t_i)$ | 400 |
| Initial visual-stimulus-matched rate autocorrelation function | $C_{r_A}^m(t_0, t_i)$ | 2500 |
| Initial baseline opto response autocorrelation function | $C_{\Delta r_A}^b(t_0, t_i)$ | $(\Delta E_{r_A}^b)^2 + 1000$ |
| Initial auxiliary opto response autocorrelation function | $C_{\Delta r_A}^a(t_0, t_i)$ | $(\Delta E_{r_A}^a)^2 + 4000$ |
| Initial visual-stimulus-matched opto response autocorrelation function | $C_{\Delta r_A}^m(t_0, t_i)$ | $(\Delta E_{r_A}^m)^2 + 25000$ |

### Algorithm 1

Naive DMFT algorithm for calculating rate mean and autocorrelation function with baseline-plus-Gaussian profiles

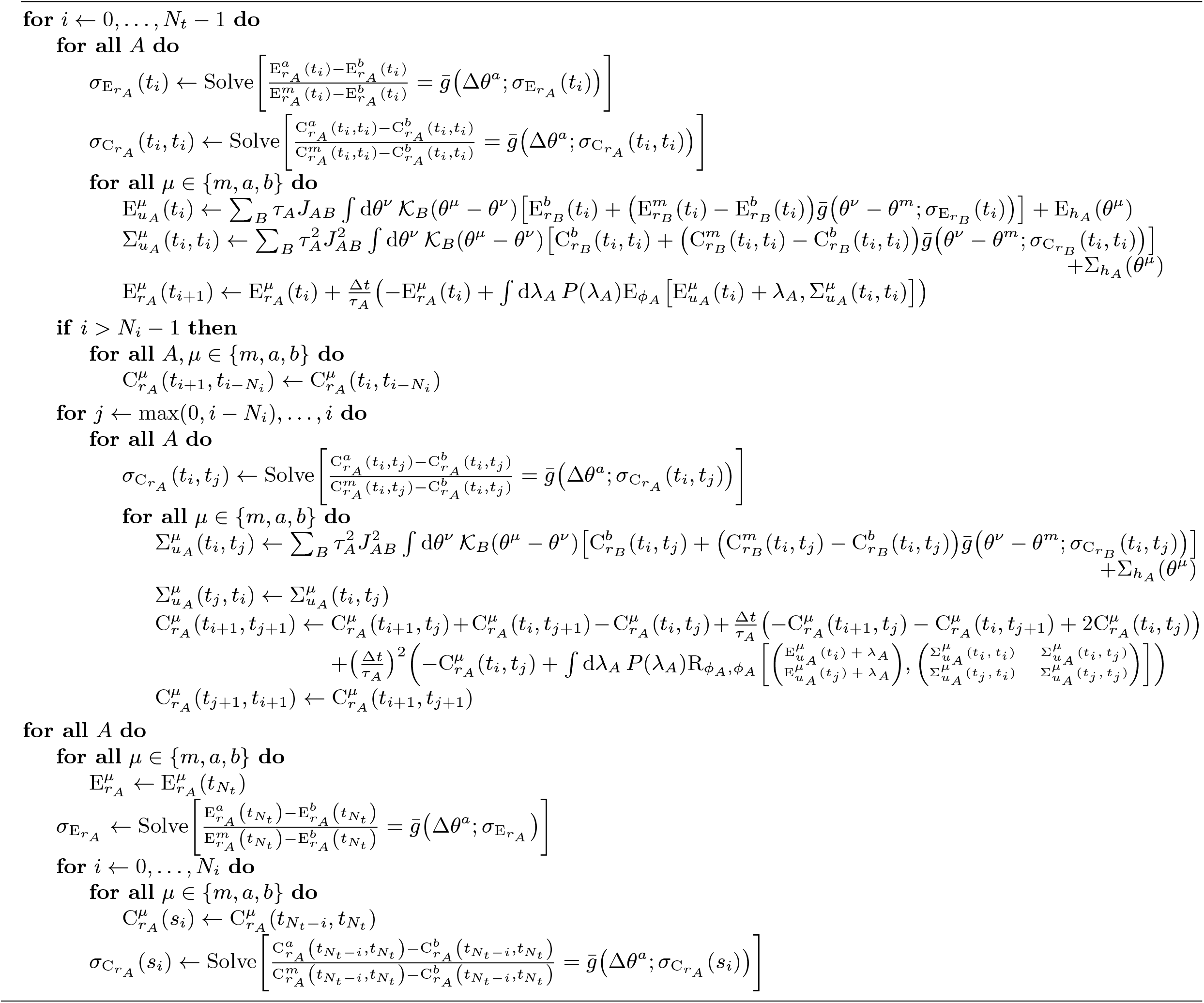

### Algorithm 2

Naive DMFT algorithm for calculating the optogenetic response autocorrelation function with baseline-plus-

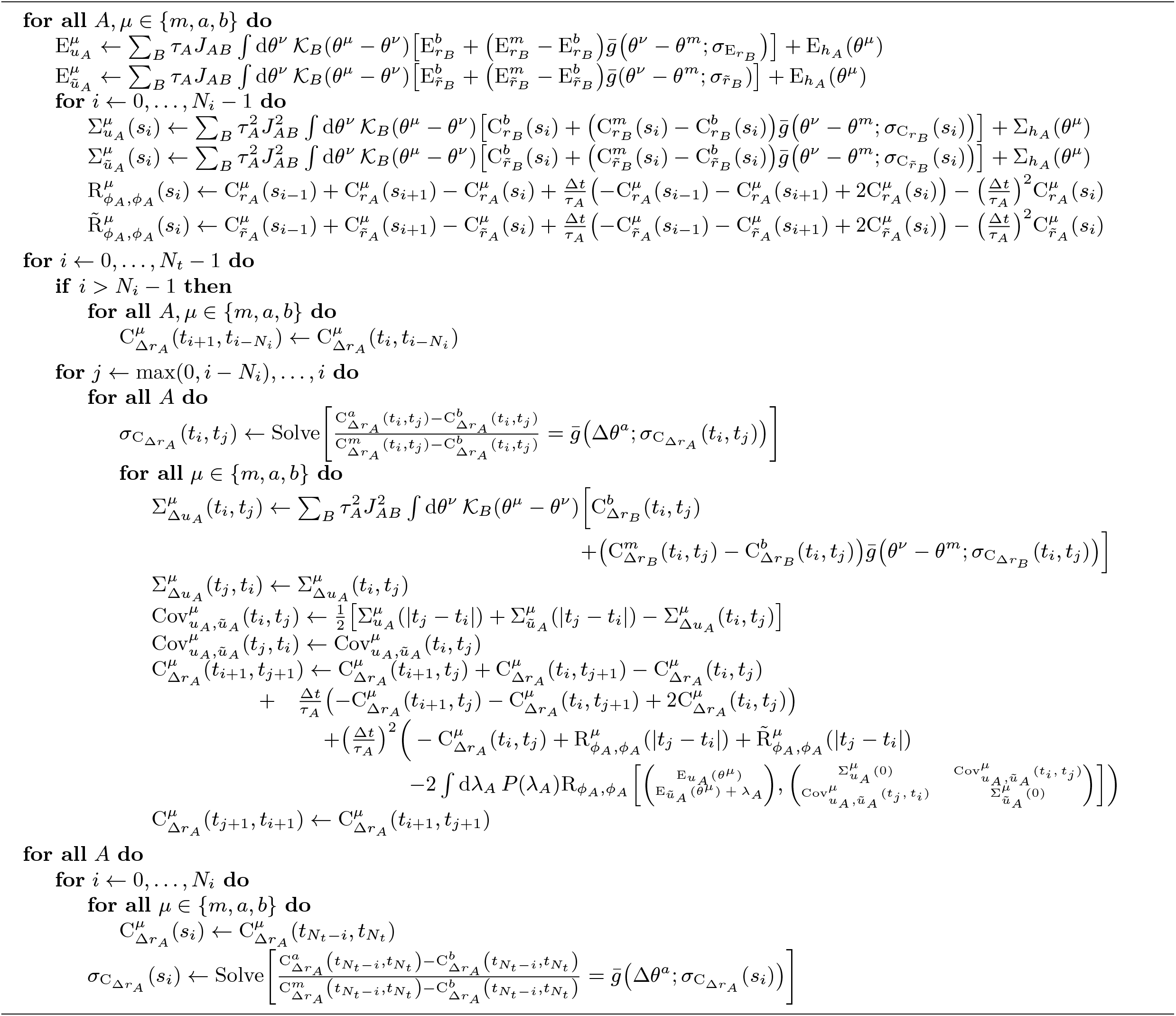

**Fig. S1.**
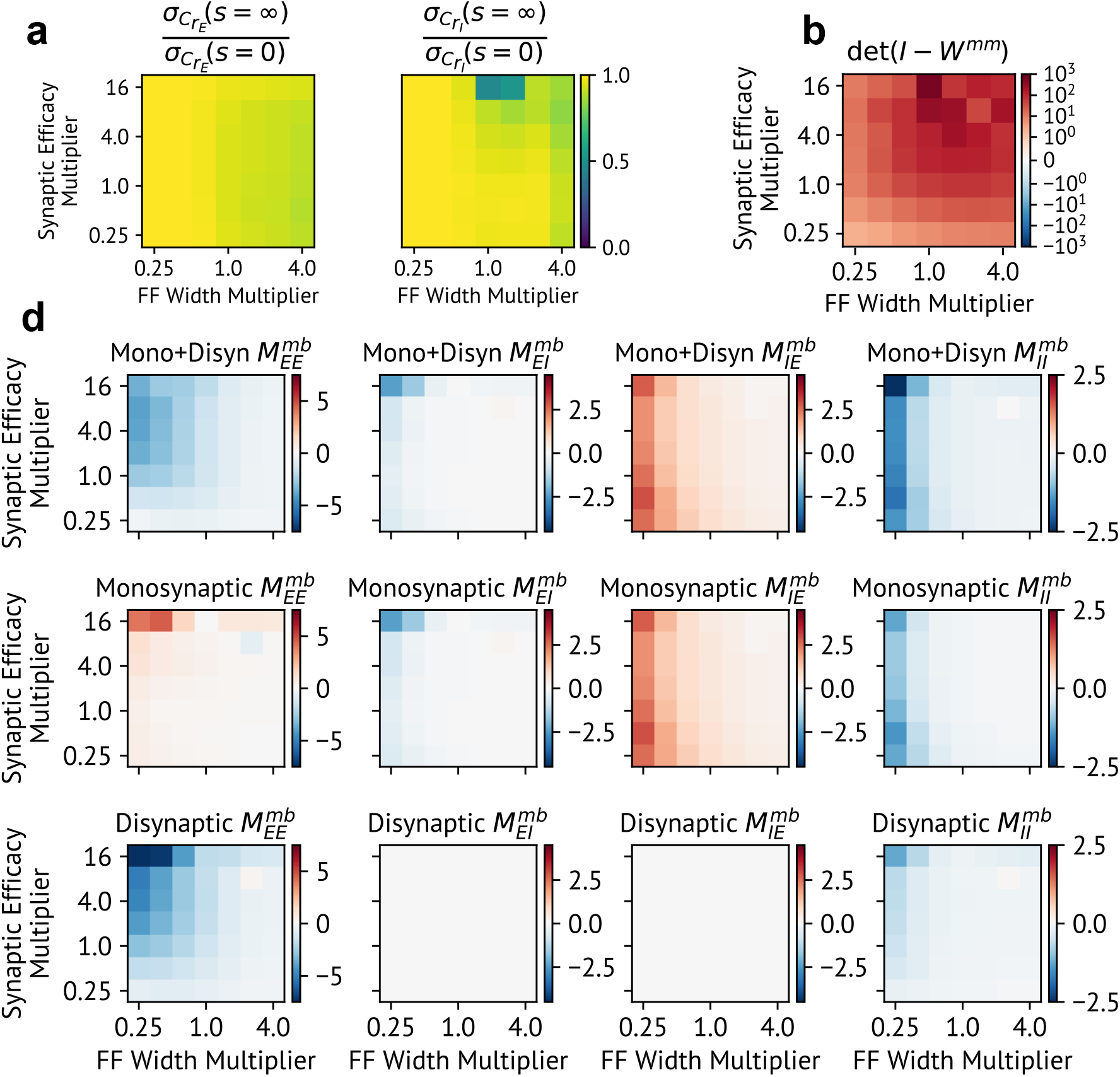
The time-lag-dependence of the rate autocorrelation tuning width is generally weak, the decoupled matched network is stable, and the effective baseline-to-matched interaction matrix features *E* →*E* suppression due to the disynaptic contribution. **a** Except at the highest coupling considered, the rate autocorrelation tuning width at infinite time lags is generally close in value to the tuning width at zero time time-lag for most values of the network coupling and structure strength, indicating weak time-lag-dependence. The left figure plots the ratio for the excitatory rate autocorrelation while the right figure plots the ratio for the inhibitory one. **b** The determinant in the denominator of Eq. (7) is positive for all values of network coupling and structure strength, verifying that the decoupled matched network is stable in the mean-interaction approximation. **c** Except for one set of network coupling and structure strengths, the monosynaptic component of the baseline-to-matched effective interaction matrix matches the sign of the presynaptic cell type while the disynaptic component is a purely inhibitory diagonal matrix whose *E* → *E* element is larger in magnitude than the corresponding monosynaptic element.

**Fig. S2.**
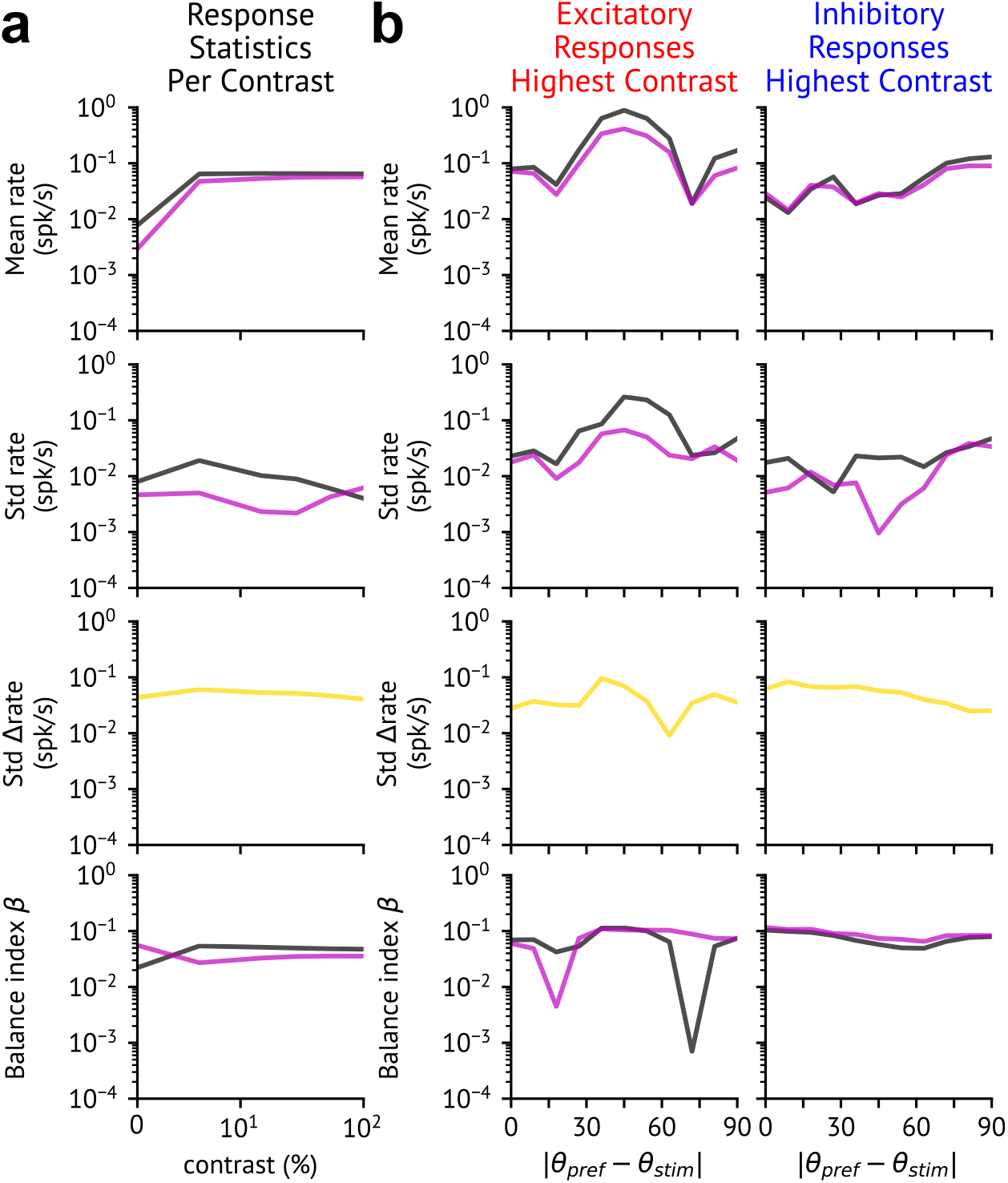
Three-site mean-field solutions for matched response statistics agree within 10% of simulation results. **a**,**b** Solid lines show relative errors of three-site mean-field response statistics compared to simulation results, organized in the same way as Fig. 2a,b. **a** Relative errors between theory and simulations of visual-stimulus-matched response statistics across visual contrast. **b** Relative errors of excitatory (left) and inhibitory (right) populations against absolute distance from preferred orientation to stimulus orientation.

**Fig. S3.**
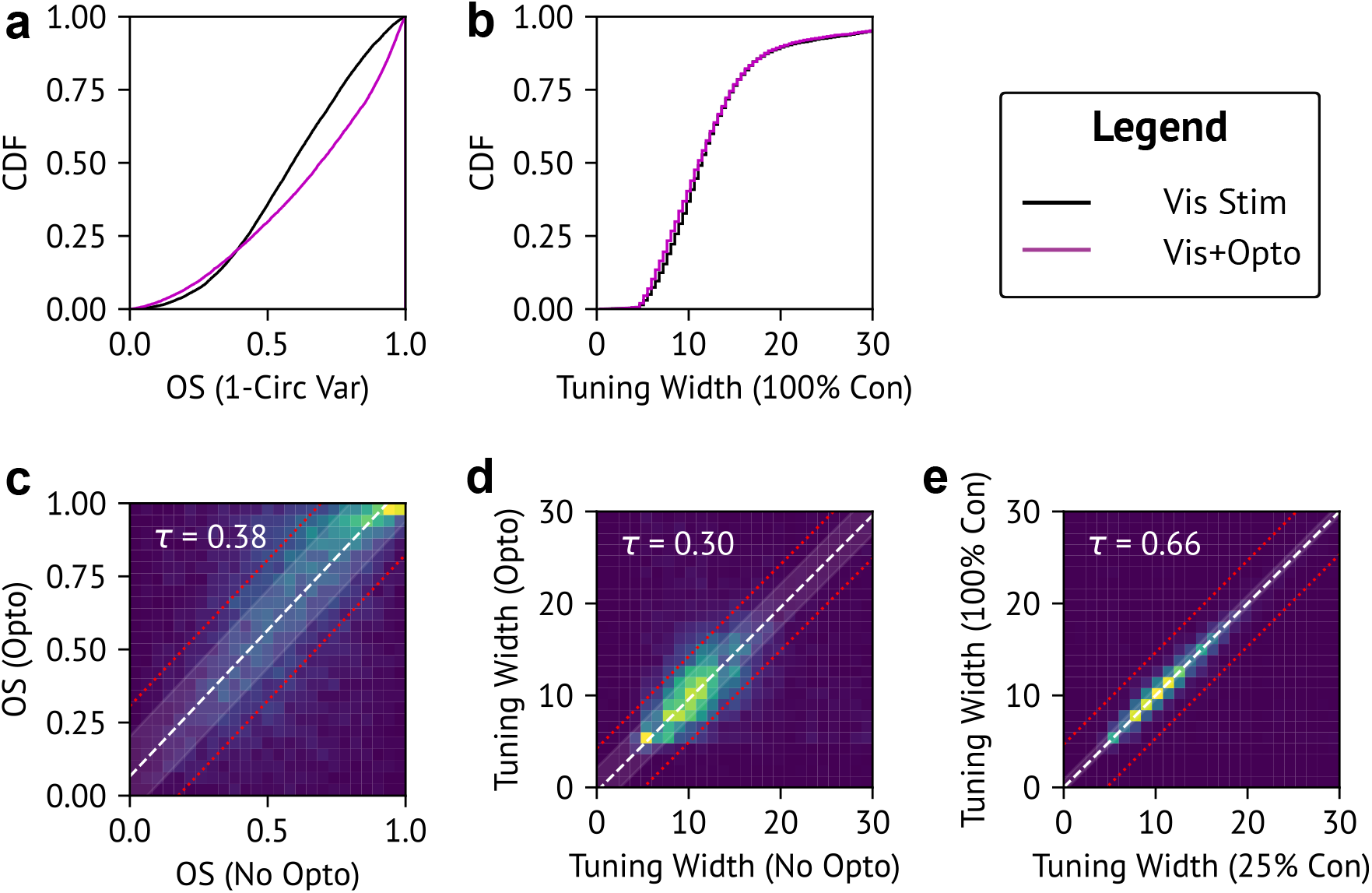
Individual model cells’ tuning curves exhibit contrast invariance and approximately optogenetic-stimulus invariant tuning. **a** The optogenetic stimulus induces a slight rightward shift of the distribution of global orientation selectivities, computed as one minus the circular variance of model cells’ tuning curves at 100% visual stimulus contrast. **b** The optogenetic stimulus only mildly changes the distribution of model cells’ Gaussian tuning widths, which are computed in the same manner as the mean response tuning widths plotted in Fig. 3a. **c**,**d** Besides a mean increase in orientation selectivity, the optogenetic stimulus does not strongly modulate individual cells’ orientation selectivities or tuning widths compared to a random shuffle. Plots show histograms of orientation selectivity (**c**) or tuning width (**d**) at maximum visual contrast without (x-axis) vs with (y-axis) optogenetic stimulation across model cells. White dashed line shows the median change in the plotted quantity across conditions, whereas the white band shows interval between the lower and upper quartiles of changes across conditions. Red dotted lines indicate the interval between the lower and upper quartiles of shuffled data. Kendall tau correlation statistic shown in the top-left of each plot (*p*-value essentially zero) **e** Cells exhibit approximately contrast invariant tuning curves. Plot shows histogram of tuning widths at 25% (x-axis) vs 100% (y-axis) visual contrast.

**Fig. S4.**
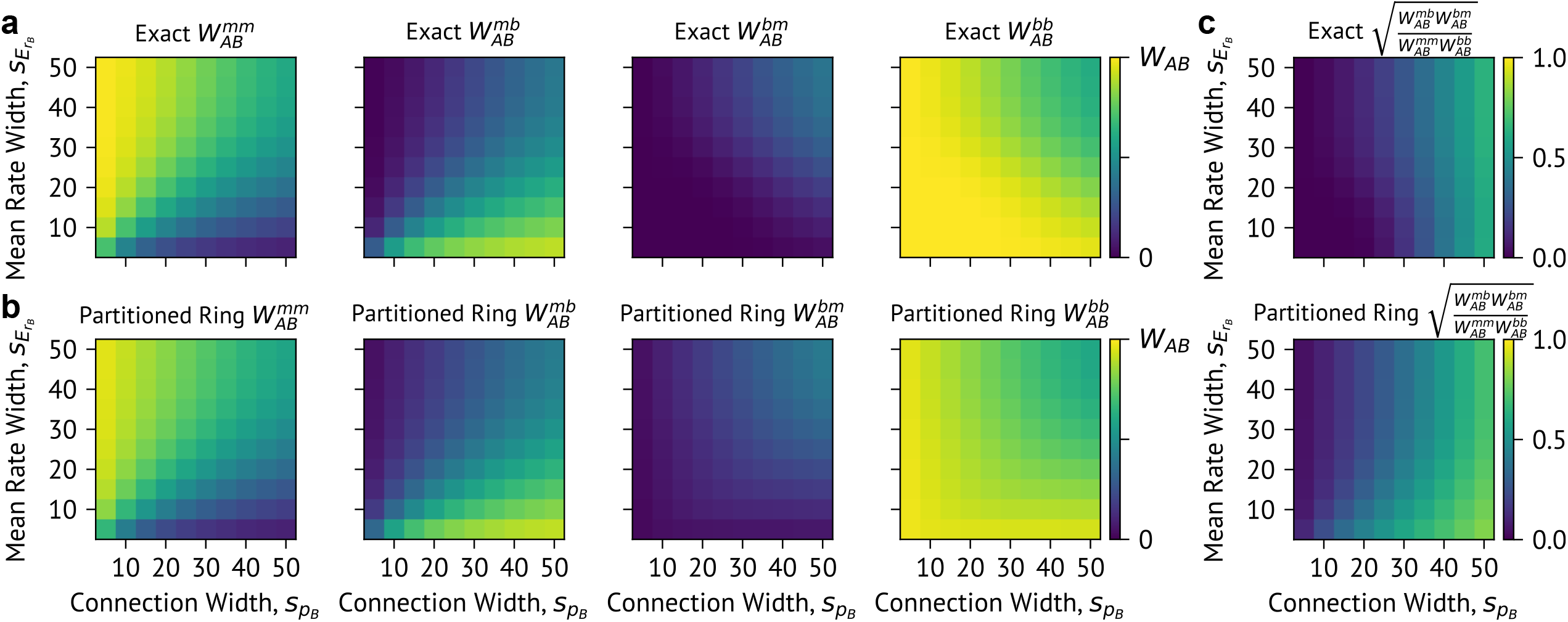
Mean computed from partitioned ring well approximate mean weights from full analysis. **a** Exact mean weights 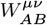 between sites *µ, ν* = {*b, m*} as a function of the connection width 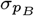 and the mean rate widths 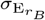 of the presynaptic cell type *B* = {*E, I*}. Analytical expressions for these quantities are written in Eq. (4) of the Main Text. **b** Mean weights computed in the partitioned ring analogy, given by Eq. (S28). **c** The spectral radius of 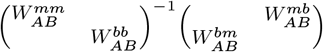 computed exactly and from the partitioned ring analogy is always less than one, indicating that cross-site projections are sufficiently weaker than within-site projections to be treated perturbatively.

**Fig. S5.**
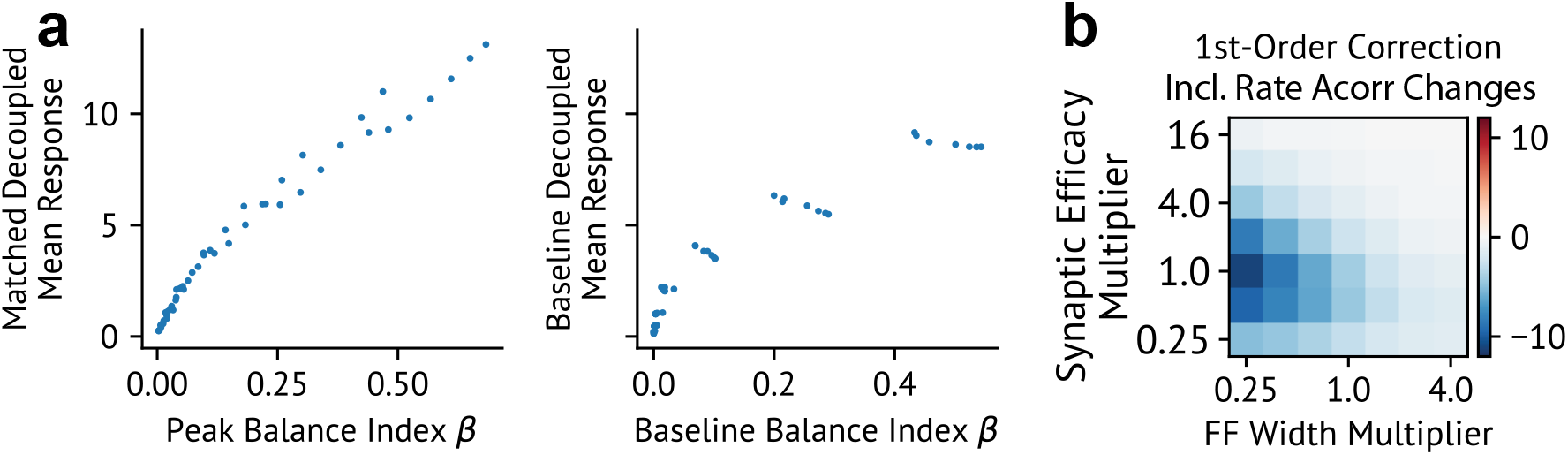
The decoupled responses primarily depend on the level of balance of each effective site, and including rate autocorrelation changes does not significantly change the first-order linear response as we vary the network coupling and structure strength. **a** Decoupled response largely only depends on the effective site’s level of balance. Blue dots show the visual-stimulus-matched (left) or baseline (right) decoupled response as a function of balance index among different models with varied network coupling and structure strengths. **b** The first order correction to the linear response of the matched activity including rate autocorrelation changes nearly matches the correction computed in the mean-interaction approximation (compare with Fig. 4c). Plot shows the correction from the linear response analysis varying feedforward input widths (x-axis) and synaptic efficacy (y-axis).

**Fig. S6.**
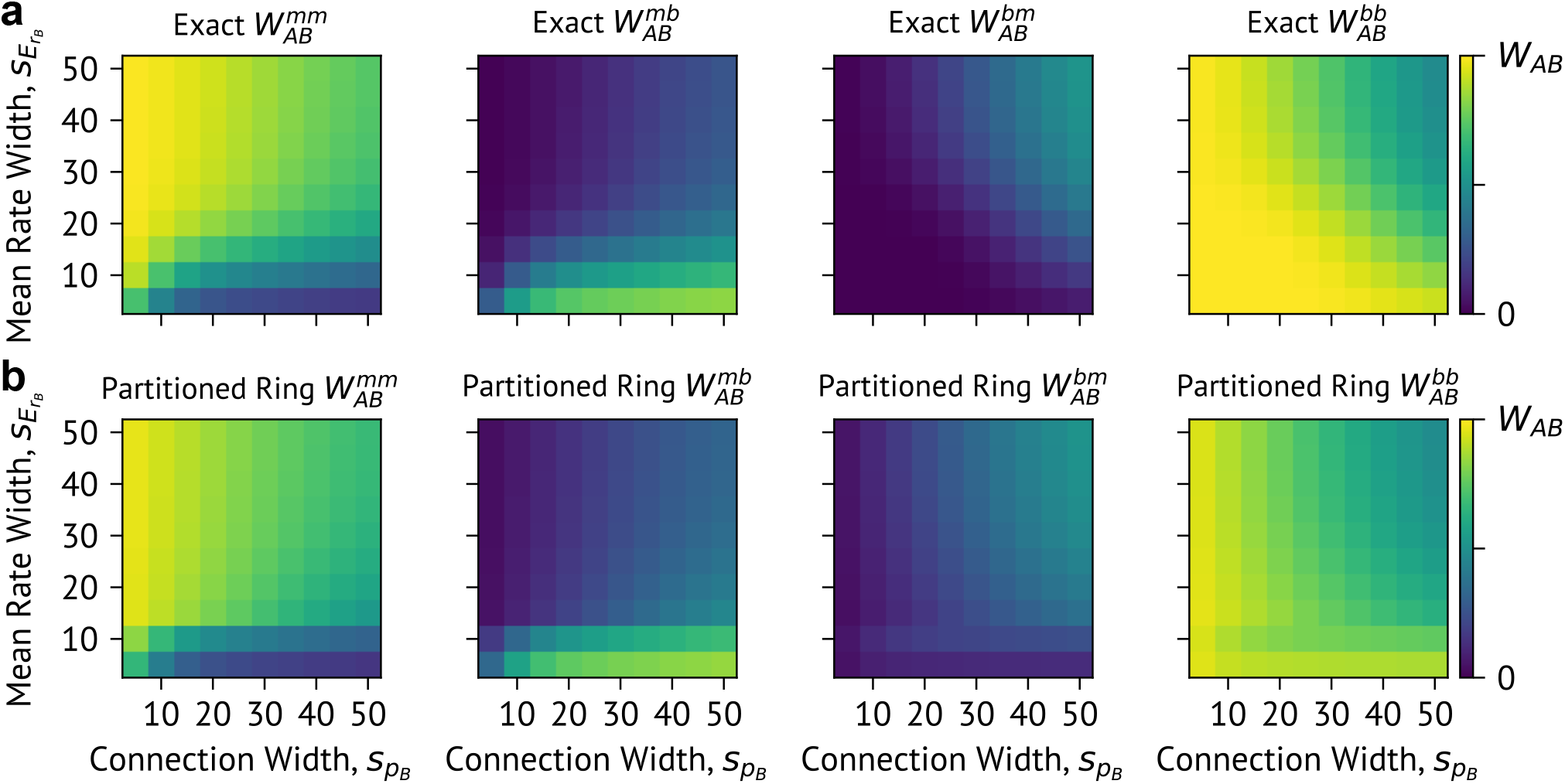
Mean computed from partitioned ring well approximate mean weights from full analysis, even for two grating network. **a** Exact mean weights 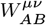 between sites *µ, ν* = {*b, m*} as a function of the connection width 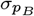 and the mean rate widths 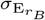 of the presynaptic cell type *B* = {*E, I*}. Analytical expressions for these quantities are written in Eq. (4) of the Main Text. **b** Mean weights computed in the partitioned ring analogy, given by Eq. (S28).

**Fig. S7.**
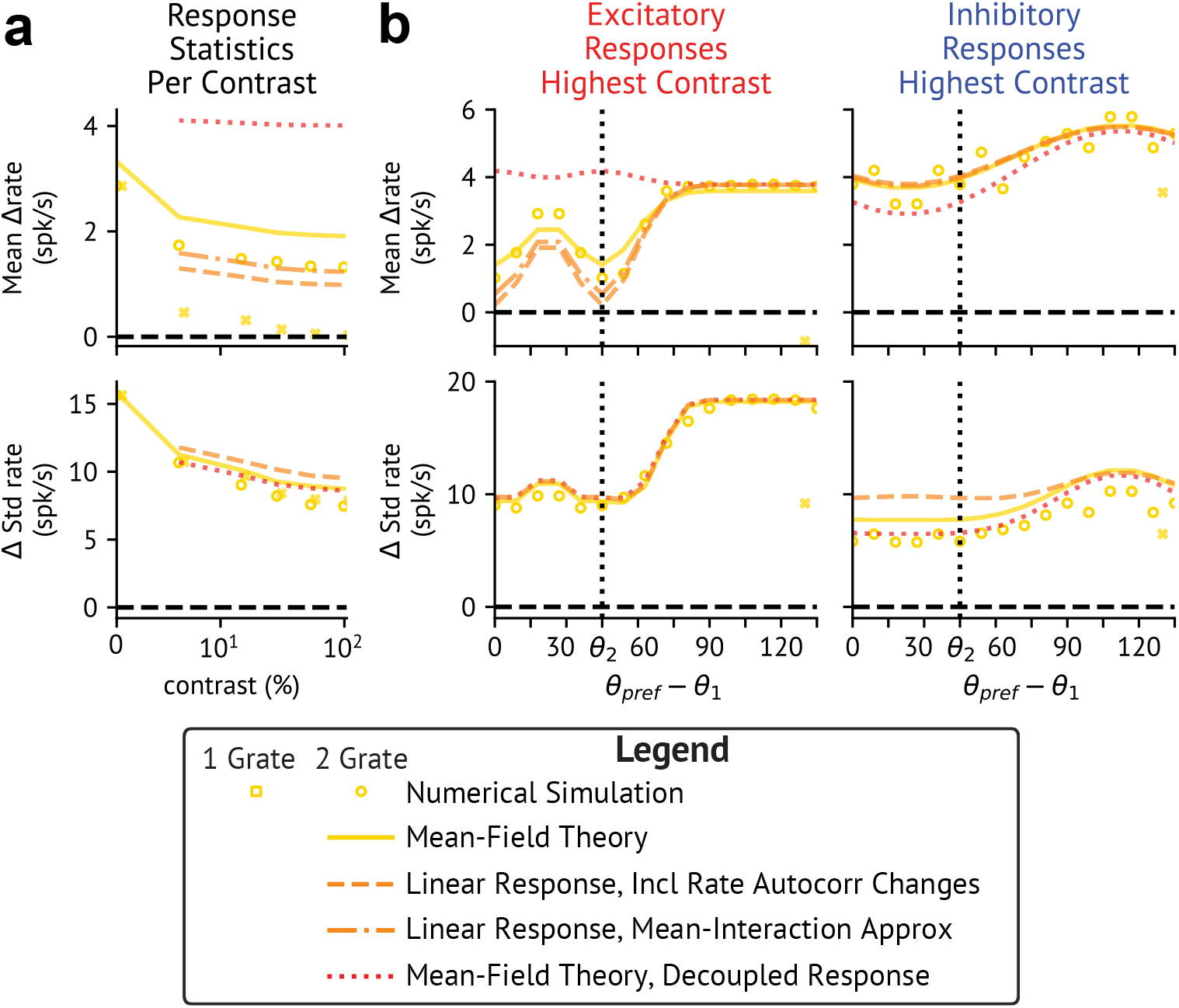
Recomputing linear response with one-grating baseline-to-matched projections produces fully suppressed excitatory matched responses. **a**,**b** Yellow bands, circles, and lines show standard deviation of optogenetically-induced change in rate from the data, simulations, and three-site mean-field theory, respectively. Orange dashed lines and red dotted lines show response statistics computed in the coupled or decoupled effective two-site network, respectively. Figure is organized in the same way as Fig. 4a,b. **a** Visual-stimulus-matched response statistics across visual contrast. **b** Highest contrast response statistic shown for E (left) and I (right) populations versus absolute distance from preferred to stimulus orientation.

## Appendix

### A. Extended Model Details and Numerical Simulation Methods

#### Rate Model Dynamics

We denote 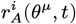 as the firing rate of the *i*-th cell of the population of type *A* at site *θ*^*μ*^. Similarly, we denote 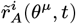 as the firing rate in the presence of the optogenetic stimulus. The firing rates evolve according to

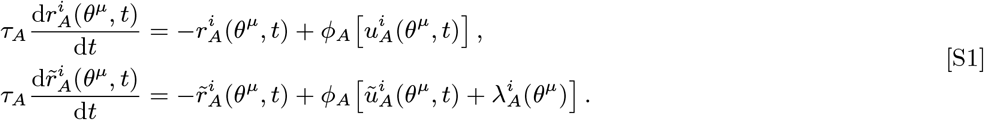

In Eq. (S1), τ_*A*_ is the membrane time constant, 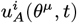 and 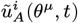 are the net non-optogenetic input currents, 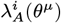 are optogenetic input currents, and *ϕ*_*A*_ is the Ricciardi activation function (1, 2):

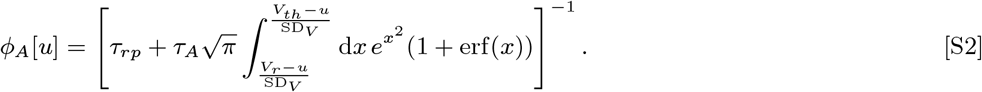

The Ricciardi function parameters are set as τ_*rp*_ = 2 ms for the refractory period, *V*_*th*_ = 20 mV for the threshold potential, *V*_*r*_ = 10 mV for the resting potential, and SD_*V*_ = 10 mV for the amplitude of the membrane potential fluctuations.

The net non-optogenetic inputs are defined as:

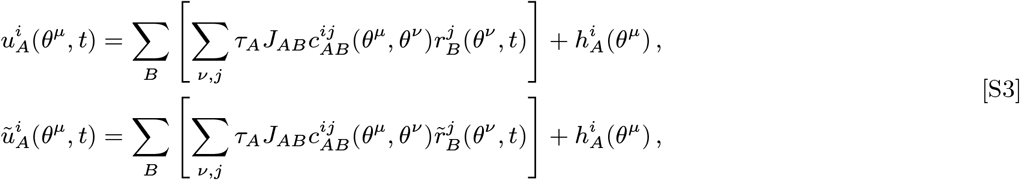

Above we denote *J*_*AB*_ as the synaptic efficacy of the corresponding connection type (see Methods in the Main Text for parameterization) and 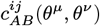 as the adjacency matrix, which is equal to 1 if there is a connection from cell *j* of type *B* at site *θ* ^*ν*^to cell *i* of type *A* at site *θ*^*μ*^, which occurs with probability

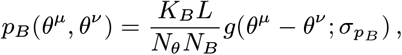

and zero otherwise.

#### Numerical simulation details

Models consist of *N*_*θ*_ = 20 sites on the ring, with *N*_*E*_ = 400 E cells and *N*_*I*_ = 100 I cells for a total of 10, 000 neurons on the ring or 500 neurons per site, except where otherwise noted. Networks are initialized with zero firing rates, and Eq. (S1) is numerically integrated using the Dormand-Prince 5(4)-order adaptive step integration scheme (3) for 5 s. We record each neuron’s activity and separate excitatory (including optogenetic currents) and inhibitory net inputs every Δ*t* = 0.002 s of simulation time. Rates and inputs are time-averaged over the final second of integration and time-averaged quantities are denoted with angle brackets, *i.e*. 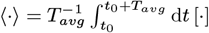. where *t*_0_ = 4 s and *T* = 1 s.

We simulate model dynamics twice, once without optogenetic currents (λ_*A*_ = 0) and once with laser stimulation (λ_*E*_ ≠ 0).

The optogenetic response or the change in activity induced by the laser is defined as

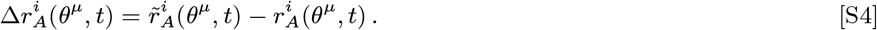

Similarly, we define the change in the non-optogenetic net input as 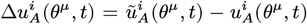. We measure eight statistics per population: the mean rates with 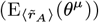 and without optogenetic input 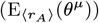, the mean optogenetic response 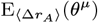, the standard deviation of the rates with 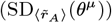 and without 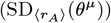 optogenetics, the standard deviation of the optogenetic response 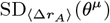, and the mean balance index *β* with and without optogenetic input. The balance index for a single cell is defined as the ratio of a cell’s rectified total input to its total excitatory (recurrent plus external) input. In addition to calculating the population statistics per cell type, we also compute statistics averaged over both *E* and *I* cells at each site. These rate statistics are averaged over 100 realizations of the model’s random connectivity and inputs.

We modulate the network’s structure by fixing the mean visual-stimulus-matched afferent input 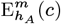 while varying the feedforward input width *s*_*h*_ by a factor α. We want the baseline to receive a similar external input distribution compared to the matched site in the wide width limit to remove as much structure as possible. To do so, we remove the baseline subtraction from Eq. (9) of the Materials and Methods (see Main Text) and parameterize the mean afferent input as

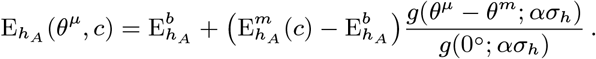

As we vary the widths, we also adjust *N*_*θ*_ to maintain the proportion of visual-stimulus-matched cells approximately proportional to α. When we adjust *N*_*θ*_, we change *N*_*E*_ and *N*_*I*_ such that the total number of E and I cells are equal between models. The exact widths, the corresponding number of sites, and the corresponding number of cells at each site are listed in the rightmost column of Table S1.

To measure tuning widths from numerical simulations, we first apply a smoothing spline (4) with a regularization parameter of 40 to the feature-dependence of the calculated rate moments, extended to the range *θ* ∈ [−90^°^, 180^°^] to enforce the periodic boundary conditions. We baseline-subtract the smoothed profile, defining the baseline as the rate moments at *θ* = 90^°^ when the network is presented with a single visual stimulus. In the one grating network, we calculate the half width at half the height of the baseline-subtracted visual-stimulus-matched rate moment (HWHH) and compute the tuning width σ As 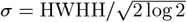. In the two grating network, we use the baseline value used in the subtraction of the one-grating network at the same contrast and optogenetic input strength to perform the two-grating baseline subtraction. If the rate moment profile intersects half the baseline-subtracted matched value four times, then the Gaussian components are non-overlapping and the HWHH of either of the Gaussian components determines the tuning width. If the profile intersects only twice, then we correct for the overlapping Gaussian components by solving for 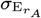 in the equation:

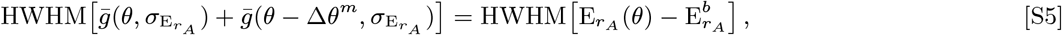

where 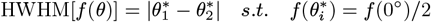 computes the HWHH of the input function.

When computing the tuning widths of individual model cells’ tuning curves, we set the smoothing spline regularization parameter to 100 and extend to the range *θ* ∈ [−180^°^, 360^°^] due to the higher level of noise in individual tuning curves compared to population tuning curves. The baseline of an individual cell’s tuning curve is computed by averaging the response in the range *θ* ∈ [90^°^ 22.5^°^, 90^°^ + 22.5^°^] relative to each cell’s preferred orientation, which we define as the value *θ*_*pref*_ that maximizes the smoothed tuning curves. We note that this value is not necessarily equal to cells’ assigned preferred orientations, but are generally close in value to their assigned orientation preference. We then compute the tuning width σ as 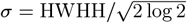 from the HWHH of the baseline-subtracted smoothed tuning curve.

#### Fitting procedure

A set of candidate network parameters Θ related to the network connectivity and the optogenetic input distribution are generated randomly, independently, and uniformly in the intervals listed in Table S2. For each set of parameters above, we simulated the network over a three-dimensional hypercube of afferent input parameters selected over the intervals listed in Table S2 and compute the visual-stimulus-matched model response statistics *Y* (Θ, CV_*h*_, *b*_*X*_, *a*_*X*_) (see Methods in Main Text for definitions of CV_*h*_, *b*_*X*_, and *a*_*X*_).

We analyze the experimental data from (5) and calculate the mean 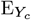 and standard error 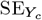 of the quantities 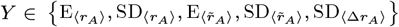 for each contrast *c*. We then define the contrast-specific loss function ℒ_*c*_(Θ, CV_*h*_, *b*_*X*_, *a*_*X*_) as the mean square error between the experimental population response statistics at a given contrast *c* and the predicted model statistics *Y* (Θ, CV_*h*_, *b*_*X*_, *a*_*X*_), weighted by the experimental standard errors. That is, the loss function is given by

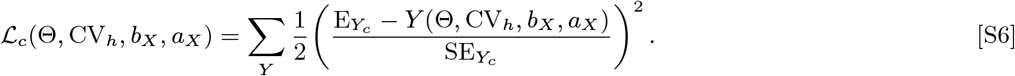

For a given Θ, we select the afferent input parameters that best minimize the contrast-specific loss at all contrasts. That is, we define the network loss function over Θ as

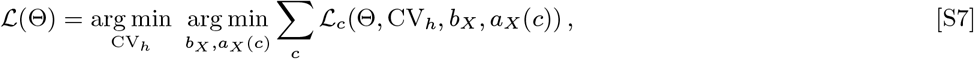

where *a*_*X*_ (*c*) is a strictly increasing mapping between contrasts and values of *a*_*X*_ with *a*_*X*_ (0) = 0.

After first performing a random search over the parameter space, for each of the model parameters yielding the lowest total loss we then produce multiple perturbed parameters to produce a new set of models to simulate. To produce perturbed parameters, we multiply each quantity in Θ by an independent random factor sampled uniformly from the interval [0.9, 1.1]. From the combined set of models from the initial random search and the subsequent perturbed search, we choose the best-fit parameters as those that minimize the loss function.

### B. Extended Dynamical Mean-Field Theory Methods

#### Self-Consistent Equations under Baseline-Plus-Gaussian Feature-Dependence Approximation

We derive the self-consistent equations for an arbitrary structured rate model in App. D. To simplify these self-consistent equations, we assume the means of 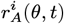 and 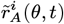 at all times and the autocorrelation functions of 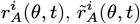, and 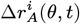 at each time pair have a baseline-plus-Gaussian feature-dependence. That is, we approximate their dependence on *θ* as:

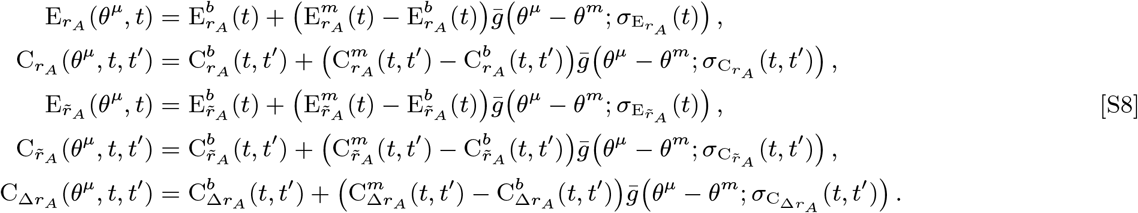

While the tuning widths of the equilibrium rate autocorrelations (*e.g*. 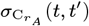) depend on the time lag |*t*^′^ − *t*|, we find that the equilibrium lag-dependence of the rate autocorrelation width is mild (Supp. Fig. 1a). Letting *θ*^*a*^ be the location of the auxiliary site and denoting with superscript *a* the rate moments at *θ*^*a*^, the rate moment tuning widths are computed as the solutions to equations of the following form (illustrated for 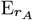, but true for all 5 moments):

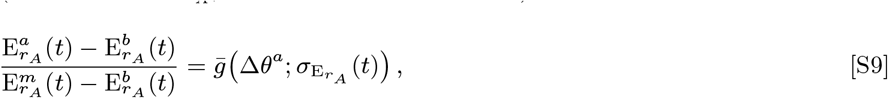

where Δ *θ*^*a*^ is the distance between the stimulus and auxiliary sites, *i.e*. *θ*^*a*^ = *θ*^*m*^ + Δ *θ*^*a*^, and where 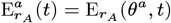.

The correlations between the rates and weights become increasingly negligible in large network limit. Furthermore, in the limit that neurons receive a large number of synapses, by the central limit theorem their recurrent inputs are well approximated as samples from a Gaussian process whose mean and autocovariance function are determined by the rate moments (6). Although the afferent inputs are gamma distributed, they are well-approximated as normally distributed due to rapidly suppressed third and higher-order moments in the limit of small CV_*h*_. Thus, the net non-optogenetic inputs are well approximated as Gaussian.

The rate means determine the mean of the net non-optogenetic inputs 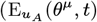 and 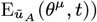, the rate autocorrelation functions determine the autocovariance functions of those inputs 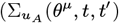 and 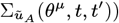, and the optogenetic response autocorrelation determines the autocovariance of the optogenetically induced change in inputs 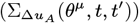. In computing the statistics of the net inputs, one must perform sums of the form 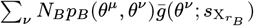, where 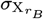 is the tuning width of a rate moment at a given time or time pair (see Eqs. (S41) and (S42) in App. D). In the limit of large *N* _*θ*_ and letting *L*/*N* _*θ*_ = Δ *θ* denote the distance between discrete ring sites, we can convert this sum to a convolution as follows:

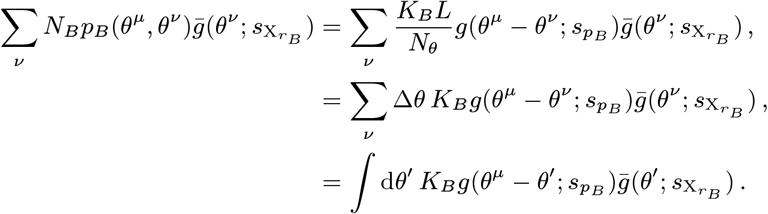

The convolution 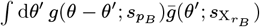 yields a function we denote as ζ, which is given by:

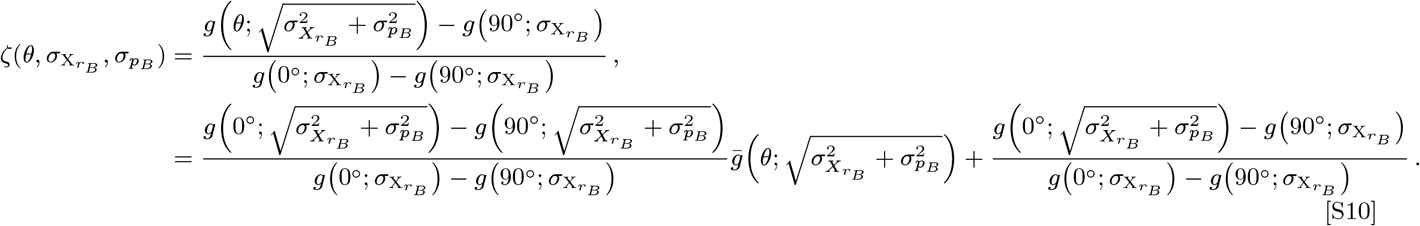

In the second line of Eq. (S10), we see that 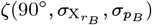 is nonzero, meaning that ζ decays to a nonzero baseline value.

We next write down the dependence of the net input statistics on the rate moments. Before doing so, we note that, in the expressions for the input autocovariances, terms appear involving the convolution of the squared rate means with the weight profiles. We neglect these terms, because they are proportional to the connection probability *p*_*B*_ (*θ* ^*μ*^, *θ* ^*ν*^), which can be neglected if *K*_*B*_ is fixed in the limit that *N*_*B* → ∞_. Empirically, we find that ignoring these terms does not significantly impact the resulting response statistics. Using a superscript *μ* = {*b, m, a*} to denote the net input statistics at the three sites, the statistics of the net inputs are then given by:

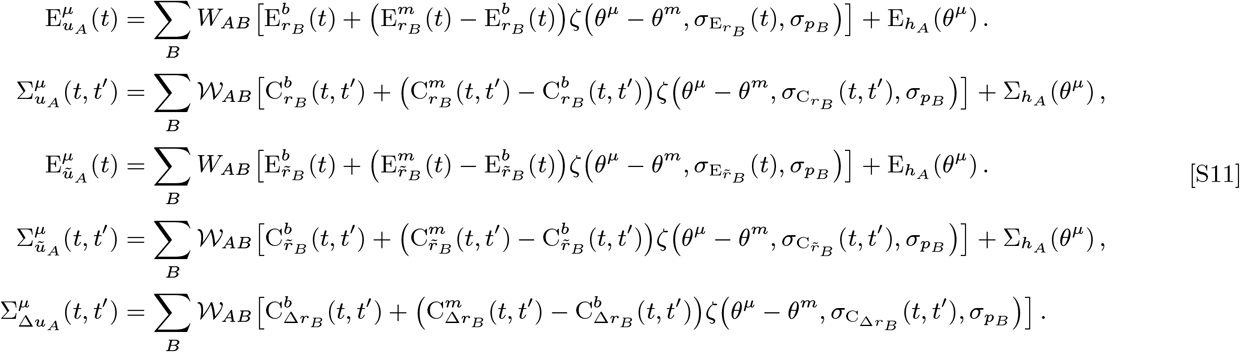

In Eq. (S11), 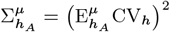 is the variance of the afferent input, whereas *W*_*AB*_ and *𝒲*_*AB*_ denote the mean and variance of the total weights, respectively, which are defined as

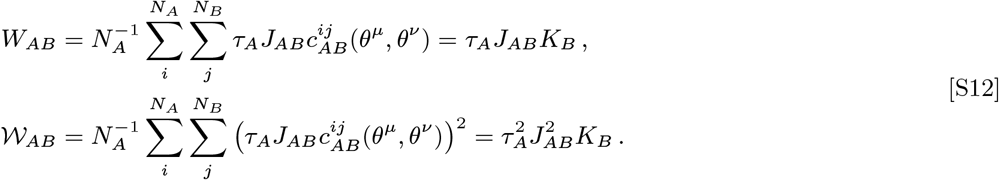

Finally, the rate dynamics induce dynamics on the rate moments. The equations governing the five rate moments of interest are as follows:

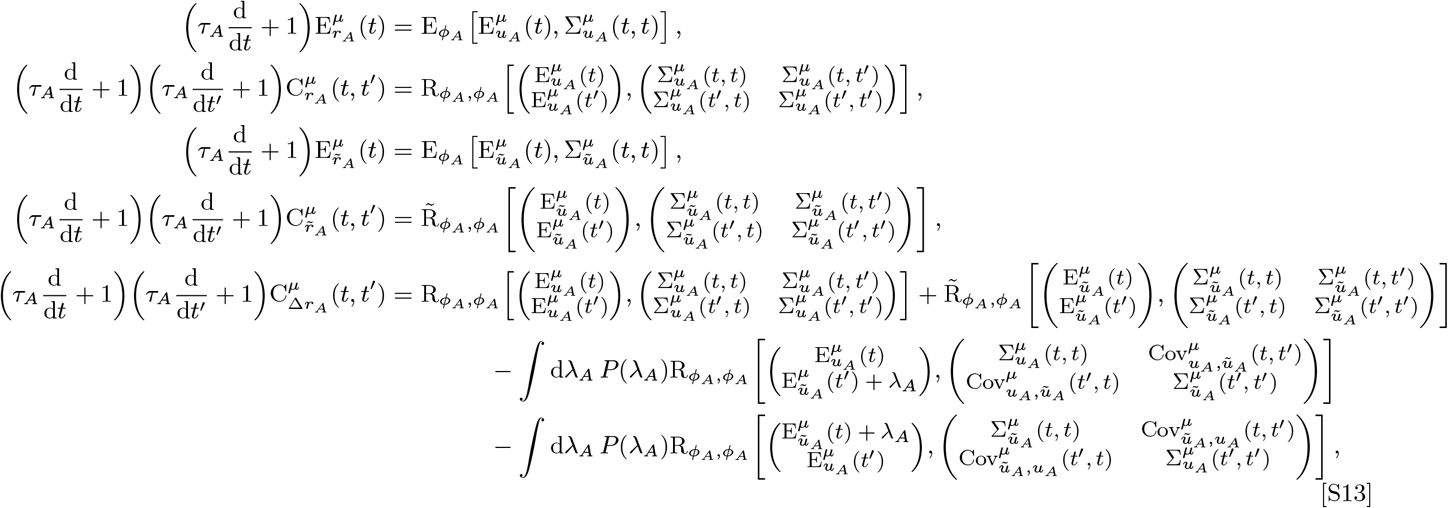

In Eq. (S13), the covariance between the initial and perturbed non-optogenetic net inputs is given by

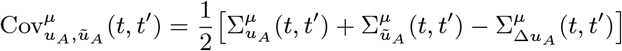

(see App. D for the derivation). In addition, the mean auxiliary functions E_*f*_ [*μ*, ∑] and 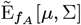 and the cross-correlation auxiliary functions 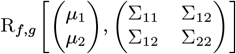 and 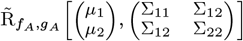 result from computing the mean or the crosscorrelation, respectively, of the subscript functions over Gaussian distributed inputs parameterized by the means and covariance matrices in their arguments. Letting 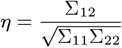, these auxiliary functions are defined as

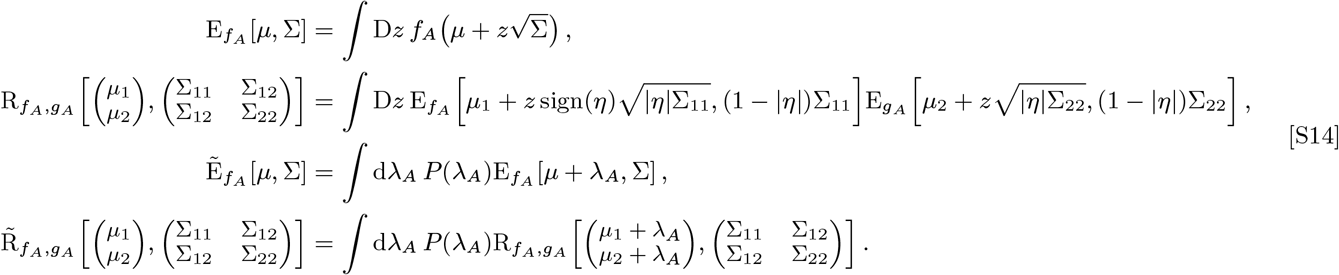

In Eqs. (S13) and (S14), *P*(λ _*A*_) indicates the density of the log-normally distributed optogenetic inputs. We first define the mean 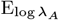 and standard deviation 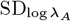 of the log of the optogenetic input λ_*A*_:

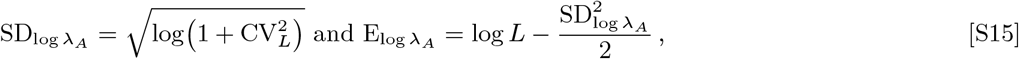

where *L* and CV_*L*_ are the mean and coefficient of variation of λ_*A*_. The distribution *P*(λ_*A*_) is then given by:

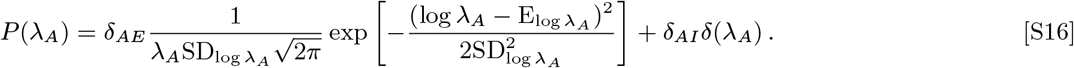

The rate moments reach equilibrium after evolving for a sufficiently long time. That is, the mean rate reaches a stationary value 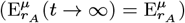 and the autocorrelation function of the rate becomes a stationary function of the difference in the time points 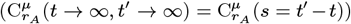. From the equilibrium rate moments, we can then compute the equilibrium net input statistics, such as 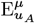 and 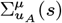. Note that we will denote dynamical times by *t* whereas autocorrelation and autocovariance time lags will be denoted by *s*. Equilibrium rate moments (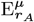 and 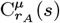) will be written with one fewer temporal argument than dynamical rate moments (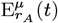 and 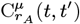) Likewise, we will use similar notation to differentate between equilibrium and dynamic net input statistics.

By replacing the dynamical rate moments in Eq. (S13) with the equilibrium rate moments (and likewise replacing the dynamical net input statistics by their equilibrium counterparts), we derive the self-consistent equations that the equilibrium rate moments must satisfy, which are as follows:

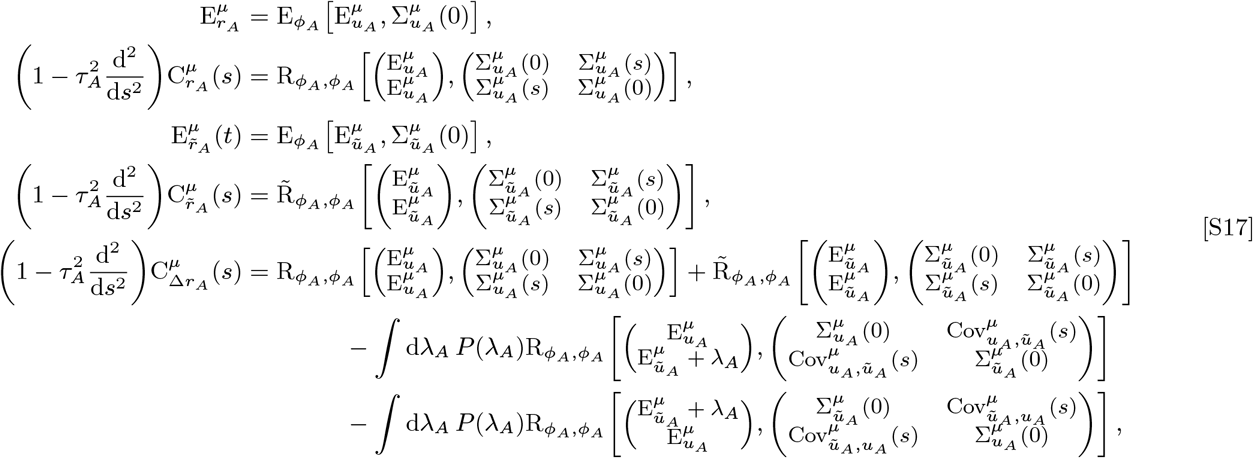

In Eq. (S17), the differential operator 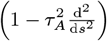 arises due to the fact that 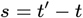, meaning that 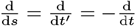, Note that if we write out the Gaussian integrals defining the mean and cross-correlation auxiliary functions in the first two lines of Eq. (S17), we would arrive at expressions equivalent to Eq. (3) of the Main Text.

However, instead of analytically solving for the steady state solutions of Eq. (S17), we can start from any arbitrary stable and valid (non-negative) initial conditions of the rate moments and evolve them using Eq. (S13) until they reach equilibrium to extract the steady-state solutions.

#### Two Equal Contrast Interocular Stimuli

We derive the generalization of our baseline-plus-Gaussian mean-field analysis in App. E. For the ring driven by two equal-contrast gratings differing in orientation by Δ *θ*^*m*^, we add a second equal-amplitude Gaussian at the second stimulus orientation to the mean afferent input. Due to the equal contrasts of the stimuli, the rate moments have identical tuning widths and visual-stimulus-matched values at the two grating orientations. We assume the rate moments have a baseline-plus-Gaussian-mixture feature-dependence, with each Gaussian centered at one of the two grating orientations. Correcting for the overlapping Gaussian components, the rate moments are parameterized as

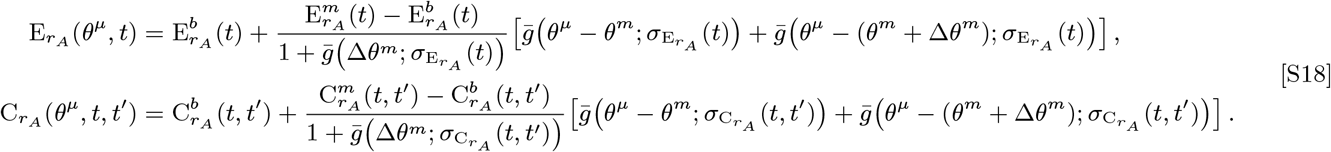

The remaining three rate moments are parameterized in a similar manner.

In the two-grating case, the auxiliary site location could equally be chosen as *θ* ^*a*^ = *θ* ^*m*^ + Δ*θ* ^*m*^ + Δ*θ* ^*a*^ or *θ* ^*m*^ Δ*θ* ^*a*^. The two have identical rate moments. If we choose the former, the tuning widths are implicitly defined by the solution to equations of the following form (illustrated for 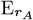, but true for all 5 moments):

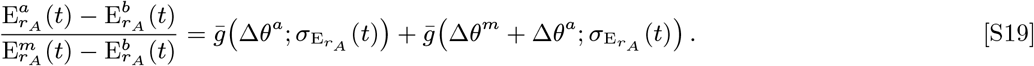

Substituting Eq. (S18) into Eqs. (S11) and performing the convolutions allows us to express the net input statistics at the visual-stimulus-matched and auxiliary sites (*μ* = {*m, a*}) as:

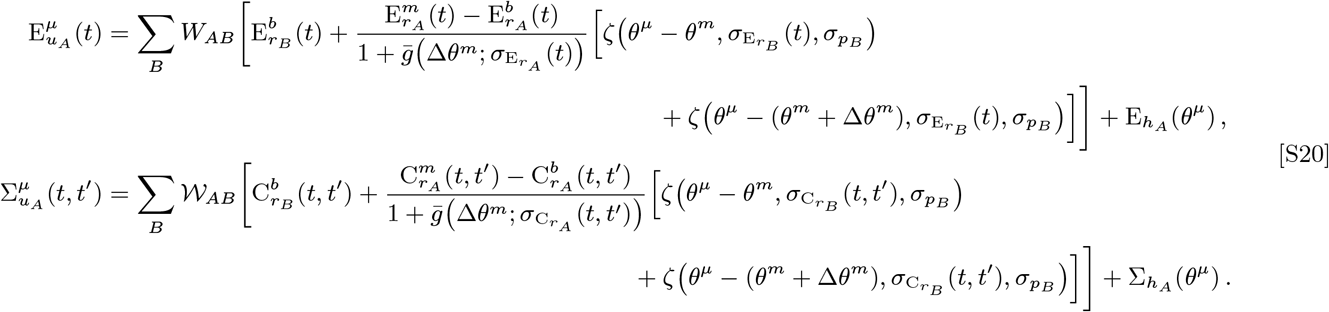

Wide overlapping Gaussians may never fully decay to the true baseline value. To account for this, we define the baseline rate moments in terms of the orientation-independent components of the net input statistics, which are given by:

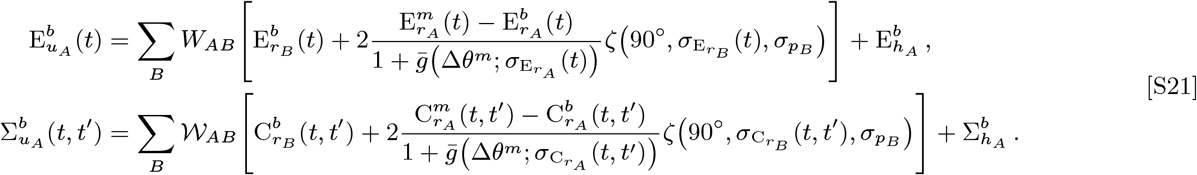

Similar expressions can be derived for the other three rate moments.

The procedure for calculating the rate moments and optogenetic response moments is similar to the single grating case, using Eq. (S19) to infer the tuning widths and Eqs. (S20) and (S21) to compute the three-site net inputs statistics. The rate moment dynamics are still given by Eq. (S13).

#### Numerical Solution to Mean-Field Equations

We can numerically solve the dynamical mean-field theory (DMFT) by discretizing Eq. (S13) in time, indexing time steps as *t*_*i*_ = *i* Δ*t* for *i* = 0, 1,…, *N*_*t*_ up to a fixed maximum time *T* = *N*_*t*_Δ*t*. Replacing the time derivatives with forward finite differences results in a naive iterative first-order Euler-like scheme for evolving the the means 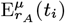 on a line of discrete times and the autocorrelations 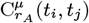 on a grid of discrete times. We can take advantage of the fact that the autocorrelation function tends to decay to a constant value for long time lags by only integrating Eq. (S13) within the band of time pairs (*t*_*i*_, *t*_*j*_) defined by a maximum time lag | *t*_*j* −_*t*_*i*_|≤ *s*_*int*_ = *N*_*i*_Δ*t*. This naive procedure is detailed as pseudocode in Alg. 1 for the rate moments and Alg. 2 for the optogenetic response autocorrelation.

The algorithm requires valid initial conditions for the rate moments at *t* = 0, *i.e*. 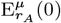 and 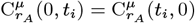 (listed in Table S3). We run Alg. 1 and extract the final mean rate 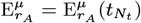 as the stationary mean rate and the final grid edge of the rate autocorrelation functions 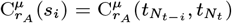 as the stationary autocorrelation as a function of the discrete time lag samples *s*_*i*_, recording up to a maximum saving time lag *s*_*sav*_ = *N*_*s* Δ_ *t*. We run this procedure twice, first with *P*(λ_*A*_) = δ (λ_*A*_) to get the rate moments without the optogenetic stimulus, and then a second time with *P*(λ_*A*_) given by Eq. (S16) to get the rate moments with optogenetic stimulation. We then use the equilibrium rate moments with and without laser stimulation to run Alg. 2 and integrate 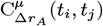, from which we extract the stationary response autocorrelation function from the final grid edge.

We list the initial conditions and the auxiliary site distance used in the dynamical mean-field analysis in Table S3. We discretize time with step size Δ*t* = 0.002 s, defining a warmup time *T*_*wrm*_ = 1.2 s in which we evolve the rate moments until they reach equilibrium and set *s*_*sav*_ = 0.4 s, which was chosen to be much longer than the decay time of the autocorrelation functions. We set the total integration time to *T* = *T*_*wrm*_ + *s*_*sav*_. and the maximum time-lag defining the band of time-points that we evolve the autocorrelation functions on is set to *s*_*int*_ = 1.5*s*_*sav*_.

We modify the procedure for evolving the rate moments to compute the stationary moments more efficiently in two ways. First, we replace the first-order Euler integration scheme used to evolve the mean rates 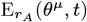 with a fourth-order Runge-Kutta method, which accelerates the convergence of the rate moments to their stationary values. Second, when 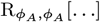 is used to compute rate autocorrelations, such as

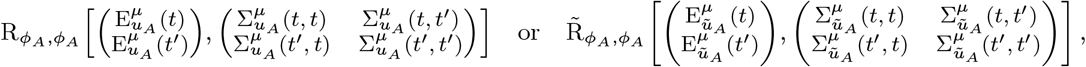

we compute them using only the mean and variance of the net inputs at the latest evolved time. That is, we make the following substitutions in Algs. 3 and 1:

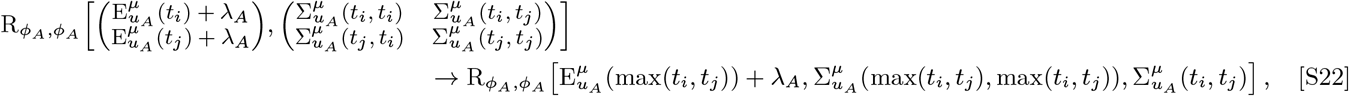

where

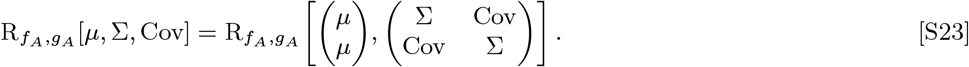

Though this assumption increases the required integration time to converge to stationary values, it reduces the number of dependent variables of the cross-correlation auxiliary functions 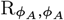 and 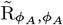 from five to three. This allows us to precompute them on a 3D grid of means, variances, and correlation coefficients and use linear interpolation to quickly calculate these quantities at the required input statistics without needing to perform costly Gaussian integrals. Similarly, we also precompute and interpolate the mean auxiliary functions 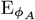 and 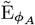 on a 2D grid of means and variances.

We cannot use the same trick when computing the cross-correlation between the rates without and with optogenetic input, defined as

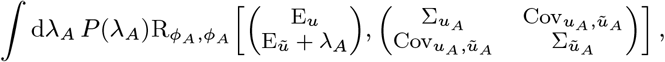

since the statistics of the net inputs without and with optogenetic stimulation are never equal. Thus, this quantity depends on five parameters (the mean and variances of the net inputs with and without optogenetic input and their covariance), meaning that precomputing and interpolating this function is infeasible. Instead, we must calculate this quantity at every step of the DMFT procedure through a Gaussian integral given by

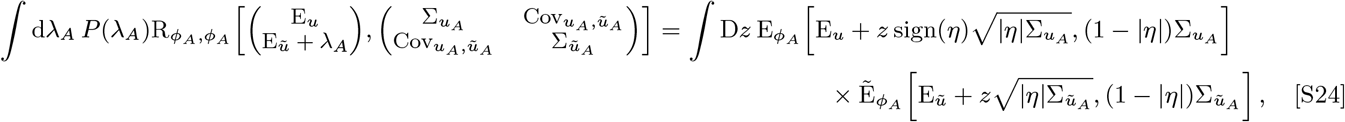

with 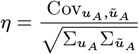

#### Computation of Time-Averaged Response Statistics

Having numerically solved for the mean-field rate moments we now must calculate the means and variances of the time-averaged rates to compare our theory results with simulations. The mean rates computed from the mean-field theory are already equal to the mean time-averaged activities. To calculate the variances of the time-averaged firing rates from the computed autocorrelation functions 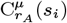 as a function of the discrete time lags *s*_*i*_, we reconstruct a 2D grid of autocorrelations at discrete pairs of time points 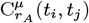 as a Toeplitz matrix where the elements along the *i*-th diagonal equal 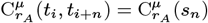 up to the time-averaging window *T*_*avg*_ = 1 s. The mean over this discrete grid of autocorrelations is equal to the mean squared time-averaged rates for that population, from which we can compute the variance of the time-averaged rates.

Lastly, we use a Monte-Carlo approach to compute the balance indices whereby we generate 20000 samples of time-averaged excitatory and inhibitory cell inputs per population from a normal distribution whose mean and variance matches those predicted by the afferent input statistics and E/I recurrent input statistics. For the optogenetically stimulated network, we also generate 20000 samples of uncorrelated optogenetic inputs, which we add to the excitatory input of the E populations.

### C. Extended Effective Two Site Model and Linear Response Analysis Methods

#### Effective Two Site Model

Labeling sites in the effective reduced network by *μ*, ν = {*b, m*}, we denote the effective mean total weights and the variance of the total weights from site ν to site *μ* as 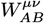 and 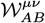, respectively. Since *W*_*AB*_ and 𝒲_*AB*_ (defined in Eq. (S12)) are the mean and variance of the total weights, *i.e*. summed across all presynaptic cells, it follows that 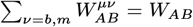 and 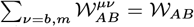 for all *μ*. Thus, we define 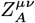 and 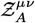 as the fraction of weight mean or variance that site *μ* receives from site ν, respectively, which allows us to express the effective weight moments of the reduced model as

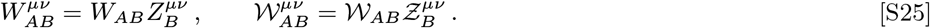

We derive the effective two-site model connectivity using the relationship between the equilibrium rate moments and net input statistics by requiring that 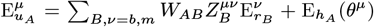 and 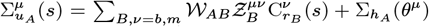.

Note that defining the two-site model connectivity in this way is equivalent to the definition of the reduced connectivity in Eq. (4) of the Main Text. Using Eq. (S11), it is straightforward to show that, for the single-grating driven network, 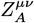 is given by

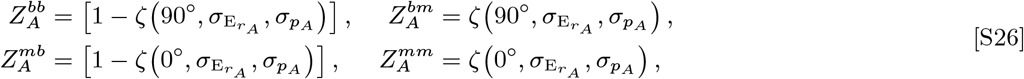

where 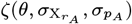 was defined in Eq. (S10). Similar equations can be derived for the fraction of weight variance 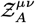 by substituting 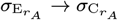 (*s*) in Eq. (S26). Because of the dependence of the rate autocorrelation function tuning widths on the time lag *s*, 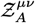 will become lag-dependent as well. However, we find that the rate autocorrelation function tuning widths have only weak time-lag-dependence except at the strongest synaptic efficacies we consider (Supp. Fig. S1a), so for simplicity we ignore the time-lag-dependence of 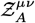.

To give a sense of scale for the effective weight moments of the effective two-site network compared to a fully unstructured network, for 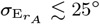 the values of *Z* are accurately approximated by

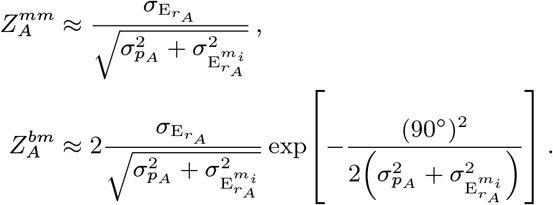

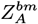 is generally small, meaning that the matched-to-baseline connectivity is typically much weaker than the baseline-to-matched connectivity. Since 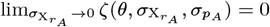, in the strongly structured (narrow tuning width) limit the mean weight fractions approach the values

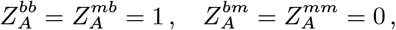

meaning that all connections are received from the baseline. Since 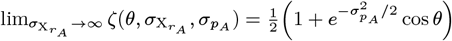, in the opposite limit of weak structure (wide tuning widths) the mean weight fractions become

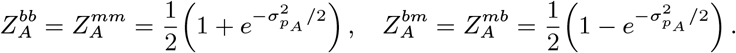

If the feedforward inputs are also weakly structured, *i.e*. the baseline and matched sites receive nearly equal afferent input, then the two-site network is invariant under an interchange of the baseline and matched site and the two-site model can be further reduced to an unstructured one-site network.

For the network driven by two equal-contrast gratings, the two-site mean weight fractions are as follows:

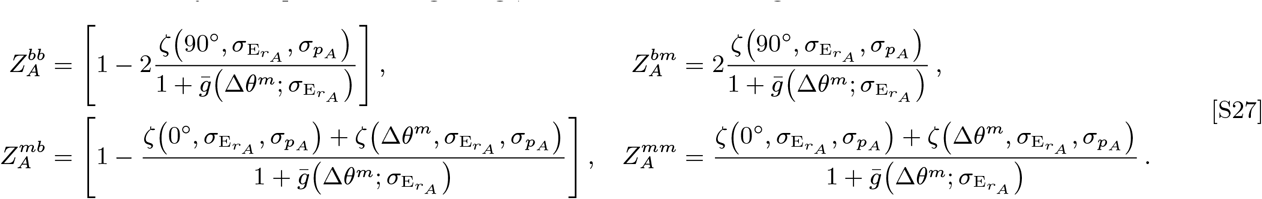

In the more general case with an arbitrary number of visual stimuli of different contrasts, we derive the reduced effective model between the untuned baseline and tuned visual-stimulus-matched sites in App. E

#### Partitioned Ring Heuristic

In the partitioning heuristic, we assign cells to physical partitions or subsets of the ring based on whether they respond more like the visual-stimulus-matched or baseline activity. Populations whose mean rate is greater than the average of the baseline and the matched mean activity of the corresponding cell type are classified as “firing like the visual-stimulus-matched activity” while the remaining locations are classified as “firing like the baseline”. That is, we define the sets 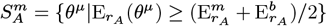 and 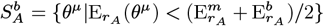 such that the size of a partition is defined as 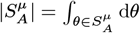. With these definitions, the fraction of mean weights in the effective two-site model is computed as

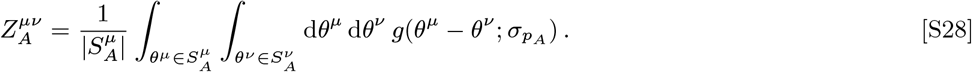

The weight variance fractions 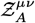 can be defined analogously by defining new partitions based on the baseline and matched rate autocorrelations. The exact mean weights and the mean weights resulting from the partitioning heuristic are plotted for comparison in Supp. Fig. S4 and S6.

#### Linear Response Analysis

In App. F we derive the general linear response of the rate moments of any random rate-based network due to a normally distributed perturbing input. To begin, we define the quantities 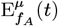 and 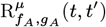 as shorthand for the mean and cross-correlation auxiliary functions (defined in Eq. (S14)) evaluated with the corresponding dynamical net input statistics of site *μ*. That is, these quantities are defined as

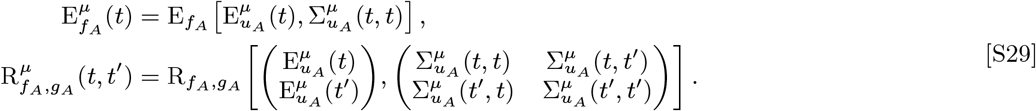

Note that 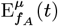 and 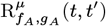 implicitly depend on the dynamical rate moments due to the dependence of the net input statistics onthe rate moments.

Next, we change coordinates such that the pairs of dynamical times for the rate autocorrelations are parameterized as 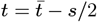 and 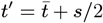. That is, 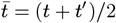 is the evolutionary time and *s* = *t*^′^ −*t* is the time lag. Finally, under this change of variables the first two lines of Eq. (S13) – which describe the dynamics of the rate means and autocorrelations – can be rewritten as follows:

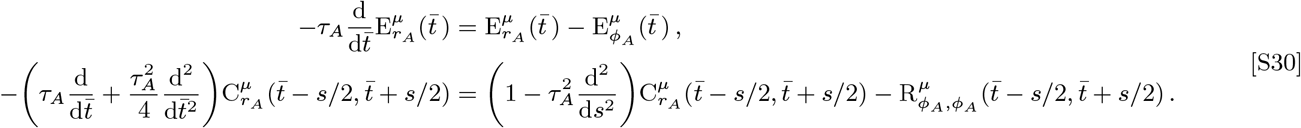

In Eq. (S30), we have separated out the derivatives with respect to the evolutionary time 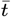 on the left side from all other terms on the right side. If we introduce a perturbation adiabatically, then the term 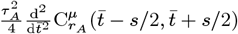 on the left side of Eq. (S30) can be approximately ignored. To compute the perturbation-induced linear response of the equilibrium rate moments, we will need to compute the Jacobian of the right side of Eq. (S30) when the system is at equilibrium. We call this quantity the rate moment Jacobian ***𝒥*** ^*μ ν*^ (*s, s* ^′^)..

We present the derivation of the Jacobian in detail in App. F. Briefly: We separate the rate moment Jacobian into ν → *μ* subcomponents ***𝒥*** ^*μ ν*^ (*s, s* ^′^). We then further separate ***𝒥*** ^*μ ν*^ (*s, s* ^′^). into four sub-blocks:

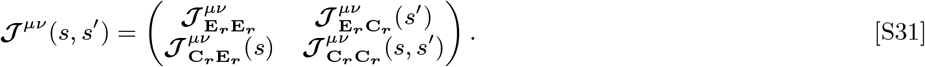

By differentiating the right side of Eq. (S30), we can derive that form of the 2 ×2 blocks defined in Eq. (S31). First, we define the equilibrium counterparts of Eq. (S29) as

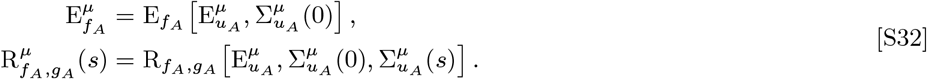

The rate moment Jacobian sub-blocks are then given by

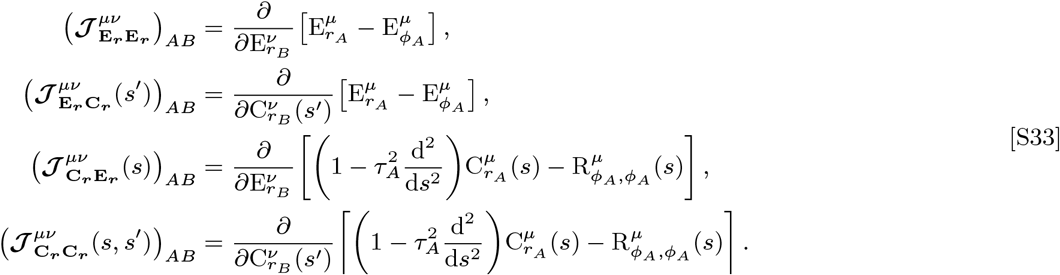

In App. H we show that inverting the Jacobian by treating the cross-site projections as perturbative results in an expansion series in powers of the effective cross-site interaction matrices ***χ***^*μθ*^ (*s, s* ^′^). These cross-site interaction matrices are given by

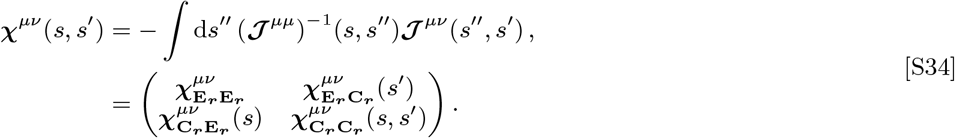

We denote the vectors of the linear response of the rate means and the rate autocorrelation functions across cell types per site as 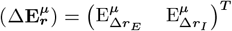 and 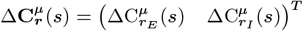, respectively. We define the decoupled response of the rate moments as the change in the moments to first order in the perturbation in the absence of dynamic recurrent interactions across sites. That is, recurrent inputs received by site *μ* from site ν ≠ *μ* are frozen at their pre-perturbation value, but the recurrence within site *μ* is kept dynamic. We denote the vectors of decoupled response of the rate mean and autocorrelation function as 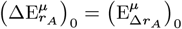 and 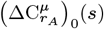, respectively. In App. H we show that the linear response can be expressed as an expansion series in terms of these cross-site interaction matrices acting on the decoupled responses. The expansion series relating the decoupled response to the linear response is given by

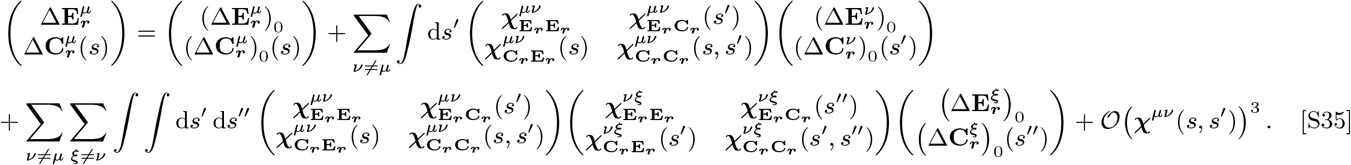

Even when the perturbing inputs are not normally distributed, the changes in recurrent input between sites are guaranteed to be normally distributed because each cell receives a large number of independent presynaptic inputs (see App. B for a more in-depth discussion of this). By treating the changes in the cross-site recurrent input as the normally distributed input, Eq. (S35) is valid even for our log-normally distributed optogenetic perturbation as long as the expansion series converges.

In App. I we also show that 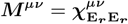 takes the form

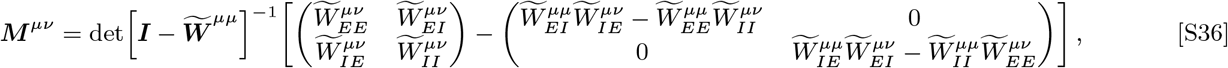

where 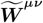 is the 2**×**2 effective coupling matrix, defined in the appendix at Eq. (S85). In the absence of disorder, the effective coupling matrix is given by the product of the mean gains per population 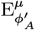 and the mean connection matrix 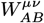. However, the disorder will augment the effective coupling in potentially unpredictable ways. Regardless, Eq. (S36) has a simple form in the strongly coupled and feedback inhibition dominated regime, as explained in the Main Text. The rate moments must be stable to perturbations in order for their dynamics to approach a stable equilibrium. However, the linear response analysis in the mean-interaction approximation – which ignores perturbation-induced rate autocorrelation changes – only yields valid results if the determinant that appears in Eq. (S36) is positive. Indeed, we find that for the best-fit network 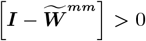 robustly even as we vary the strength of the network coupling and structure (Supp. Fig. S1b). Furthermore, although the elements of 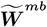 could break Dale’s law, we find that the signs of the elements obey Dale’s law for all but one set of varied coupling and structure strengths (Supp. Fig. S1c).

#### Numerical Calculation of Linear Responses

To numerically solve for the linear response from the decoupled optogenetic response, we start with the 1D array of the rate autocorrelation functions sampled at discretized time-lag steps 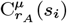 computed from the mean-field theory at the baseline, matched, and auxiliary sites. Using either Eq. (S9) or Eq. (S19), we then inferred the stationary mean rate tuning width 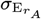 and the approximately time-lag-independent rate autocorrelation tuning width 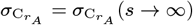. Using Eq. (S26) or Eq. (S27) (or the equivalent expression for 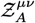) we compute the fraction of mean weights 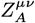 and the fraction of weight variance 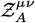. Eq. (S25) gives us the reduced model weight moments, from which we calculate the net input means 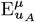 and the vector representation of the net input autocovariance functions at the same discretized time-lags 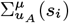 at the reduced sites *μ* = {*b, m*}. Finally, we calculate the various means and cross-correlations of the activation functions and their derivatives required for the linear response analysis by numerically differentiating the precomputed and interpolated functions using the following equations

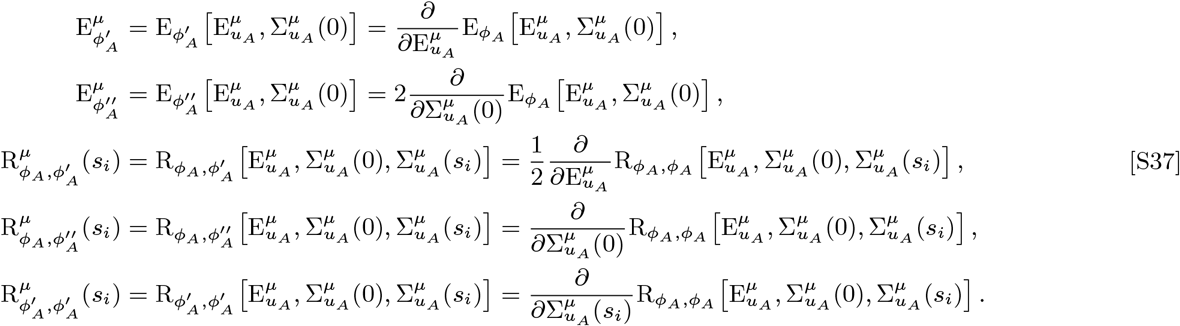

The cross-correlations are also saved as 1D arrays whose elements correspond to the discrete time-lags *s*_*i*_. In practice, 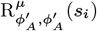 often has a “step-like” dependence on the time lag due to the course grid of input covariances used to precompute and interpolate 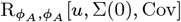. This can be fixed by applying a low-pass Fermi filter on 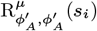, but our results do not significantly change with or without the low-pass smoothing.

We construct the subblocks of the time-lag-discretized linear response matrices ***𝒥***^*μθ*^ (*s*_*i*_, *s*_*j*_) (derived in App. F and written with continuous time lags in Eq. (S78)) as

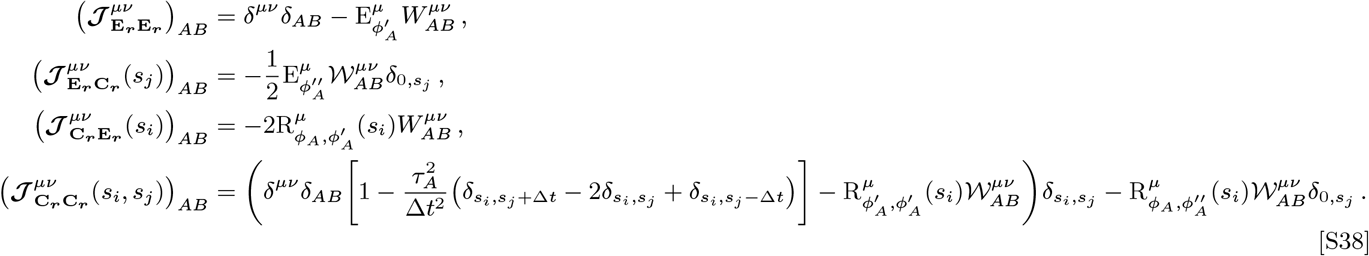

The time-lag-discretized effective cross-site interaction matrices ***χ***^*μθ*^ (*s*_*i*_, *s*_*j*_). can then be computed from Eq. (S34) by numerically inverting ***𝒥***^*μμ*^(*s*_*i*_, *s*_*j*_).

To calculate the linear responses in Fig. 5, we first reduce the visually-evoked network at a given contrast to its corresponding two-site model. From the effective coupling of the reduced model and the pre-perturbed rate moments, we then calculate the distribution of recurrent inputs that a site receives from other sites and freeze them to the pre-perturbation values. We decouple all sites by treating them as separate unstructured random networks with mean total weights 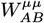 and total weight variances 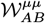. We then compute the decoupled response to the optogenetic stimulus by rerunning the DMFT procedure for each decoupled site in the presence of the optogenetic stimulus. From the non-perturbatively computed decoupled response, we use Eq. (S35) (neglecting matched-to-baseline projections) to get the linear response including rate autocorrelation changes and use Eq. 5 from the Main Text to get the linear mean response with the mean-interaction approximation. Since the tuning widths are nearly invariant across stimulus conditions, we assume that the feature-dependence of the perturbed rate moments (with either the non-perturbative decoupled response or the linear response) has the same tuning width as the visually-evoked network but with different baseline and visual-stimulus-matched values.

### D. Derivation of the Self-Consistent Dynamical Mean Field Equations of the Random Ring Network

In this appendix, we derive the self-consistent dynamical mean-field equations for a random ring driven by an arbitrary normally distributed visual input with and without optogenetic stimulation. We largely follow the derivation in (7), except we consider a network with spatial structure. We then derive the self-consistent equations for the changes in firing rates induced by the laser.

We define the mean and autocorrelation function of a population’s visually-evoked firing rates as

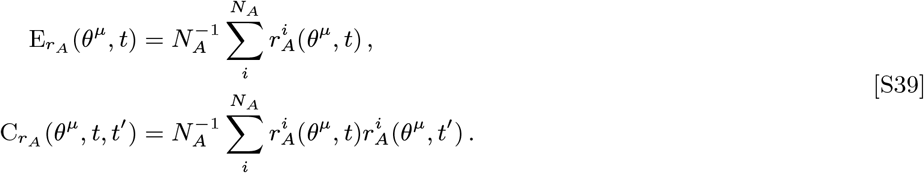

We can calculate the autocovariance function of the firing rates from the autocorrelation function using the expression 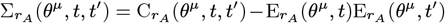. Similarly, we define the mean and autocovariance function of a population’s net visually-evoked inputs as

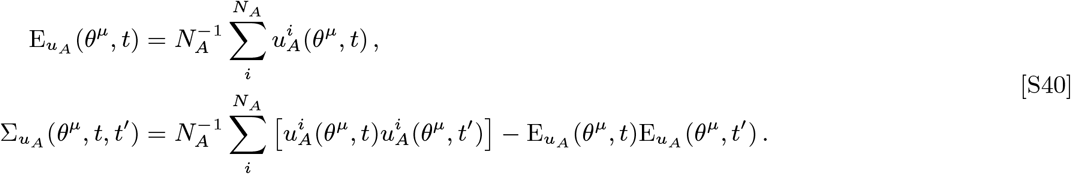

Likewise, we can define the mean and autocorrelation function of the optogenetically-evoked rates, 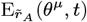 and 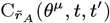, and the associated net input statistics accordingly.

If the network is highly connected such that cells receive a large number of connections from each population, the central limit theorem guarantees that a population’s recurrent inputs can be approximately described by a Gaussian process (8, 9). Furthermore, net inputs to different cells will be uncorrelated if elements in the connectivity matrix are uncorrelated. We let 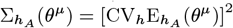 denote the variance of the afferent inputs and 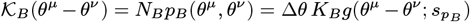 denote the density of the expected in-degree from site *θ*^*ν*^ to site *θ*^*μ*^. Substituting Eq. (S3) into Eq. (S40) and neglecting contributions of 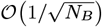 allows us to write the dependence of the mean net non-optogenetic input mean on the rate means as

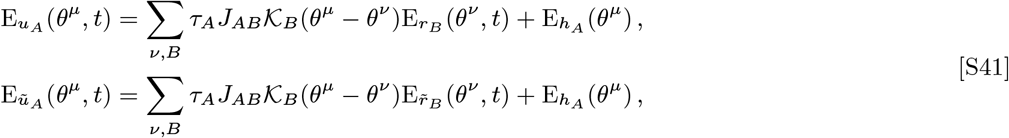

and the dependence of the net input autocovariance functions on the rate moments as

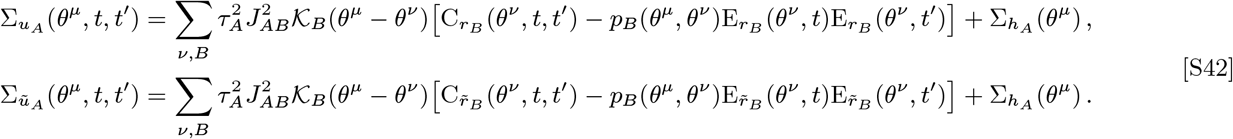

We choose the model to have sparse connectivity, where *K*_*A*_ is fixed as we vary the network size, meaning the connection probabilities *p*_*B*_ (*θ*^*μ*^, *θ*^*ν*^) scale as 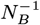. The alternative, having dense connectivity, would mean that *p*_*B*_ (*θ*^*μ*^, *θ*^*ν*^) remains fixed while *K*_*B*_ both scale proportionally with *N*_*B*_. Dense connectivity would require us to scale the synaptic efficacies as 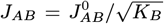 to ensure that the variance of the net inputs neither blows up nor shrinks to zero as the network size becomes large. Dense connectivity, in turn, forces the network to become unrealistically strongly coupled in the large network limit (10, 11). A sparsely connected network avoids the need to scale the efficacies, allowing the network to operate in a loosely balanced regime as *N*_*B*_ → ∞ In addition, sparse connectivity causes the term 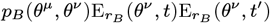 in Eq. (S42) to become negligible as we increase the network size. With the best-fit model parameters, 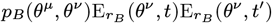 is at most only 6% as large as 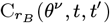. Ignoring this term does not noticeably change the rate moments calculated from the mean-field theory, so we will not consider this term to simplify our analysis under the ansatz that the feature-dependence of the rate moments is baseline-plus-Gaussian shaped.

To simplify the expressions of the self-consistent equations, we introduce two auxiliary functions. The first is the mean auxiliary function 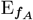 :

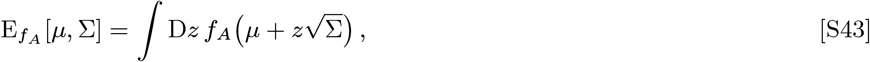

where 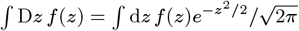 indicates a standard Gaussian integral over *f*(*z*).

To compute cross-correlations over joint-Gaussian random variables we introduce three uncorrelated standard-normal random variables *y*_1_, *y*_2_, and *z*. Letting 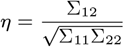 denote the correlation coefficient between *x*_1_ and *x*_2_, we generate correlated random inputs distributed as 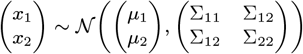 from weighted linear sums of *y*_1_, *y*_2_, and *z* as follows:

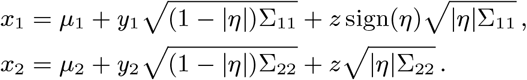

It is straightforward to show that *x*_1_ and *x*_2_ have the desired means and covariance matrix. Using this parameterization for correlated inputs, we define the cross-correlation auxiliary function 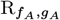 as

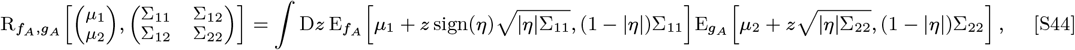

It will often be useful to define an alternative cross-correlation auxiliary function with equal input means and variances

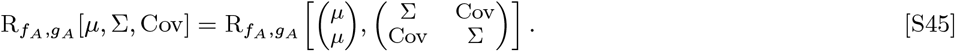

In the presence of log-normally distributed optogenetic inputs, we must integrate them separately from the normally distributed recurrent and afferent inputs. We introduce modified auxiliary functions to account for this:

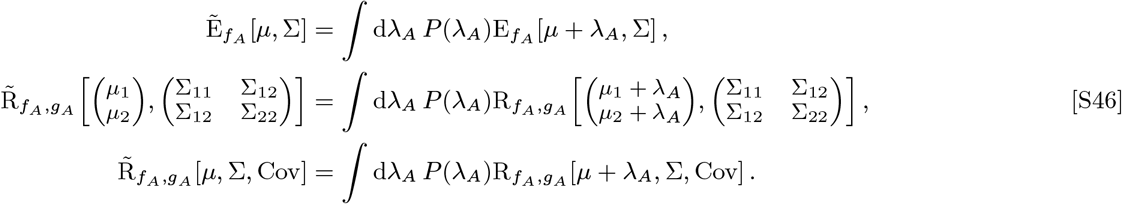

The density of optogenetic inputs is defined as:

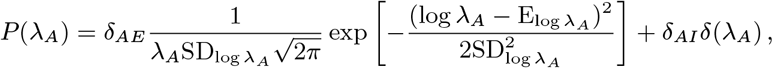

where 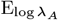 and 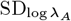 are given by Eq. (S15).

We take the population average of Eq. (S1) to derive the equation governing the dynamics of the rate mean in response to only visual inputs, *r*_*A*_, and in response to visual and optogenetic inputs, 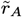, which are given by

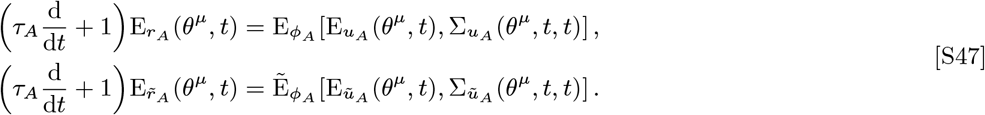

By taking the population average of the product of two copies of Eq. (S1), one at time *t* and another at time *t*^′^, we derive the equation governing the dynamics of the rate autocorrelation function, which are given by

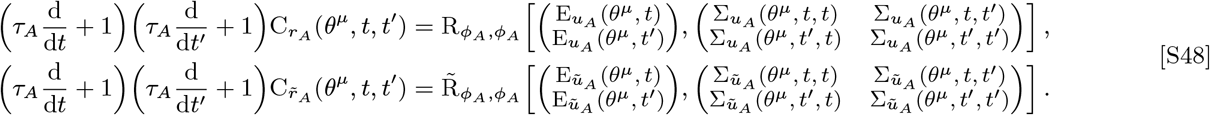

Eqs. (S41), (S42), (S47) and (S48) are the full set of self-consistent dynamical mean-field equations for the firing rate moments on the random ring.

We can numerically solve the dynamical mean-field theory (DMFT) by evolving the rate moments Eqs. (S47) and (S48) from any arbitrary stable and valid (non-negative) initial conditions until the mean rate reaches a stationary value 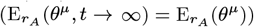 and the autocorrelation function of the rate reaches a stationary function of the difference in the time points 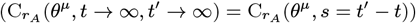. The numerical procedure is similar to the process outlined in Alg. 1, except that we must track the discrete mean rates and rate autocorrelations at all discrete sites *θ*^*μ*^. The pseudocode for the procedure on the ring, with no assumptions about the feature-dependence of the rate moments, is outlined in Alg. 3.

#### Algorithm 3

DMFT algorithm for calculating rate mean and autocorrelation function with rate-based dynamics

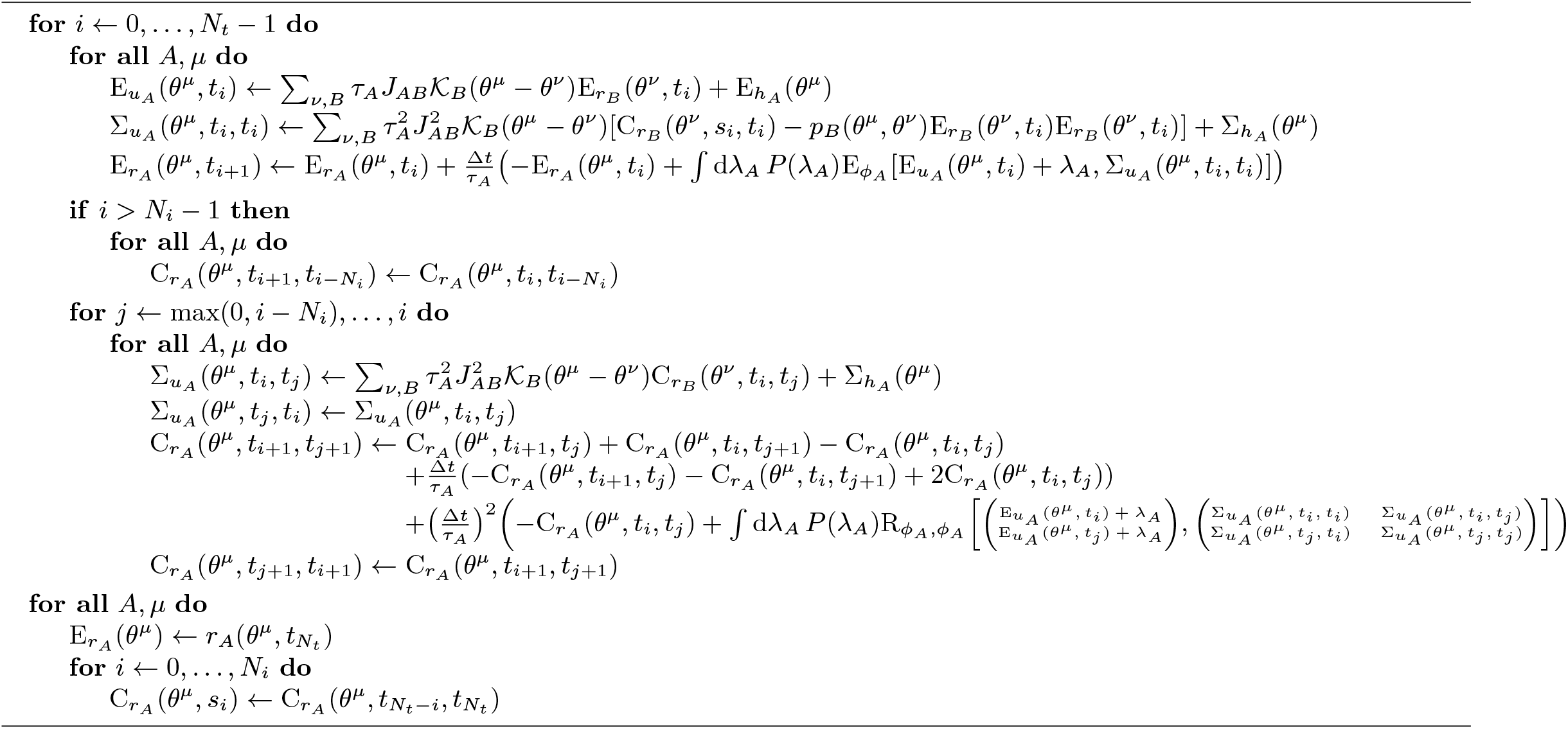

Next, we define the optogenetic responses, *i.e*. the optogenetically induced changes in the rates, as 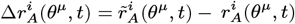. We define its mean 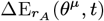 and autocorrelation function 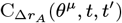 in terms of population averages as in Eq. (S39). Note that (7) chooses to self-consistently compute the autocorrelation function between 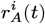 and 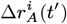. However, this quantity does not generally have a baseline-plus-Gaussian feature-dependence, and so we choose to self-consistently compute the autocorrelation function of the rate changes, which is generally baseline-plus-Gaussian shaped.

We also define the change in the net non-optogenetic inputs 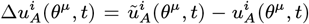, whose autocovariance function’s dependence on the rate change autocorrelations is given by

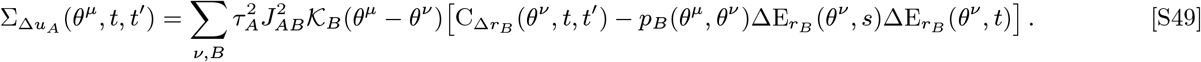

By subtracting the two lines of Eq. (S1), we derive the equation governing the dynamics of the optogenetic responses:

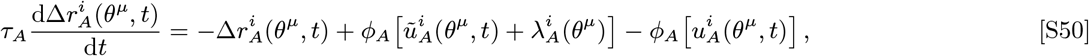

By taking the population average of the product of two copies of Eq. (S50) at times *t* and *t*^′^ and defining the cross-covariance function between the net non-optogenetic inputs without and with the optogenetic stimulus as

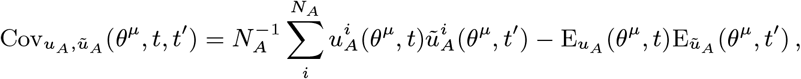

we derive the equation governing the dynamics of the autocorrelation function of the changes in rates as given by:

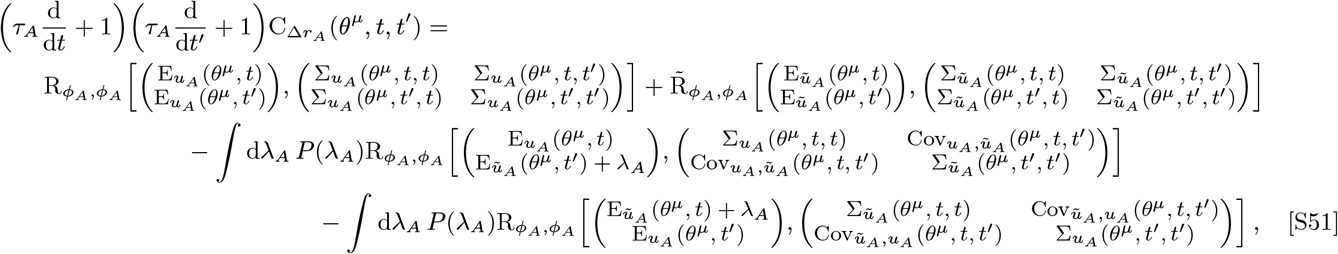

Eqs. (S49) and (S51) are the self-consistent equations for the optogenetic response moments.

To calculate the cross-covariance function between the initial and optogenetically perturbed inputs 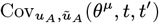, we first note that after a sufficiently long time evolution of the rate dynamics, the chaotic fluctuations of the firing rates and the net inputs before and after optogenetic stimulation will be independent of one another. Thus, 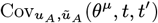 becomes symmetric under the exchange of *t* and *t*^′^ for large times. Since this is true for large times, we can approximate it as true for all times without changing the long-time-evolved stationary state. We use the fact that 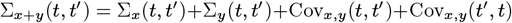 to write

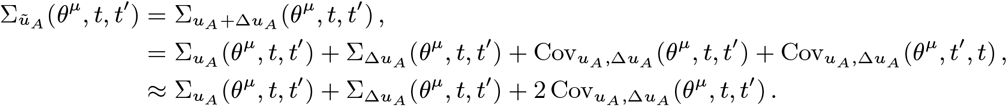

Due to the linearity of the cross-covariance function, we can write

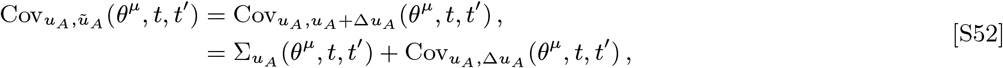

From the above two equations, we can derive that the cross-covariance function between the inputs under the approximation of time-exchange symmetry is as follows

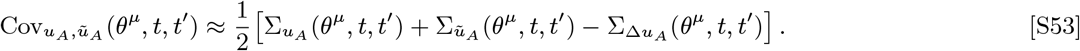

Assuming we’ve already calculated the stationary rate moments with and without optogentic input, we compute the autocorrelation function of the changes in rates by evolving Eq. (S51) from stable and valid initial conditions until it reaches a stationary function of the time lag 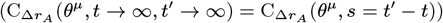. The procedure is similar to Alg. 3 and is outlined in Alg. 4.

#### Algorithm 4

DMFT algorithm for calculating the optogenetic response autocorrelation function with rate-based dynamics

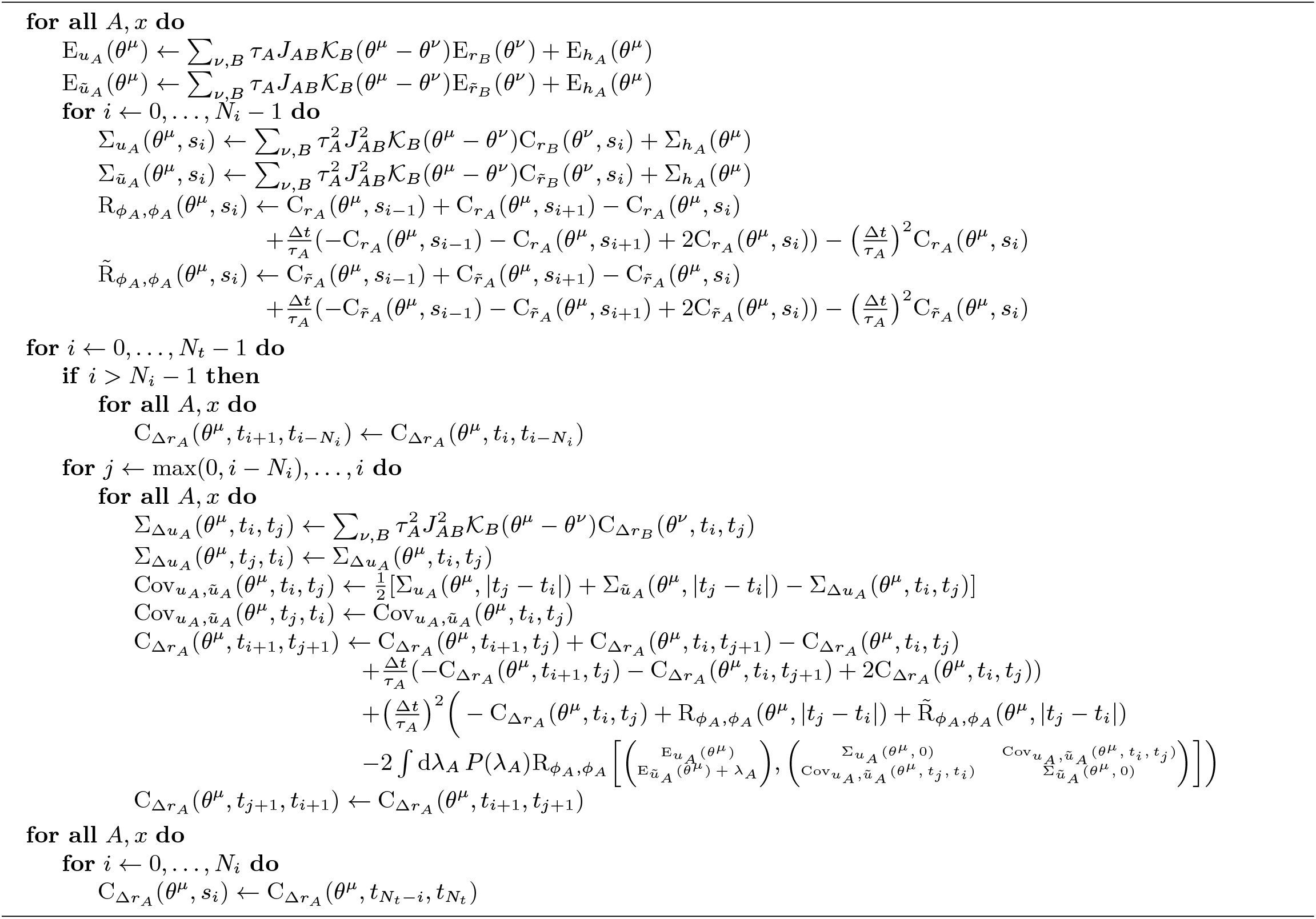

### E. Derivation of the Equivalent weight moments in a Reduced *N* +1-site Network

In this appendix, we derive the equivalent weights of an effective *N*+1-site network for a random ring driven by *N* visual stimuli. Due to potential overlap between nearby Gaussian components, deriving the equivalent weights for the reduced *N*+1-site network for *N* > 1 is more complicated than the *N* = 1 case and deserves its own appendix section. To highlight this difficulty, denote each visual-stimulus-matched location as *m*_*i*_ for *i* = 1, 2,…,*N* and consider a baseline-plus-Gaussian-mixture *a*(*θ*;^*μ*^) parameterized as

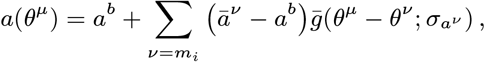

where 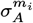 is the tuning width of the *i*-th Gaussian component. In the presence of more than one Gaussian component, unless there are only two orthogonal Gaussian components, then in general 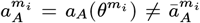 due to overlapping Gaussian components. However, by evaluating the above expression at the visual-stimulus-matched locations, we can derive a linear relationship between the true values at the visual-stimulus-matched locations 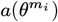 and the baseline summed amplitudes of the Gaussian components 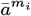. Defining ***O***_***a***_ as the *N*×*N* matrix of Gaussian overlap with elements

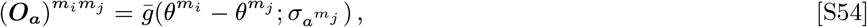

this linear relationship can be written as

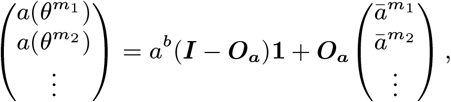

where **1** is the vector whose elements are all one. Note that if 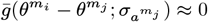 for *i* ≠ *j* then ***O***_***a***_ ***I*** and 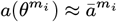. Having formally defined the Gaussian overlap matrix, we can choose to parameterize *a*(*θ*^*μ*^) in terms of the actual visualstimulus-matched values 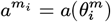 by choosing 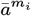 such that

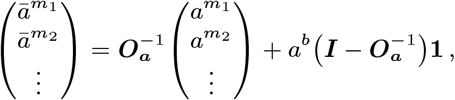

or expressed in terms of the elements of the inverse Gaussian overlap matrix:

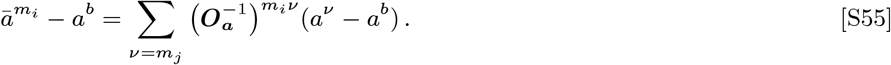

With this in mind, our equilibrium rate moment ansatzes will be parameterized as:

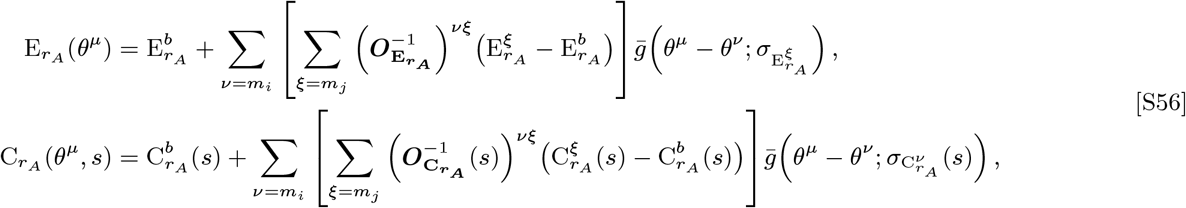

Wide, overlapping Gaussian components may never decay to the true baseline values on a ring. To address this, we will define the baseline rate moments in terms of the baselines of the net input statistics. We denoting the *B* → *A* mean recurrent input as

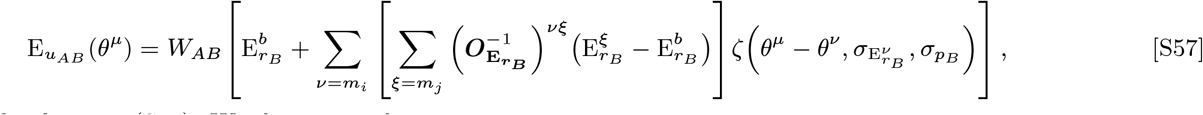

where ζ was defined in Eq. (S10). We then write the mean net inputs as

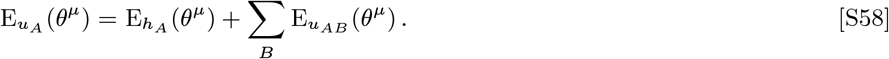

We let 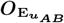 be the overlap matrix of the Gaussian components of the *B* → *A* recurrent input, where the *i*-th Gaussian component has width 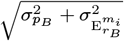. We then define the baseline of the mean *B* → *A* recurrent input as the value 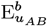 that is uniquely determined by the following parameterization of 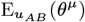 :

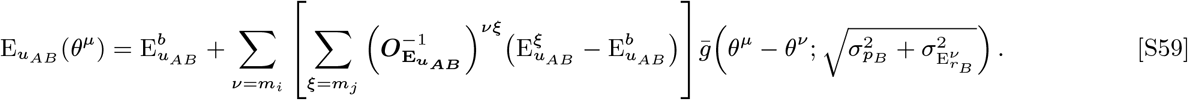

This yields the following expression for the baseline of the mean recurrent inputs:

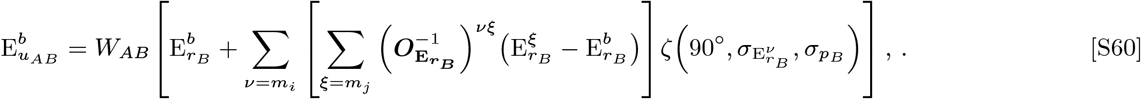

We note that if the Gaussian components in the rate moments are sufficiently narrow and spread out such that a physical location *θ*^*b*^ on the ring can be identified that satisfies the property 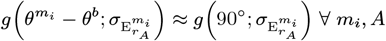, then evaluating Eq. (S57) at this *θ* ^*b*^ will result in approximately the same expression as Eq. (S60).

Now we evaluate Eq. (S57) at the visual-stimulus-matched site locations, which gives us

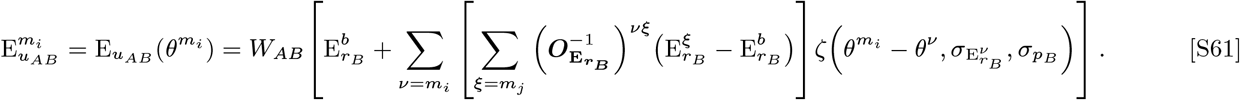

Using the relationship 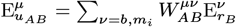 to define the mean total weights of the *N*+1-site network, from Eqs. (S55), (S60) and (S61) we derive that the mean weight fractions are given by

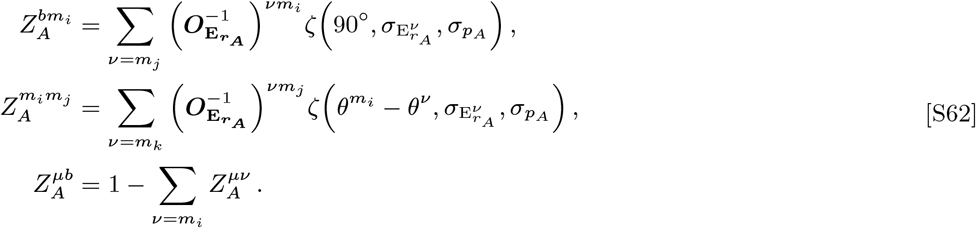

Likewise, the fractions of weight variance between sites 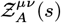 is given by Eq. (S62) under the following substitutions: 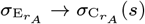 and 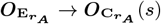. Similar to the results for the two-site network, 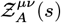 explicitly depends on the time lag *s*, but its dependence is generally mild and well approximated as time-lag-independent.

When the network is driven by two equal-contrast gratings, then the overlap matrices are symmetric 2 × 2 matrices. Summing over the rows of their inverses results in the expression for the mean rate given by Eq. (S18) and the expressions for the two-site mean weight fractions given by Eq. (S27).

### F. Derivation of Linear Response to Normally-Distributed Input Perturbations

In this appendix, we derive the equations for the linear response of the network rate moments to a normally distributed perturbing input. The self-consistent equations that the equilibrium rate moments must satisfy are listed in Eq. (S17). For notational brevity, we denote the mean and cross-correlation auxiliary functions evaluated with the equilibrium net input statistics at site *μ* as

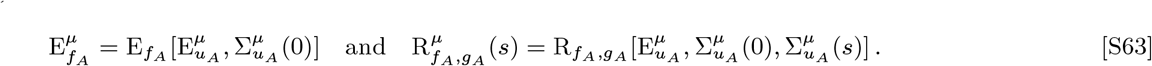

We note that while we have dropped the arguments of 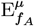 and 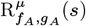, these quantities are functions of the equilibrium rate moments due to the dependence of the net input statistics on the rate moments. In summary, in the absence of an optogenetic stimulus, the equilibrium rate moments satisfy following set of equations:

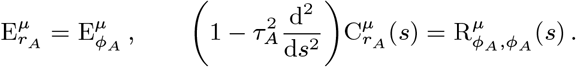

We now introduce a normally distributed perturbation that changes the net input statistics to 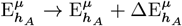 and 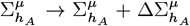. The goal now is to compute the linear response of the rate moments induced by the perturbation: 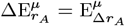 and 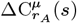 To begin, we compute the change in 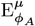 induced by the perturbation as follows:

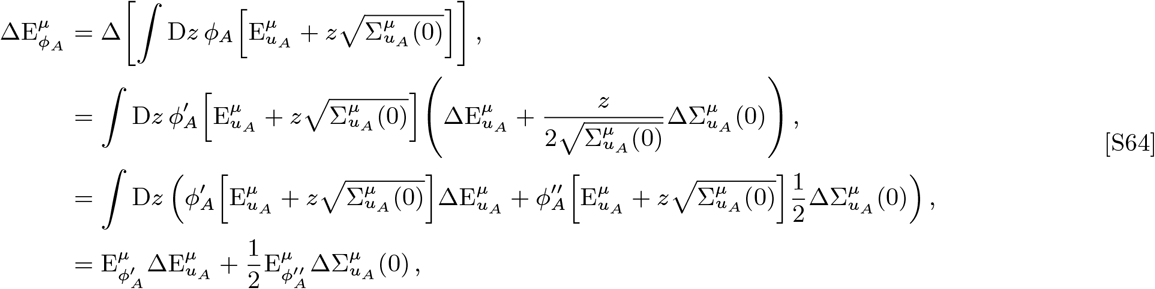

In the third line we use Bussgang’s theorem (also sometimes known as Stein’s lemma), which states that 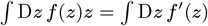.

Next, we calculate the perturbation-induced change in 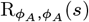 as follows:

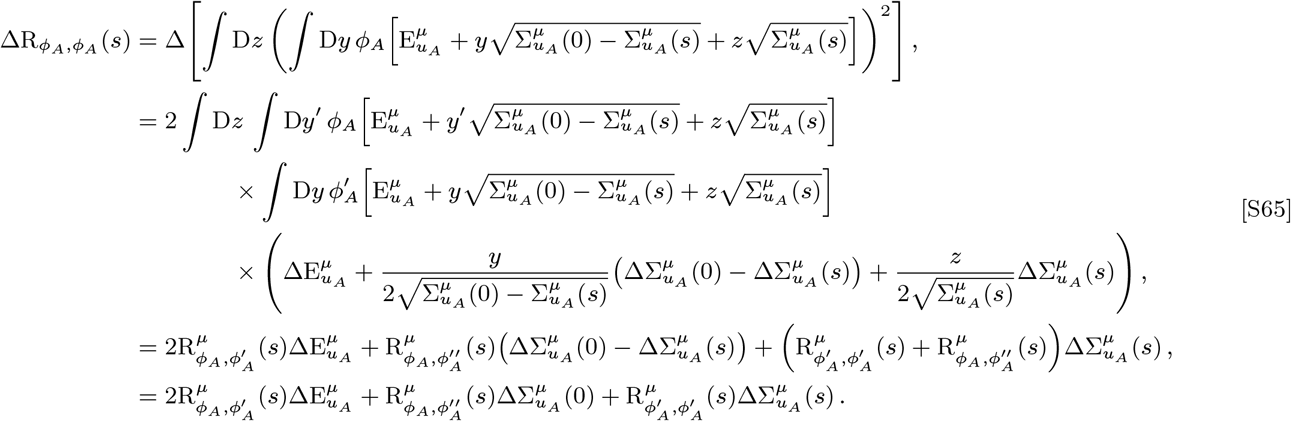

If the weights are unchanged by the perturbation, then the perturbation-induced changes in the net input statistics are given by

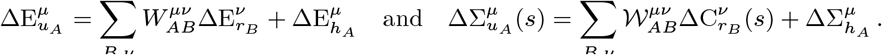

By substituting the above experssions into Eqs. (S64) and (S65), we derive a coupled system of equations that the linear response of the rate moments must satisfy:

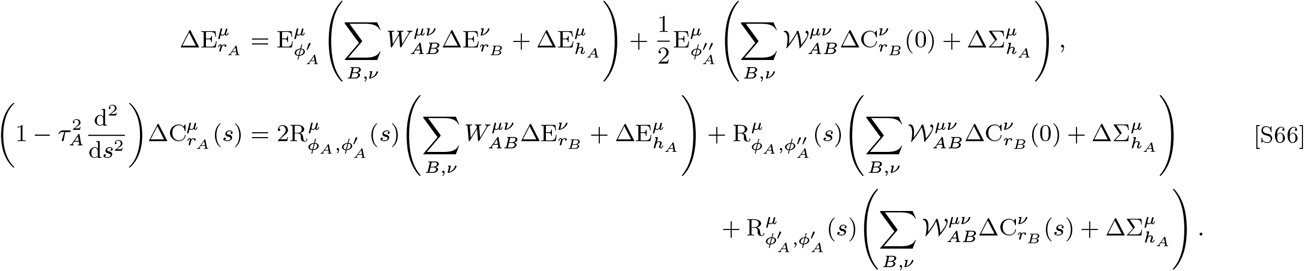

We define the linear response driving terms as the terms in Eq. (S66) that are proportional to changes in the external inputs such as 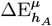 and 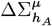. Under this definition, the linear response driving terms are given by

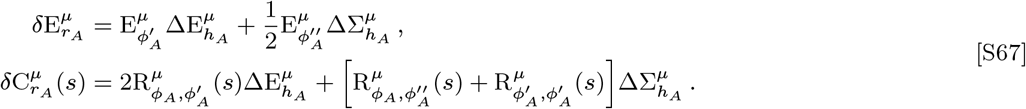

We can now rearrange Eq. (S66) to derive the relationship of the rate moment responses on the linear response driving terms. To express this relationship more compactly, we denote Δ**E**_***r***_ and Δ**C**_***r***_(*s*) as the values of 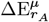 and 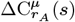 across all populations (*i.e*. all cell types and sites) flattened into one-dimensional vectors. We similarly define δ **E**_***r***_, δ **C**_***r***_(*s*), **E**_***f***_, **R**_***f***,***g***_(*s*), and ***τ***^***2***^ as population-flattened vectors whose entries are equal to the corresponding values from each population. The choice of how we arrange populations to elements of the flattened vector elements naturally defines the population-flattened matrices ***W*** and. **𝒲** Finally, we denote D(***a***) as a diagonal matrix whose entries along the diagonal are equal to the entries of the vector ***a***. Now that we have defined the population-flattened representations of the terms appearing in the linear response, then Eq. (S66) can be written in matrix notation as

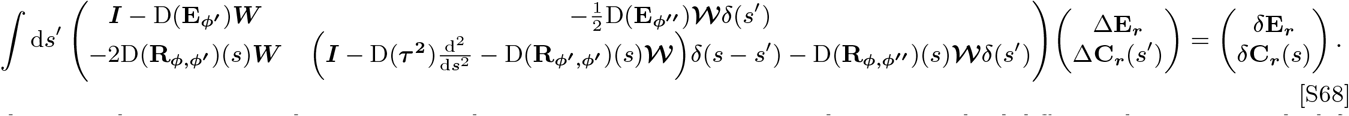

Thus, in order to compute the rate moment linear response, we must invert the matrix-valued differential operator on the left side of Eq. (S68).

Though it is likely impossible to analytically invert this matrix-valued differential operator, we can symbolically express the solution to Eq. (S68). We first define ***G***_0_(*s, s*^′^) as the matrix-valued Green’s function that satisfies

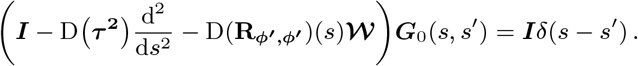

We note that ***G***_0_(*s, s*^′^) cannot be expressed analytically, so we refer to this Green’s function symbolically. Next, we define the function ***G***(*s, s*^′^) as the solution to the following equation:

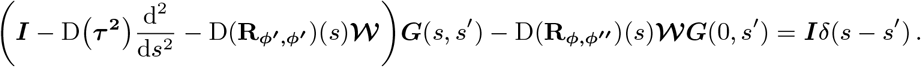

By using the generalization of the Woodbury matrix identity for linear operators, **δ*G***(*s, s*^′^) *can* be expressed in terms of ***G***_0_(*s, s*^′^):

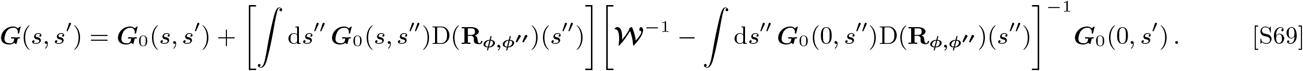

Finally, we define ***δ G***(*s, s*^′^)) such that ***G***(*s, s*^′^))+ ***δ G***(*s, s*^′^)) is the solution to the following equation:

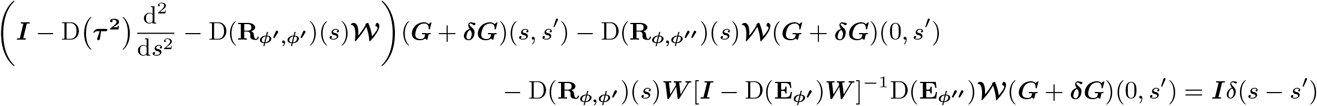

Again, using the Woodbury matrix identity, ***δ G***(*s, s*^′^) can be expressed as

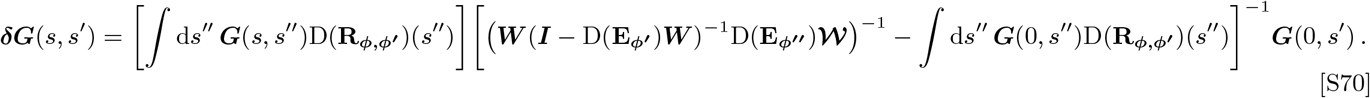

Having defined these functions, the rate moment linear response can then be expressed as

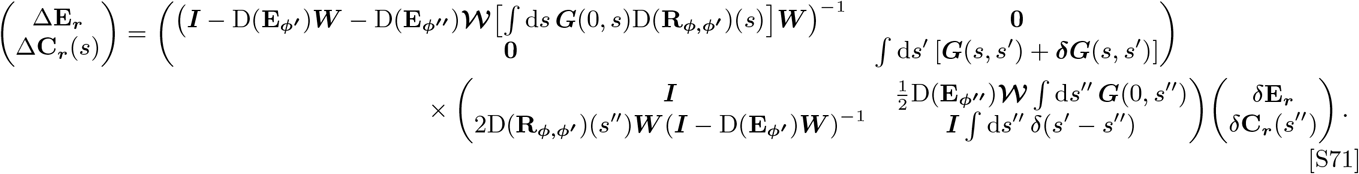

One limit in which we can exactly solve Eq. (S68) is if the cross-correlation functions appearing in the equations can be assumed to be independent of the time-lags, *i.e*. 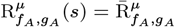 can all be expressed in Fourier space as follows:. In this case, the functions ***G***_0_(*s, s*^′^), ***G***(*s, s*^′^), and **δ*G***(*s, s*^′^) can all be expressed in Fourier space as follows:

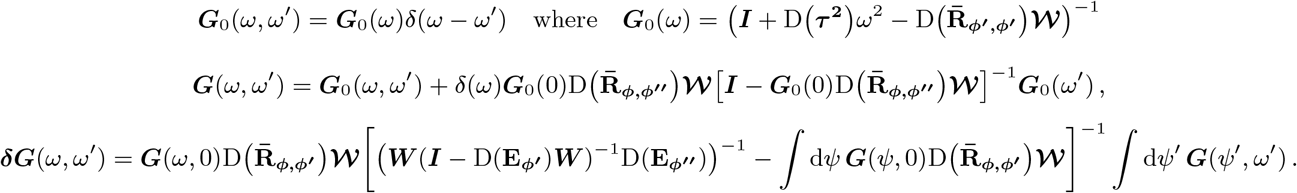

Thus, in Fourier space we can express the exact linear response under this approximation as

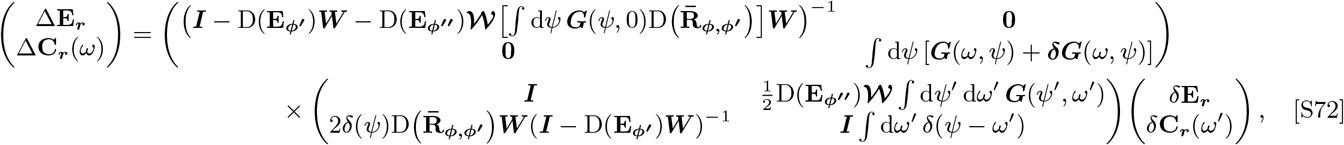

Above, we explicitly considered the case that the perturbing input is normally distributed. However, we have empirically found that we can approximately compute the linear response to the optogenetic stimulus by computing the linear response driving terms as

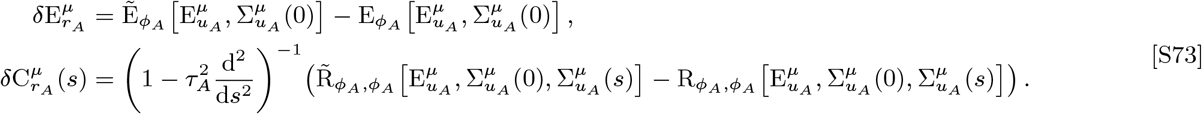

However, we choose not to compute the linear response driving terms using Eq. (S73) because it adds additional error to our linear response estimate compared to the approach we employ in the Main Text based on non-perturbatively computing the decoupled response (see App. C). since the only errors incurred by computing the linear response from the non-perturbative decoupled response are due to our assumption that the tuning widths are unchanged by the perturbation.

Finally, we shall discuss numerical approaches to solving Eq. (S68). To deal with the second derivative w.r.t. the time lag *t* in Eq. (S68), we can discretize the lags into even time points *s*_*i*_ = Δ*s* × *i* for *i* = 0, 1,…, *N*_*t −*_1 to replace the second derivative with central finite differences. The second derivative in Eq. (S68) then becomes

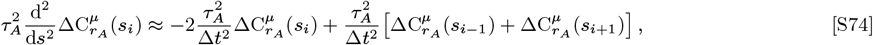

where we can impose our boundary conditions by defining 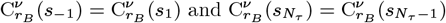 since the rate autocorrelation function is an even function of σ and decays to a constant value for sufficiently long time lags (6). By flattening δ **E**_***r***_, δ **C**_***r***_(*s*), **E**_***f***_, **R**_***f***,***g***_(*s*), in time and using Eq. (S74), we can then cast Eq. (S68) into the form of a matrix equation that can be solved numerically.

### G. Connection between Linear Response on the Random Ring to the Response on *N* +1-site Network

The linear response developed in App. F is general enough to be applied to any rate-based random network, including the random ring model. In this appendix, we highlight the approximations necessary to connect the linear response of the full ring with the response of the eFFective two-site network. Reintroducing the explicit dependence of the ring location *θ*^*μ*^, we denote the mean and cross-correlation auxiliary functions evaluated on the equilibrium net input statistics at site *θ*^*μ*^ as

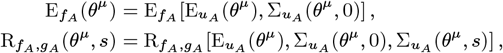

Having previously defined 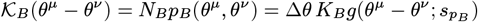 in App. D we can express Eq. (S66) as

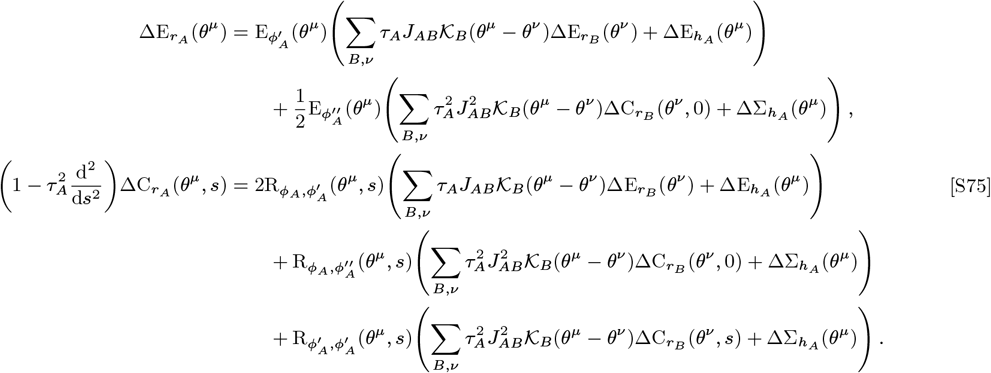

The reduction from the random ring to the effective *N*+1-site model requires that the feature-dependence of the rate moments be well-described by a baseline-plus-Gaussian-mixture whose tuning widths are unchanged by stimulus conditions, including upon adding the perturbation. Thus assumption forces the rate moment responses to be parameterizeable as:

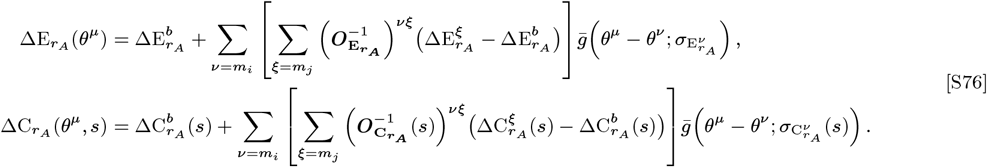

Now we compute the convolution between the mean weights and the mean rate responses as follows:

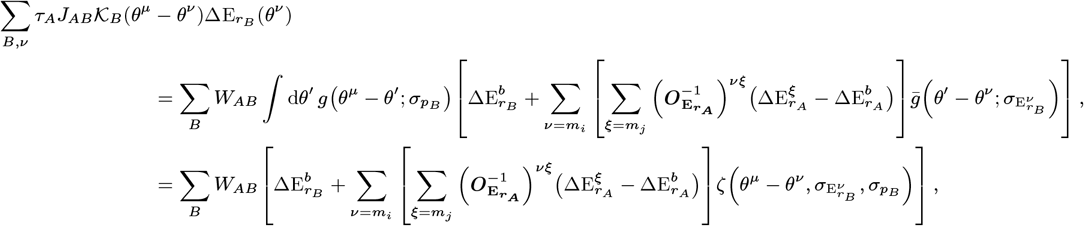

The convolution 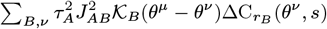 can be derived in a similar fashion.

Evaluating these convolutions at 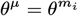 and using the definitions of 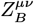 from Eq. (S62) results in the expression

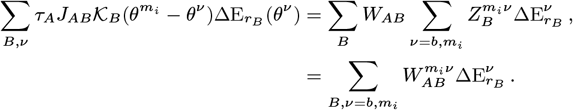

We then evaluate Eq. (S75) at 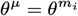, resulting in

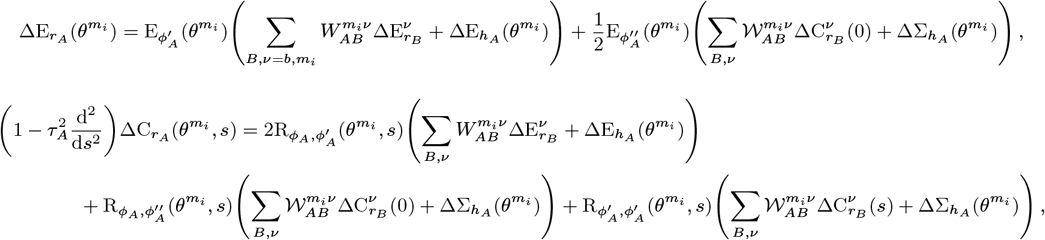

By substituting quantities on the ring at 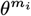 with the equivalent quantities on the effective reduced model – *i.e*. identifying 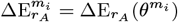 and 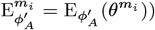 and so on – we will have matched the rate moment responses at visual-stimulusmatched sites between the ring and the *N*+1-site network.

If the Gaussian components in the rate moments are sufficiently narrow and spread out – or if the network is only driven by a visual stimulus with a single Gaussian component – then we can identify a location *θ*^*b*^ on the ring that satisfies the property *g* 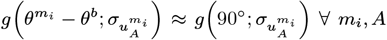. In this case alone, evaluating Eq. (S75) at *θ*^*b*^ will match the linear response susceptibilities of the baseline in the effective reduced model. If this is not the case, then the baseline does not physically exist on the ring and must be defined in terms of the baseline net input, as explained in App. E. Doing so involves similar algebra as above, which we will omit for brevity, but results in a matching between the baseline rate moment responses on the ring and the *N*+1-site network.

### H. Derivation of Linear Response as Expansion Series in Powers of Cross-Site Projections

The effective cross-site projections are often sufficiently weak in strength compared to within-site couplings that they can be treated perturbatively, allowing us to express the linear response susceptibilities as an expansion series in powers of the cross-site projections. In particular, this expansion series empirically converges for all two-site networks considered in this work. In this appendix, we derive the form of this series expansion. Let us define the rate moment Jacobian 𝒥 as the matrix-valued linear operator on the LHS of Eq. (S68) (see Methods in the Main Text or App. C for a justification of why this can be interpreted as a Jacobian):

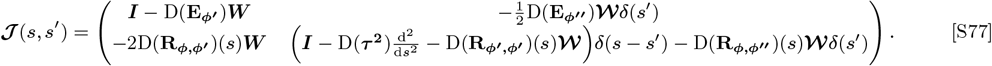

We let 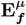 and 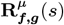 be the subcomponent of the population-flattened vectors corresponding to site *μ*. Similarly, we denote ***W*** ^*μθ*^ and ***𝒲*** ^*μθ*^ as the ν → *μ* sub-components of the population-flattened matrices of the mean and variance of the total weights. Then the ν → *μ* subcomponent of the rate moment Jacobian can be written as

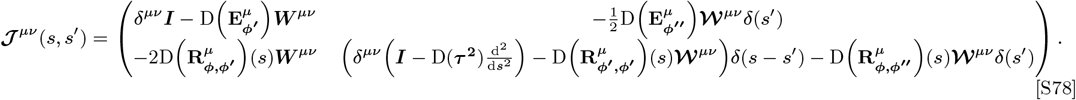

Inspired by Eq. (S71)The within-site (*μ* → *μ*) component of the From here onwards we will suppress the time-lag dependence of the rate moment Jacobian and its subcomponents for notational brevity. However, one must interpret quantities such as (***𝒥*** ^*μμ*^) ^− 1^ ***𝒥*** ^*μθ*^ as involving an implicit integral over the time lags, *i.e*. 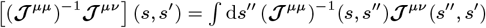, where the form of (***𝒥*** ^*μμ*^) ^− 1^ (*s, s*^′^) is similar to the product of the matrices in Eq. (S71).

The rate moment Jacobian expressed in terms of its sub-components is given by

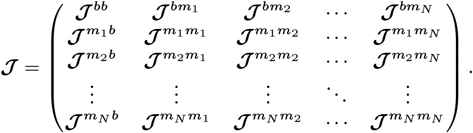

We now separate out cross-site projections from within-site couplings, defining

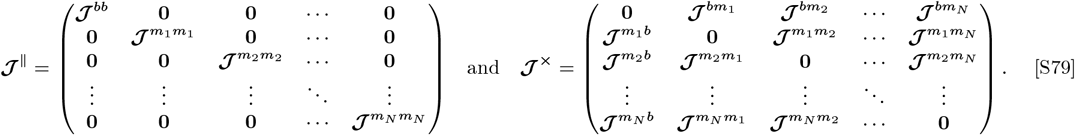

If the spectral radius of the operator (***𝒥*** ^‖^)^− 1^ ***𝒥*** ^***×***^ is less than one, then the inverse of the rate moment Jacobian can be expressed as 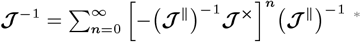

Since ***𝒥*** ^‖^ is a block-diagonal matrix, its inverse is a block-diagonal matrix in which the sub-blocks along the diagonal of (***𝒥*** ^‖^)^−^ are the inverses of the sub-blocks along the diagonal of ***𝒥*** ^‖^ This fact allows us to express the product − (***𝒥*** ^‖^)^− 1^ ***𝒥*** ^***×***^ as

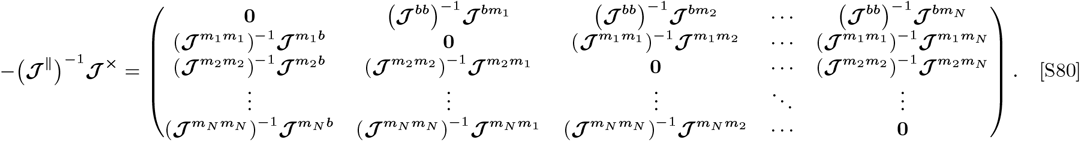

Defining the effective cross-site interaction matrices 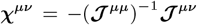 for *μ* ≠ ν, the inverse of the rate moment Jacobian can be succinctly written as

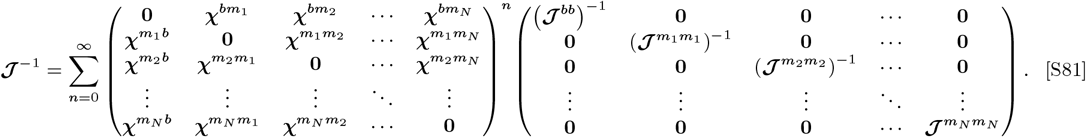

Finally, we denote 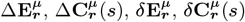 as the site *μ* sub-components of the corresponding population-flattened vectors.

Defining the decoupled responses, *i.e*. the linear responses in the absence of effective cross-site interactions, as ^†^

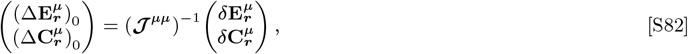

the rate moment responses can be expressed as

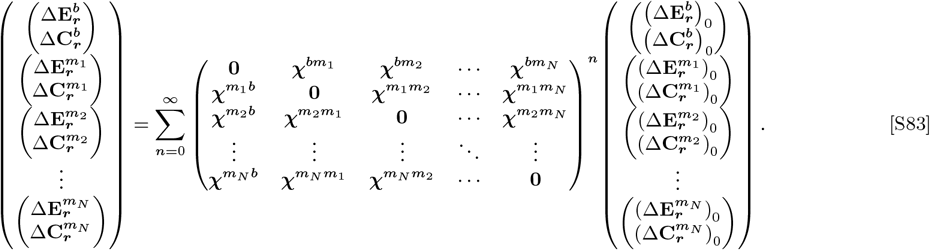

### I. Derivation of Effective cross-site Interactions

In this appendix, we elucidate the structure of the effective cross-site interaction 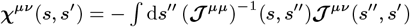 (Following the same steps as in App. F to compute (***𝒥*** ^*μμ*^)^− 1^(*s, s*^′^), we first let ***G***^*μμ*^(*s, s* ^*°*^) and ***δG***^*μμ*^(*s, s*^′^) be defined analogously to Eqs (S69) and (S70), but using the *μ* subcomponents of the population flattened vectors and matrices. Then ***χ***^*μθ*^ (*s, s*^′^) is given by

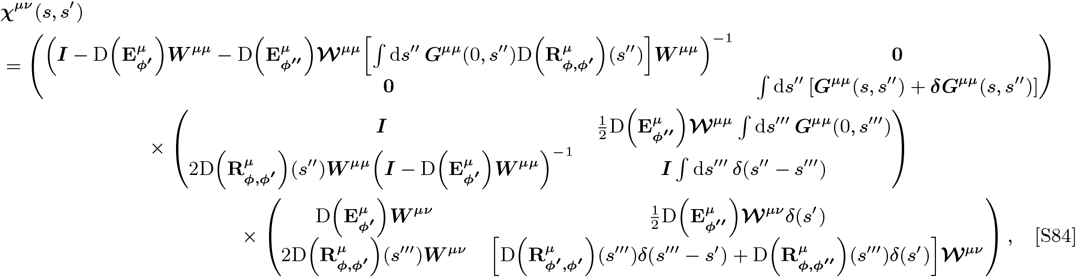

We define the effective coupling matrix as

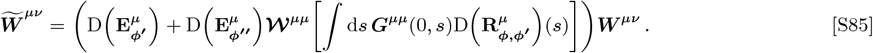

In the absence of disorder (𝒲^*μμ*^ = 0), the effective coupling matrix is simply the product of the mean gains times the mean weights. The effective coupling matrix is augmented by the presence of disorder, making it so that we cannot make general statements about the signs of the elements of this matrix. However, we find 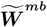 empirically obeys Dale’s law for all but one combination of network coupling and structure strengths considered in this study (Supp. Fig. S1c).

We further divide the effective cross-site interaction matrices into four sub-blocks, defined as

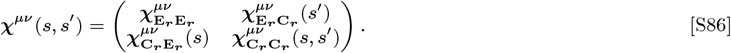

Using Eq. (S84), these four sub-blocks are then given by

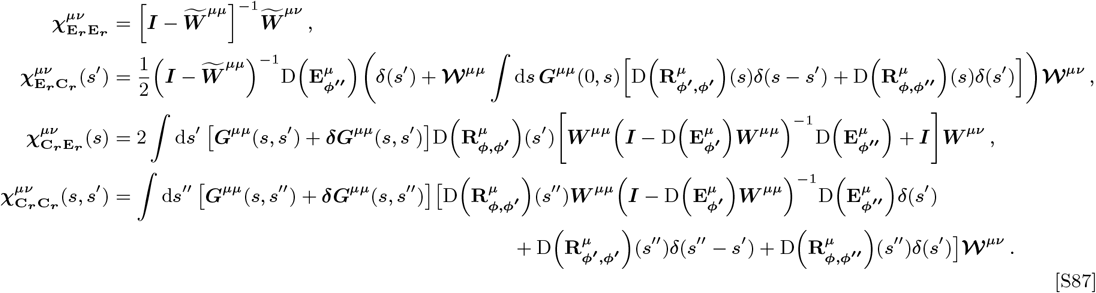

For additional notational brevity, we denote 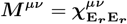, which becomes the only relevant term in the linear response in the mean-interaction approximation (*i.e*. if we approximate 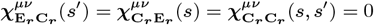). We can simplify the expression for 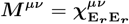 by using the fact that ***A*** ^− 1^ = det(***A***) adj (***A***) where adj (***A***) is the adjugate of ***A***. Similar to matrix inverses, the adjugate of a product of matrices obeys the identity adj (***AB***) = adj (***B***) adj (***A***). For 2 × 2 matrices specifically, adj (***I*** + ***A***) = ***I*** + adj (***A***). Finally, since our model assumes that the connection probability only depend on the presynaptic cell type, the mean weights can be expressed as 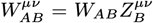 (see Eq. (S25) in App. C). Inspired by this factorization of the mean weights, we define a quantity 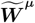 whose entries 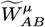 are given by

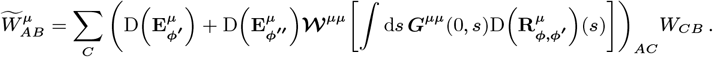

Thus, the effective coupling matrix can be factorized as 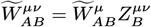, or expressed in matrix notation as 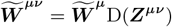.

Therefore, for an EI network, ***M*** ^*μθ*^ can be simplified as follows

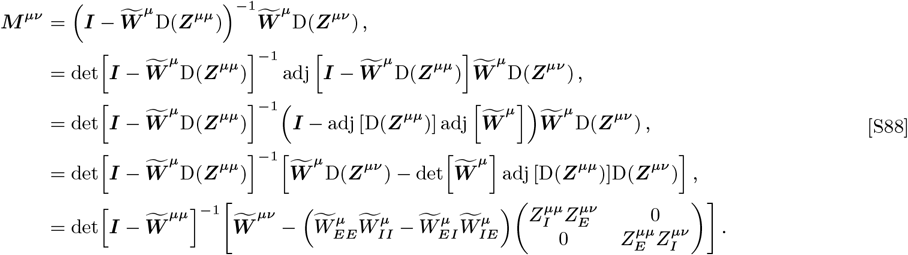

The last line of Eq. (S88) is equivalent to Eq. (7) of the Main Text. Intuitively, 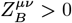 since site *μ* must receive a postitive contribution to its mean total weights from site ν (see Supp. Figs. S4 and S6). Therefore, the disynaptic term of ***M*** ^*μθ*^ is a diagonal matrix whose sign depends on det 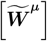. If 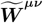 obeys Dales law, which 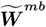 generally does (see Supp. Fig. S1c), then the condition det 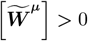 is equivalent to the condition that 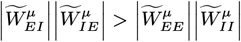. Thus, networks that satisfy this condition are “feedback-inhibition-dominated”.

In networks with more than two cell types generally adj (***I*** + ***A***) ≠ ***I*** + adj (***A***). However, adj (***I*** + ***A***) can be written as a sum of terms with up to *n* − 1 factors of the mean *W*_*AB*_. One can show that the term containing only *n* −1 factors of the weights is exactly equal to adj (***A***), meaning ***M*** ^*μθ*^ in a network with *n* cell types can be written as

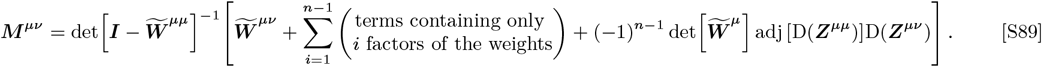

Thus, in the limit of strong coupling in which the final term dominates, the effective cross-site interaction matrix is approximately proportional to a diagonal matrix.

Finally, we remark about the stability of the decoupled network, which is an additional necessary condition required to be able to compute the linear response as an expansion series in powers of the interaction matrices. That is, even if Eq. (S83) converges, the decoupled response is undefined if the decoupled network is unstable. From Eq. (S30), the decoupled network is stable if the eigenvalue density 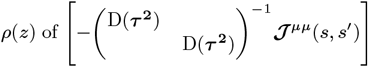 has nonzero support only in the region Re{*z*} < 0 for every site *μ*. This stability condition is equivalent to the condition that the eigenvalue density of 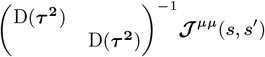 has nonzero support in the region Re *z* > 0. If we discretize time lags and handle this operator numerically, then we can define the trace and determinant of this operator in terms of the trace and determinant of the time-lag-discretized matrix. These are then given by

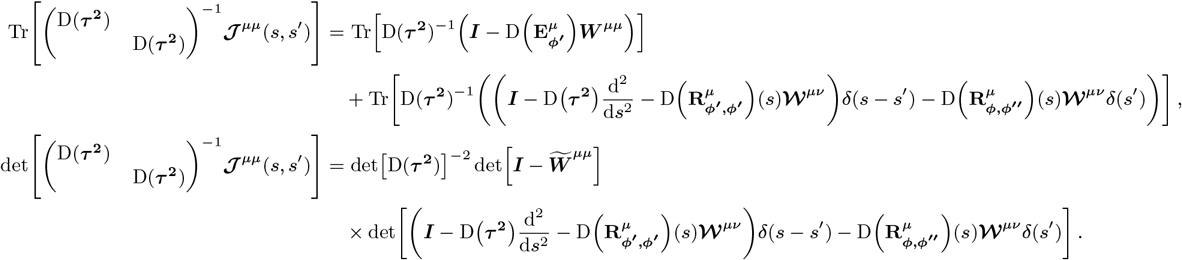

Suppose that the discretized operator 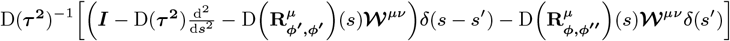 has only eigenvalues with positive real part. Then the conditions for the decoupled network to be stable are

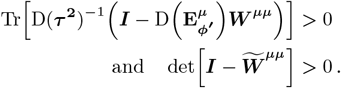

We can arrive at the effective interaction matrix in the mean-interaction approximation by simply letting 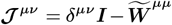.

In this case, the stability conditions for the decoupled matrix are

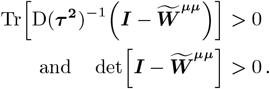

Therefore, it could be said that the mean-interaction approximation is only appropriate if these two conditions are met. Therefore, when the mean-interaction approximation is valid, it should be the case that det 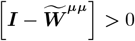 as we discussed in the Main Text.

### J. Generalization of Methods to Higher Dimensional Feature Spaces

Our analysis readily generalizes to *d*-dimensional toroidal features spaces where the feature dependence of the rate moments are well approximated as a baseline plus a mixture of products of wrapped Gaussian functions. Denoting toroidal locations, Gaussian widths, the number of discrete points along each edge, and the extent of each dimension as *d*-dimensional vectors ***θ, s, N*** _***θ***_, and ***L***, respectively, we will define the multidimensional wrapped Gaussian as 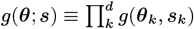 and the normalized baseline-subtracted wrapped Gaussian as 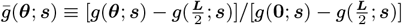. The connection probabilities will be parameterized as

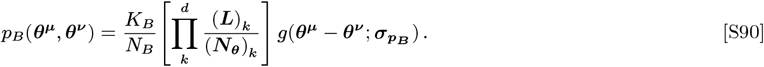

Defining the elements of the overlap matrix as

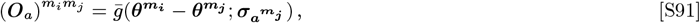

then the feature dependence of the rate moments will be approximated as

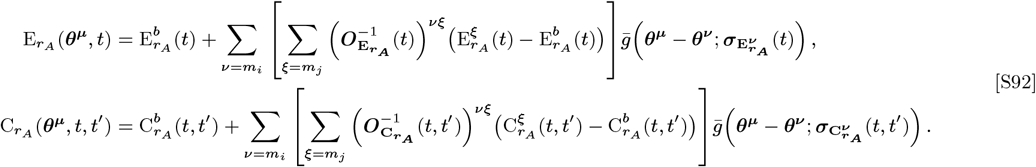

In the *d*-dimensional space, convolving the baseline-plus-Gaussian rate moments by the Gaussian weight profiles results in a modification of the function ζ. Letting 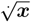 and ***x***^2^ denote the element-wise square-root and squaring of a vector, the generalization to ζ is given by

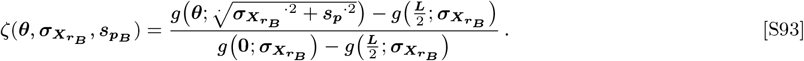

There are now *N*(*d* + 1) +1 degrees of freedom in each of the rate moments, parameterized by the baseline, the values at the *N* visual-stimulus-matched sites, and the *d* Gaussian widths per visual stimulus. These parameters can be specified by *N*(*d* + 1) sites corresponding to the *N* visual stimulus features and *Nd* auxiliary locations, plus the theoretical baseline.

## Footnotes

* For completeness, if we reintroduce the time-lag-dependence back into the Jacobian matrices then the pertrubative series will be of the form 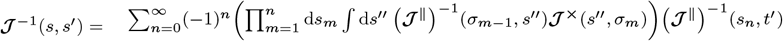 where *s*_0_ = *s*.

† If we reintroduce the implicit time integrals then the definition of the decoupled susceptibilities is 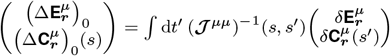

